# Widespread position-dependent transcriptional regulatory sequences in plants

**DOI:** 10.1101/2023.09.15.557872

**Authors:** Yoav Voichek, Gabriela Hristova, Almudena Mollá-Morales, Detlef Weigel, Magnus Nordborg

## Abstract

Much of what we know about eukaryotic transcription stems from animals and yeast, however, plants have evolved separately for 1.6 billion years, leaving ample time for divergence in transcriptional regulation. Here, we set out to elucidate fundamental properties of cis-regulatory sequences in plants. Using massively parallel reporter assays across four plant species, we demonstrate the central role of sequences downstream of the transcription start site (TSS) in transcriptional regulation. Unlike animal enhancers that are position-independent, plant regulatory elements depend on their position, as altering their location relative to the TSS significantly affects transcription. We highlight the importance of the region downstream of the TSS in regulating transcription by identifying a DNA motif that is conserved across vascular plants and is sufficient to enhance gene expression in a dose-dependent manner. The identification of a large number of position-dependent enhancers points to fundamental differences in gene regulation between plants and animals.

## Introduction

Eukaryotes have diverged for ∼1.6 billion years^1^. Despite their immense diversity, our basic knowledge of eukaryotic transcription is mainly based on observations in yeast and a few animals. The knowledge gap is even more striking when we consider transcriptional regulation in the context of multicellularity, which requires regulatory mechanisms to enable cell-type-specific gene expression. Complex multicellularity arose independently at least six times, including once in animals and once in plants^2,3^. These independent inventions require specialized mechanisms of transcriptional regulation to allow each cell type to express a different set of genes. The mechanisms that support this complexity likely evolved with the emergence of multicellularity^4^, and we already know that plants and animals solved many of the challenges associated with multicellularity very differently. In particular, there is no reason to believe that what is true for animals is also true for plants when it comes to principles that go beyond the basic transcriptional machinery^5,6^. Yet, how regulatory sequences function is widely assumed to be similar to animals and yeast^7^. Here we show that the basic property of the majority of animal enhancers - position-independence - does not hold for plants.

## Results

We first set out to determine the typical locations of regulatory regions near genes in Arabidopsis by large-scale mapping of variants underlying expression quantitative trait loci (eQTL). Therefore, we analyzed genotypic and rosette-transcriptomic data from the Arabidopsis 1,001 Genomes Project to identify cis-eQTL within 10 Kb of each gene^8,9^. While we expected to find most eQTL in proximal promoters upstream of the transcription start site (TSS), we discovered a similar proportion of eQTL downstream of the TSS (Fig. 1A). As eQTL are more likely to occur where the density of single-nucleotide polymorphisms (SNPs) is higher, the lower sequence diversity downstream of TSSs made the downstream eQTL enrichment even more unexpected (Fig. 1A). This pattern was consistent across multiple gene expression datasets, and accounting for linkage between SNPs only intensified it (Fig. S1). Our eQTL analysis pointed to potential central regulatory regions downstream of the TSS in Arabidopsis being wide spread.

**Figure 1:**
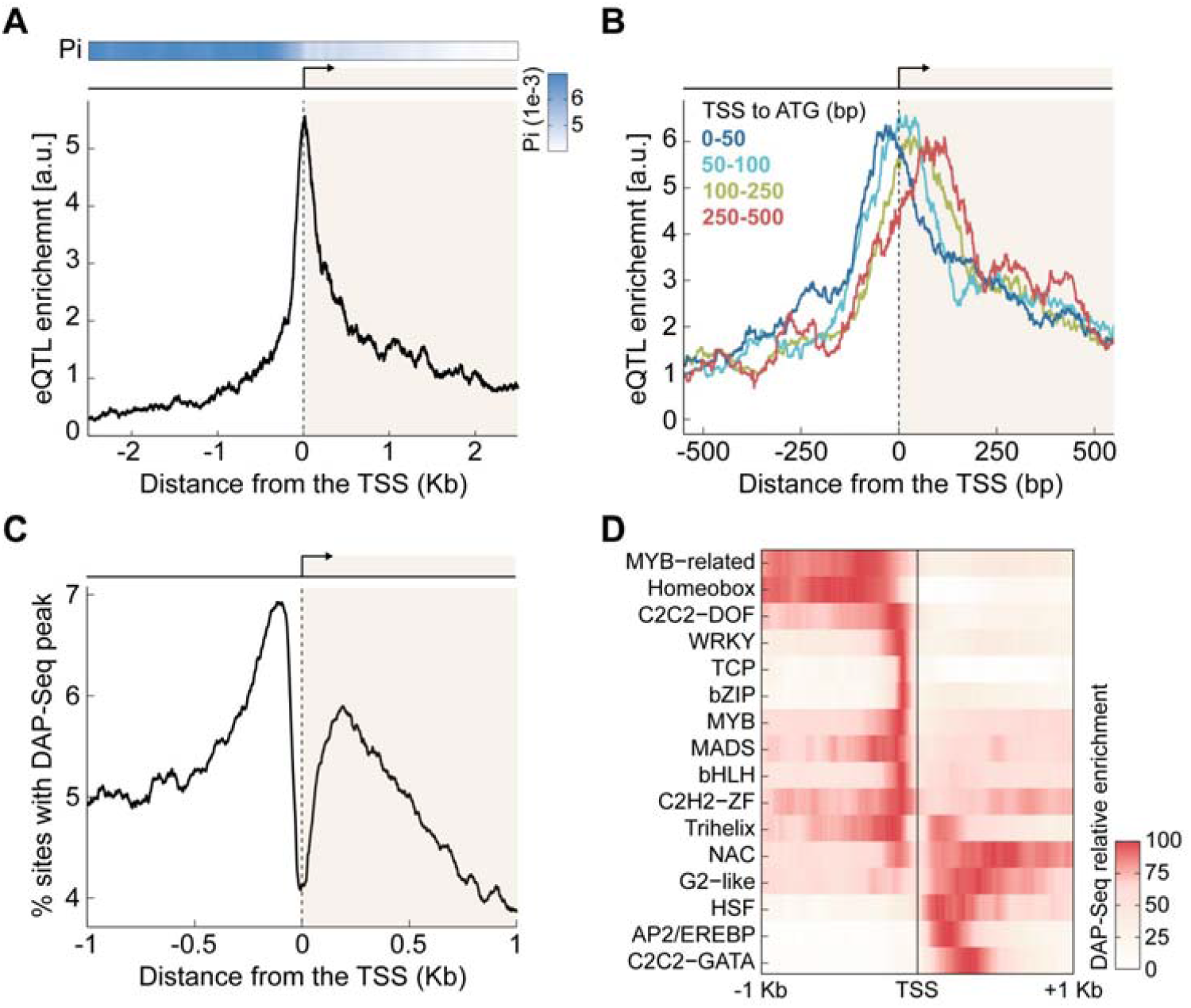
Evidence for transcription-regulatory sequences downstream of the TSS. **(A)** eQTL enrichment (below) and nucleotide diversity (Pi, above) near TSSs. **(B)** eQTL enrichment for genes with different TSS-to-ATG distances. Group counts: 1,088 (0-50), 968 (50-100), 1,122 (100-250), and 577 (250-500). **(C)** Proportion of sites with a DAP-Seq^13^ peak center, based on data for 529 TFs. **(D)** DAP-Seq peak enrichment, as in **C**, consolidated for each TF family with at least 10 members in the dataset^13^; maximum signal for each TF family was scaled to 100. In **A-C**, data smoothed using a 100 bp rolling window. a.u., arbitrary units.

We next examined possible explanations for the observed eQTL distribution. If causal eQTL variants within transcripts are in sequences controlling mRNA stability, these should be more frequent in exons than introns, which are removed by splicing. While this is what is seen for human eQTL^10^, we observed no such preference for exons in Arabidopsis (Fig. S2A-E). eQTL were also not enriched toward the end of transcripts (Fig. S3), even though 3’ untranslated regions (3’ UTRs) have known roles in controlling mRNA stability^11,12^. Finally, we asked whether eQTL were more likely to occur outside coding regions, which have strong sequence constraints. Indeed, eQTL tended to be most frequent just downstream of the TSS for genes with longer TSS-to-ATG distances (including 5’ UTRs and introns, Fig. S2F) and just upstream of the TSS for genes with shorter TSS-to-ATG distances (Fig. 1B & S2G). These findings suggest that downstream regulatory regions are enriched between the TSS and the start codon and affect transcription rather than mRNA stability.

If the location of eQTL variants in the proximity of genes results from variation in transcription rate – as opposed to mRNA stability – then chromatin and transcription factors (TFs) are likely to be involved. The first observation in agreement with this was that histones H3.1 and H3.3 enrichment downstream of the TSS moves away from the TSS with increasing TSS-to-ATG distances (Fig. S4). Second, binding sites of 529 TFs, as measured by DNA affinity purification sequencing (DAP-Seq)^13^, have two prominent peaks, upstream as well as downstream of the TSS (Fig. 1C). Individual TFs have a preference for binding on only one side of the TSS, with similar preferences for members of the same TF family (Fig. 1D & S5). In-vivo data of three TFs binding confirmed the preference of TFs to bind on either side of the TSS (Fig. S6). We do not think that inaccuracies in the TSS annotations greatly affect our results, given that the clear dip in TF binding sites is centered on annotated TSSs. These analyses support that many genes have a transcription-regulatory region downstream of the TSS.

To systematically investigate the role of sequences downstream of the TSS in controlling gene expression, we designed a massively parallel reporter assay^14^ (MPRA, Fig. 2A). We synthesized 12,000 160-long bp fragments, derived from regions 40-200 bp upstream or 40-360 bp downstream of the TSSs of highly expressed Arabidopsis genes, excluding 80 bp around the TSS (-40 bp to 40 bp), which contains the core promoter. Downstream-derived fragments included exons and introns, except for donor and acceptor splicing sites. We inserted these fragments in their original orientation on either side of the TSS of a GFP reporter gene. Insertion-free constructs served as controls. The downstream insertion site was located in an intron of the reporter gene, to rule out effects due to altered mRNA sequence on mRNA stability. For robust quantification, multiple variants were generated for each insertion, with a 15 bp random barcode within the transcript. Barcodes and tested regulatory fragments were linked by DNA sequencing, and transcriptional activity was read out by RNA sequencing.

**Figure 2:**
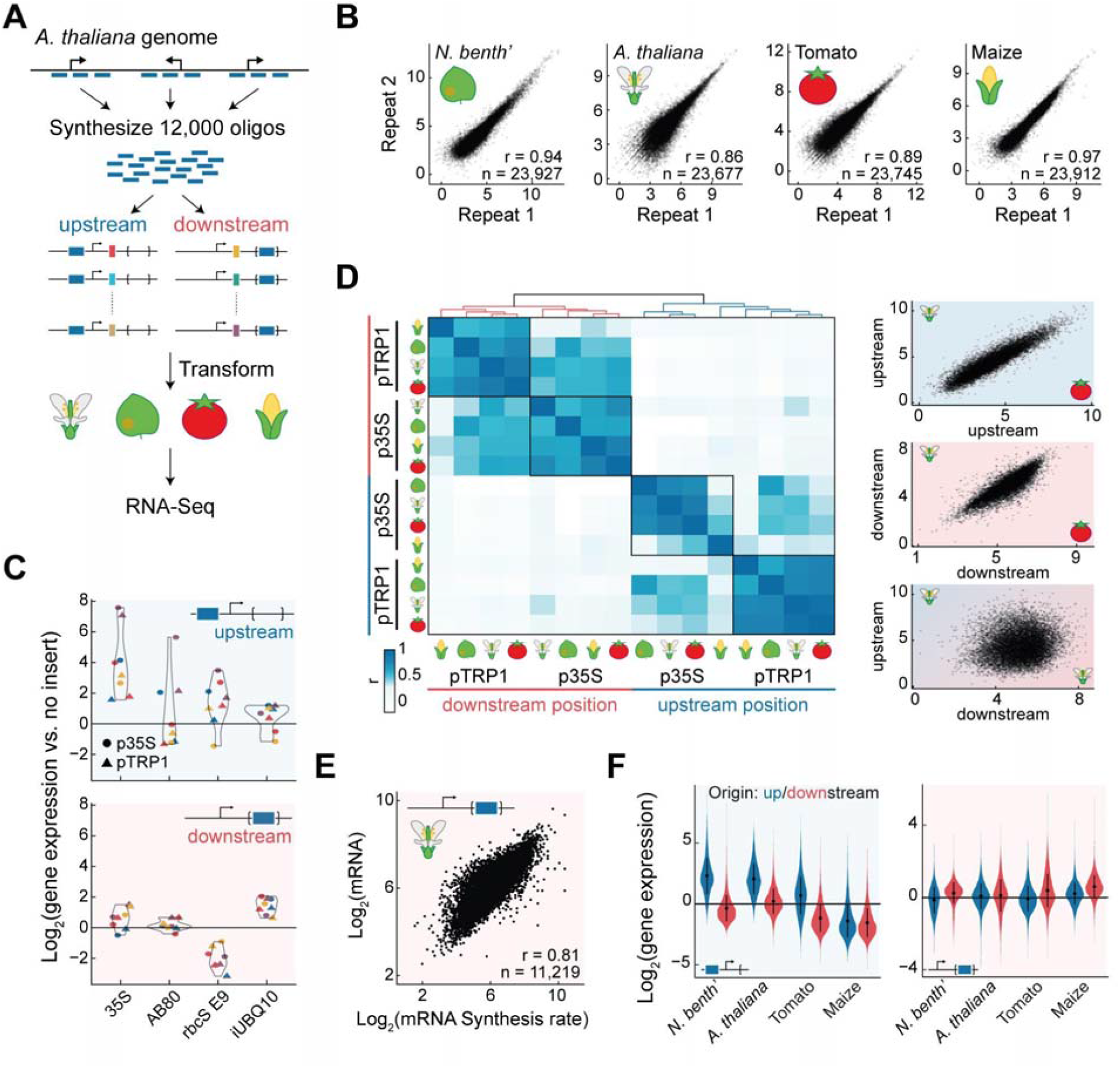
Position-dependent enhancers reside inside transcribed regions. **(A)** MPRA overview: 12,000 fragments (160 bp), originating from upstream or downstream of the TSS of Arabidopsis genes, were synthesized, pooled, and inserted upstream of the TSS or within the intron of a reporter gene, and tagged by barcodes. Following transient transformation into one of four species, barcoded RNA sequencing quantified expression. **(B)** High reproducibility in MPRA experiment replicates, demonstrated here for CaMV 35S-minimal promoter-based libraries, plotted as log_2_(gene expression). Pearson’s correlation coefficients *r* and number of fragments *n* are indicated. **(C)** Comparison of expression of constructs with control enhancer fragments and no-insert constructs, for upstream (top) and downstream (bottom) insertions. Depicted for *N. benthamiana* (purple), *A. thaliana* (blue), tomato (red), and maize (yellow); construct backgrounds are coded by symbol shape. **(D)** Left, Pearson’s correlation coefficients across all libraries, hierarchically clustered. Construct background and insertion position are indicated. Right, activity of pTRP1-based constructs, as log_2_(gene expression), compared between upstream (top, r=0.92) and downstream (middle, r=0.85) libraries of Arabidopsis and tomato, and between upstream and downstream libraries of Arabidopsis (bottom, r=0.13). Correlations include 11,736±241 fragments. **(E)** Comparison of mRNA steady-state levels and synthesis rates in downstream MPRA with pTRP1 constructs. Correlation and fragment counts as indicated in **B. (F)** Comparison of activity of upstream-(blue, 3,979±5 fragments) and downstream-derived (red, 7,941±3 fragments) fragments relative to no-insertion constructs, when fragments are positioned upstream (left) or downstream (right) of the TSS. Constructs are p35S-based. Error bars depict the mean ±1 standard deviation.

We used two different GFP reporter constructs to provide different promoter contexts (Fig. S7): the 46 bp CaMV 35S minimal promoter, commonly used to test plant enhancers, in combination with a short synthetic 5’ UTR, and a 700 bp Arabidopsis *TRP1* promoter fragment including its 5’ UTR, previously used to study the effect of introns on gene expression^15^. In order to derive conclusions with broad applicability to flowering plants, we quantified activity of the libraries in four different species - in Arabidopsis, tomato, and maize using transfection of leaf protoplasts, and in *N. benthamiana* using leaf infiltration of *Agrobacterium tumefaciens* into mesophyll cells. We reasoned that the use of two different transformation methods would increase the robustness of our conclusions. Reproducibility was ensured through three to four replicated experiments (Fig. 2B and S8).

As a control, a small fraction of the synthesized fragments were from known enhancers, previously examined in MPRA^14^. In most cases, these fragments increased expression when placed upstream of the TSS (Fig. 2C, S9A), but not when placed inside introns, in agreement with previous results^14^. Segments of the *UBQ10* intron, known to enhance expression^16^, were also included in the library. These intron-derived fragments drove higher expression when inserted into the intron of the reporter gene rather than when inserted upstream of the TSS (Fig. 2C).

The position-dependent effects observed for the known enhancers seem to be representative for the majority of tested fragments. We found that fragments had similar activity independently of species, promoter, or how they were introduced into the host cell (Fig. 2D, S9B-C). In contrast, the relative activity greatly changed when the same fragment was inserted either upstream or downstream of the TSS, even when using the same backbone and species. We were surprised by this lack of correlation; one possibility is that the downstream insertion mainly affects mRNA stability. To test this, we modified our MPRA setup to measure mRNA synthesis directly (Fig. S10A). We transformed Arabidopsis leaf protoplasts with pTRP1-based constructs containing the fragment library downstream of the TSS and added 5-ethynyl uridine (5-EU) 20 min before RNA harvest, followed by purification of 5-EU-containing mRNAs^17^. Sequencing of these newly synthesized mRNAs revealed that the main effect of inserting fragments downstream is on transcription rate (Fig. 2E, S10B). These results suggest that, unlike the position-independence seen for animal enhancers, the activity of flowering plant enhancers is strongly dependent on their position relative to the TSS.

The original genomic location of the fragment played a substantial role as well. Generally, fragments increased expression when positioned in their original position relative to the TSS, but the extent varied between backbone and species (Fig. 2F and S11). Enhancers were relatively more effective in the CaMV 35S promoter than in the *TRP1* promoter construct, in which fragments insertions disrupted the *TRP1* genomic sequence (Fig. S11). In maize, fragments often reduced reporter expression when inserted upstream, regardless of genomic origin, in agreement with previous observations^18^. This finding, along with the strong correlation among the relative activity of fragments across all libraries, suggests that while absolute levels are strongly influenced by backbone and species, the relative effects of different fragments in the same position are similar across species.

An immediate question that arises from our observation is how sequences downstream of the TSS control transcription activity. Given the results in Fig. 1D, we suspected that TFs promote transcription in this region. Although TFs often work in concert, we hypothesized that even the DNA-binding motifs of single TFs will be more abundant in strong downstream enhancers. Thus, we searched for 6-bp sequences (6-mers) whose presence downstream of the TSS was associated with increased or decreased expression (Fig. 3A-B & S12). We found more 6-mers that promoted expression than 6-mers that repressed expression when found downstream. These 6-mers are thus potentially part of sequence motifs bound by TFs downstream to the TSS.

**Figure 3:**
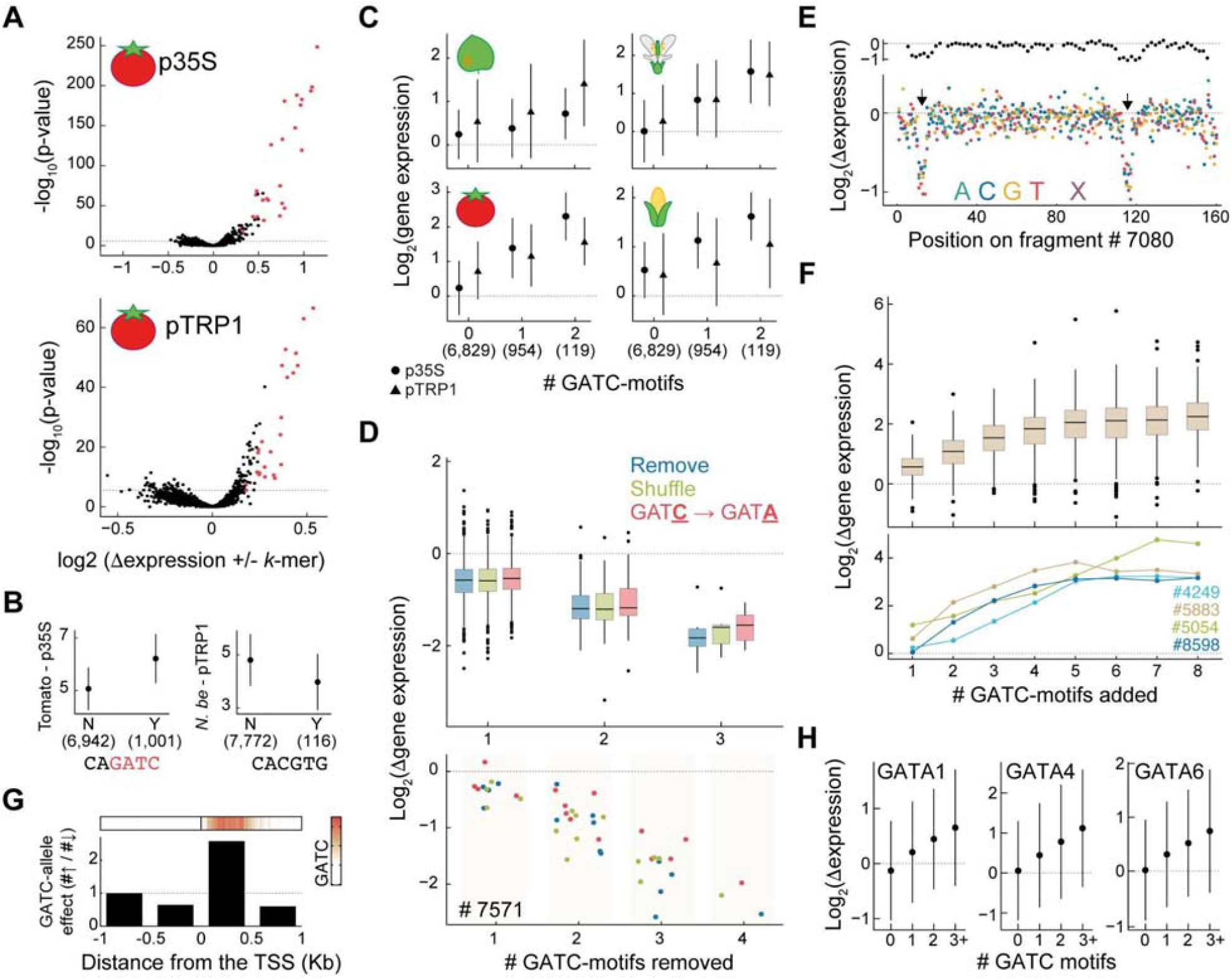
The GATC motif is sufficient to increase expression when positioned downstream of the TSS. **(A)** Association of 6-mers with downstream-MPRA activity. Each 6-mer’s presence (or absence) in downstream-derived fragments was compared based on the difference in average log_2_(gene expression) (x-axis) and Mann-Whitney U test p-value (y-axis). Displayed for p35-(top) and pTRP1-based (bottom) constructs in tomato. Red points denote 6-mers with G-A-T-C; dashed line marks the 5% Bonferroni multiple testing threshold. **(B)** Activity distribution for downstream-derived fragments with (Y) or without (N) 6-mers when inserted downstream. Plotted as log_2_(gene expression) in the MPRA for CAGATC in tomato with p35S-based constructs (left), and for CACGTG (G-box^19^) in *N. benthamiana* with pTRP1-based constructs (right). **(C)** Relative activity of downstream fragments when inserted downstream, as a function of the number of YVGATCBR consensus motifs in the tested fragments. Group sizes: 6,855 (no motif), 956 (1 motif), and 119 (2 motifs). Backbone and species are indicated. **(D-F)** Differences in activity of the p35S-downstream library between original and mutated fragments. **(D)** Effect of removing GATC motifs on activity of 823 fragments containing one (top), and a specific example with four motifs (bottom). Motifs were removed by deletion, 8 bp shuffle, or GATC-to-GATA mutation. **(E)** Deep-mutagenesis of fragment #7080: effects of 10 bp deletions (top), 1 bp deletions (X), or 1 bp mutations (bottom). Arrows highlight GATC motifs. **(F)** Effect of adding GATC-motifs for 221 fragments with incremental motif additions (top), and four specific examples (bottom). **(G)** GATC-motif-gain/loss alleles in the 1,001 Genomes population of accessions linked to nearby gene expression. The bar-graph showcases the ratio of number of significant associations with higher vs. lower expression in the GATC-motif allele, grouped by distance to the TSS (bottom). Top, GATC motif’s enrichment in proximity to the TSS. **(H)** Gene expression response to overexpression of GATA TFs (see subfigures) versus GFP, plotted for genes with 0, 1, 2, or ≥3 GATC motifs within 500 bp downstream of the TSS. The groups contain 11,885±420, 4,174±124, 973±27, and 350±7 genes, respectively. Overexpression of GATA TFs or GFP was driven by double 35S promoters in Arabidopsis protoplasts, which were harvested 8 hours after transformation. Error bars in **B-C** and **H** represent mean ± 1 standard deviation. Boxplots in **D** and **F** display median, IQR, and 1.5x IQR, with outliers as points. The numbers in parentheses in **B-C** indicate the number of fragments.

Across species and backbones, 6-mers including a GATC sequence had the strongest effect (Fig. 3A, S12-13). To quantify the GATC effect, we combined the six 6-mers with the strongest effect into a 8-bp YV**GATC**BR motif (Y=CT, V=ACG, B=CGT, R=AG, Fig. S14), referred to as the “GATC motif”. Transcriptional activity increased with the number of GATC motifs in the fragment, with each copy associated with an average increase in expression of nearly 50% (Fig. 3C, S10B). The effect was minimal with fragments inserted upstream of the TSS (Fig. S15).

To investigate the effects of the GATC motif further, we synthesized 18,000 additional oligonucleotides, each a variant of a fragment from our initial pool as described below. These fragments were inserted downstream of the TSS in both backbones, and their effect on gene expression were measured in Arabidopsis protoplasts across three replicates (Fig. S16). First, we tested the requirement of the motif by focusing on 841 downstream-derived fragments containing a GATC motif. By deleting, shuffling, or modifying the core GATC to GATA, we effectively removed these motifs. We found that such removal led to an average 50% decrease in gene expression, regardless of mutation type (Fig. 3D, S17A).

To supplement the GATC-focused mutation analysis, we conducted a deep mutational scan of 13 downstream-derived fragments. For each, we (i) deleted every set of 10 consecutive base pairs, and (ii) either mutated each nucleotide to its three alternatives or deleted it. This resulted in 736 derivatives from each original fragment. Any change to the core 4 bp of the GATC motif decreased activity, underscoring the motif’s strict constraints (Fig. 3E, S18-19). As expected, these analyses also revealed additional sequences that do not include GATC motifs as important for enhancing activity of the tested fragments (Fig. S20).

We next explored the sufficiency of GATC motifs for enhancing gene expression. We started with a random set of 221 fragments from our initial set (166 downstream- and 55 upstream-derived) and incrementally added 1 to 8 GATC motifs to the fragments. Expression consistently increased with each added copy, even for upstream-derived fragments (Fig. 3F, S17B-C). Remarkably, 97% of these fragments enhanced expression as soon as at least 4 GATC motifs were added. The enhancement was a function of the basal activity of each fragment, with the increased activity of highly active fragments becoming saturated after a single addition, and the activity of the initially least active fragments remaining unsaturated even after adding eight GATC motifs (Fig. S17D). This finding suggests that the GATC motif and other activity-enhancing sequences may act by the same mechanism to increase expression of the reporter constructs.

Finally, to confirm the inferences from our synthetic MPRA mutational analysis of the GATC motif, we explored the effects of natural variation in the GATC motif by returning to the Arabidopsis 1001 Genomes Project data^8,9^. We identified gains and losses of GATC motifs near TSSs, and asked how these correlated with expression of the affected genes. We categorized significant associations based on whether the allele with the GATC motif had higher or lower expression. Consistent with our MPRA findings, an enrichment of higher expression was observed exclusively in the GATC-motif allele situated downstream of the TSS, particularly within the initial 500 bp (Fig. 3G, S21). Intriguingly, this is also where the GATC motif is predominantly found (Fig. 3G), reinforcing its role in enhancing gene expression when located downstream of the TSS.

What might be the mechanisms underlying the observed effects of GATC motifs? In plants, the GATC motif is recognized by GATA TFs^13^ (see supplementary note S1, Fig. S22), which are linked to diverse biological functions^20^. The Arabidopsis genome encodes 30 of these TFs. Available DAP-Seq data^13^ reveal GATA factor binding enrichment within 500 bp downstream of the TSS (Fig. 1D). In this region, regardless of genomic context, 7,397 genes have at least one GATC motif (Fig. S23A-B). Transient overexpression of three different GATA TFs in Arabidopsis leaf protoplasts confirmed that these genes are direct targets of GATA TFs (Fig. 3H, S24-25). Gene ontology analysis shows these genes to be enriched in processes related to the Golgi apparatus, endoplasmic reticulum, endosomes, and vesicle-mediated transport (table S6). Given its prevalence, association with the secretion system, and the evidence for conservation between species (Fig. S26), the GATC motif likely acts as a widespread and conserved regulatory signal in diverse biological functions.

Enhancer sequences typically consist of multiple DNA motifs that are targeted by specific TFs, of which Arabidopsis has more than 1,500^21^. Given this diversity, individual regulatory motifs have generally limited power to predict absolute levels of gene expression. We found nevertheless a strong positive relationship between the occurrence of the GATC motif within 500 bp downstream of the TSS and gene expression (Fig. 4A). This relationship was driven by motifs in all genomic contexts, with motifs in introns and UTRs showing a stronger association (Fig. S27). Analysis of all 6-mer counts, both downstream and upstream of the TSS, showed that the GATC motif has the strongest association with gene expression (Fig. 4B). This identifies the downstream GATC motif as a singularly potent regulatory sequence.

**Figure 4:**
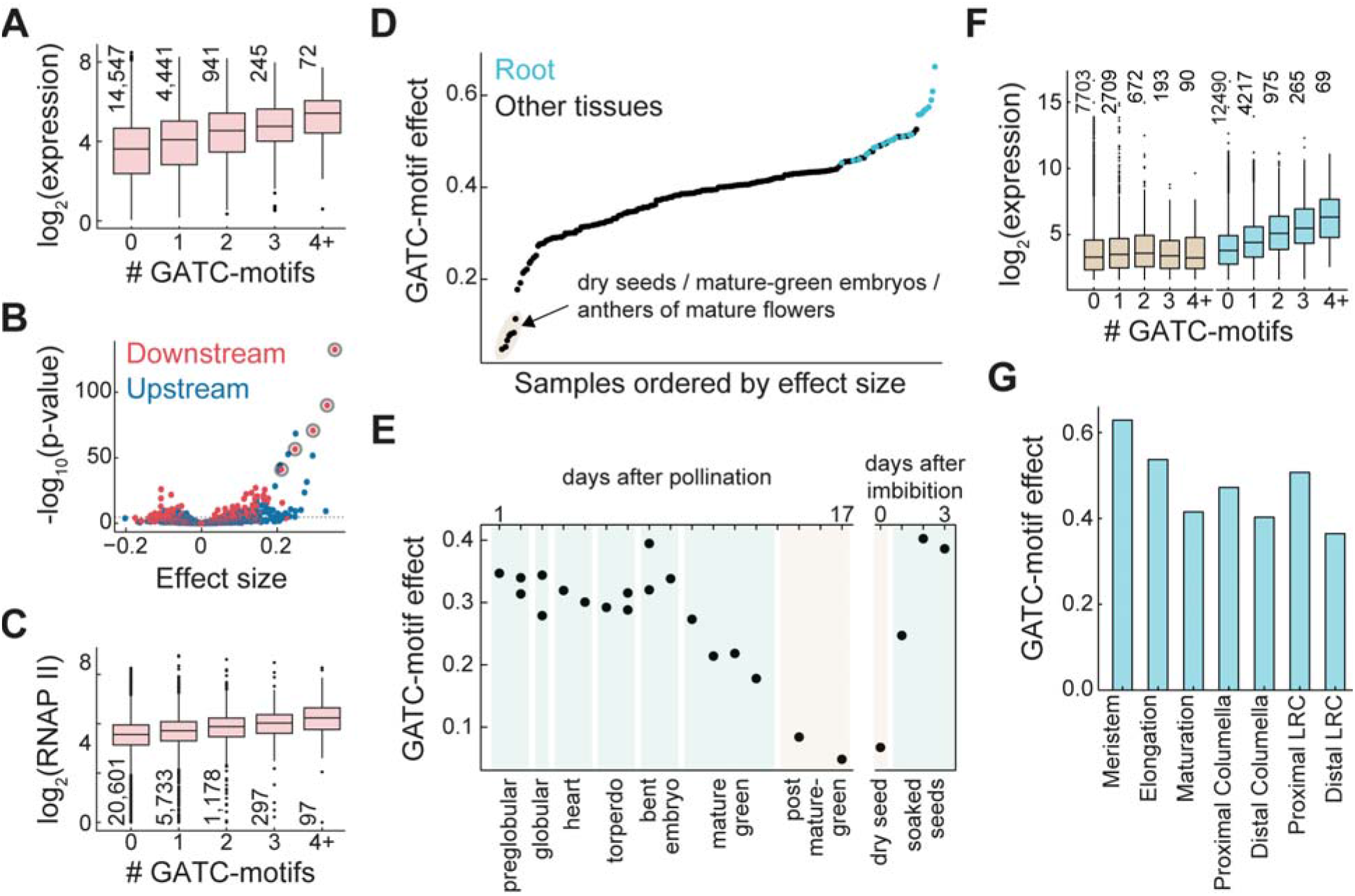
Cell-type specificity of the GATC-motif effect on gene expression. **(A)** In aerial parts of Arabidopsis seedlings^28^, gene expression correlates with the number of GATC motifs within 500 bp downstream of the TSS. Expression values are depicted for various motif counts, with “4+” representing 4-9 motifs. A linear fit reveals a GATC-motif effect size of 0.4 (p-value: 5*10^−128^), indicating the average expression increase for each added motif. **(B)** For all 6-mers within 500 bp up- or downstream of the TSS, effect size and p-value are determined as in A. The five most significant downstream 6-mers, all containing the G-A-T-C sequence, are highlighted with circles. A 5% Bonferroni threshold indicated by a dashed line. **(C)** Average RNA polymerase II occupancy at genes plotted as in **A**, with an effect size of 0.17 (p-value: 10^−107^). **(D)** GATC-motif effect sizes from a compendium of 200 tissue-specific gene expression data sets^25–27^, as determined in **A**. Samples with the lowest effect sizes are shaded and detailed. **(E)** Chart of GATC-effect size during embryo/seed development and upon imbibition^26,27,29^. **(F)** Expression values in dry seeds (brown) and seedling roots^26^ (blue) plotted as in **A**, with effect sizes of 0.07 (p-value: 2.6*10^−3^) and 0.57 (p-value: 4*10^−400^), respectively. **(G)** GATC-effect sizes across different root developmental stages, averaged from single-cell expression data^30^. Boxplots in **A, C**, and **F** display median, IQR, and 1.5x IQR with outliers as points, number of genes per category is also indicated.

As the effects of the GATC-motif are strong enough to be observed in genome-wide measurements, we can investigate its function using the many other resources available for Arabidopsis. As one example, if the GATC motif indeed works primarily through transcription and not mRNA stability, we expect it to affect chromatin measurements. Indeed, occurrence of GATC-motifs is correlated with the active marks H3K4me3 and H3K36me3^22^, as well as RNA polymerase II occupancy^23^ (Fig. 4C, S28A-B). Moreover, we observed a correlation with genome-wide measurements of mRNA synthesis but not mRNA half-life^24^ (Fig. S28C-F). These results further support an effect through transcription, in accordance with our mRNA synthesis measurements in the MPRA (Fig. S10B).

Our MPRA inferences came only from enhancer activity in leaf cells, so we were curious whether the GATC motif was also effective in other tissues. Analyzing a compendium of gene expression in different tissues and developmental stages verified once more the potent activity of the GATC motif in increasing expression, yet also revealed a roughly 3-fold fluctuation in the impact of the GATC motif^25–27^ (Fig. 4D). Its influence was smallest in specific seed developmental stages: decreasing from mature green embryo stages through seed drying, then rebounding upon germination^27^ (Fig. 4E-F).

Conversely, the strongest effects were seen in roots (Fig. 4D). Single-cell expression data from Arabidopsis roots^30^ pinpointed the meristem as the region most associated with the GATC-motif, with decreasing effects through the elongation and maturation zones (Fig. 4G). This trend held true across various root cell types (Fig. S29). Similarly, in the vegetative shoot apex^31^, the GATC motif’s impact diverged between cell types – for example, mesophyll cells showing muted effects compared to the pronounced effects in epidermal cells (Fig. S30). Overall, the GATC motif’s regulatory role spans the entire body plan of the plant, being modified by tissue and cell type. In addition, the expression of GATA TFs, especially from subfamily A^20,32^, correlates with the effect of the GATC motif (Fig. S31). This suggests that the GATC motif functions like a general rheostat, modulating gene expression of thousands of genes across plant cell types, likely through GATA TFs.

To evaluate the conservation of the GATC-motif’s influence on gene expression, we correlated the number of GATC motifs in the 500 bp downstream region with gene expression across various land plants^26,33–40^. Consistent with our MPRA findings in four flowering plants, GATC-motif count correlated with gene expression in all flowering plants examined (Fig. 5, S32). This conservation extended to the gymnosperm *Pinus tabuliformis* and the fern *Ceratopteris richardii*, albeit with reduced effect-size compared to flowering plants. In the lycophyte *Selaginella moellendorffii* the association of the GATC-motif with gene expression was markedly weaker, though still significant. Among bryophytes, there was a modest effect in *Marchantia polymorpha*, with relatively very weak statistical support, and no clear effect in *Physcomitrium patens*. Overall, the impact of the GATC-motif, and by extension of downstream regulatory sequences, is conserved in vascular plants, with a weaker influence outside flowering plants.

**Figure 5:**
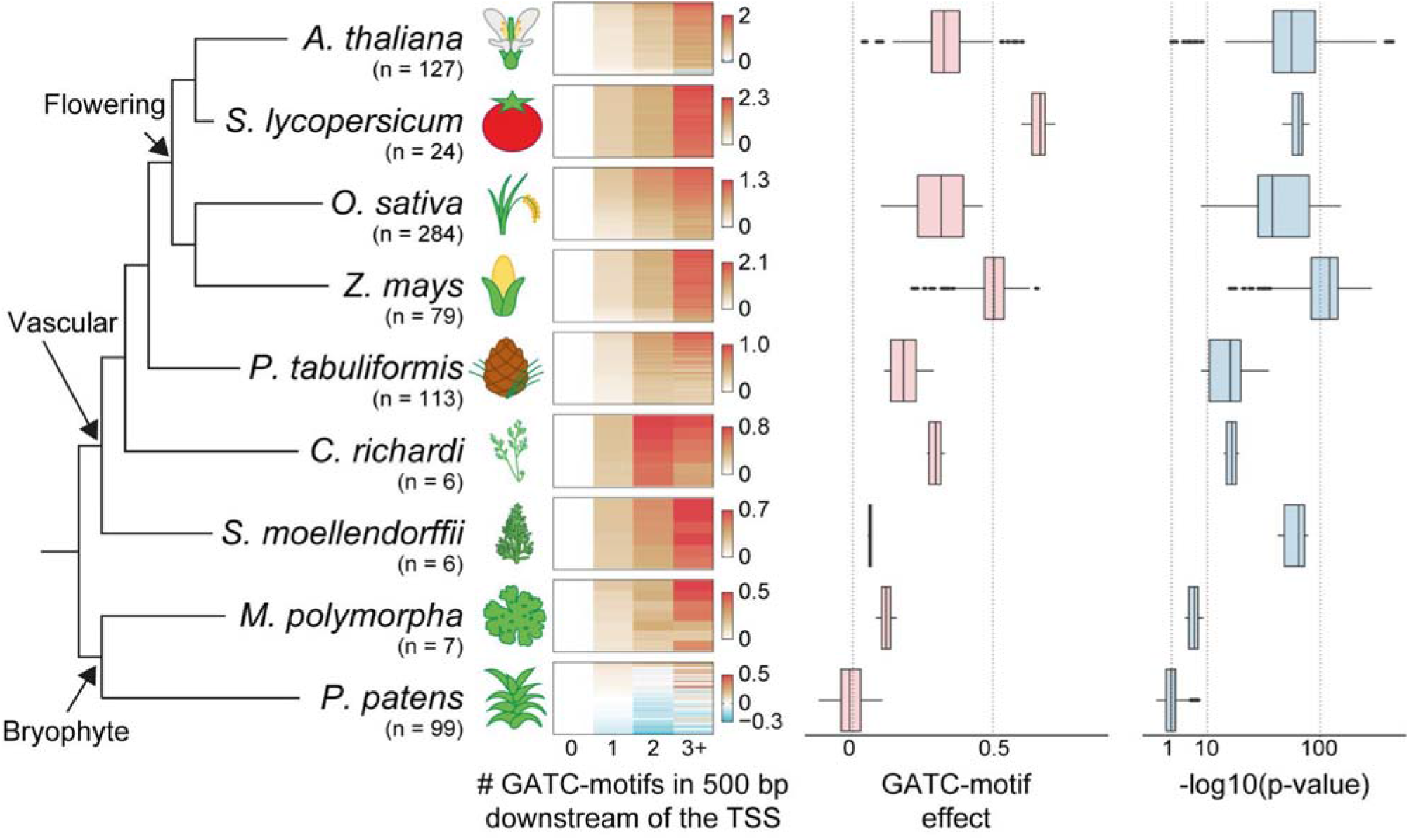
Downstream GATC motif correlates with gene expression in vascular plants. Heatmap of average log_2_(expression change) of multiple transcriptome datasets for genes, categorized by 0, 1, 2, or ≥3 GATC motifs within 500 bp downstream of the TSS, across land plant species. Number of datasets (*n*) indicated. Expression is normalized to the 0-motif group, with species-specific color scales. The effect on expression (slope) and significance of association (p-value) of the GATC motif, as in Fig. 4, are presented. Boxplots show median, IQR, and 1.5x IQR, with outliers. The right x-axis is square-root scaled.

In summary, we have identified the 500 bp region downstream of the TSS as a critical site for transcription regulation for a large fraction of plant genes. We demonstrate that the function of regulatory sequences near the TSS is dependent on their position relative to the TSS, making them distinct from animal enhancers. We further examined a specific downstream GATC-motif that modulates transcription in a dose-dependent manner through GATA TFs. In our analysis, the effect size of the GATC motif surpassed that of any other short DNA motif, even those located upstream of the TSS. The motif apparently acts as a regulatory module, operating much like a rheostat in tuning gene expression between cell types, throughout vascular plants.

Our findings are consistent with previous observations of differences in transcriptional regulation between plants and animals, including special core promoter DNA motifs^41^, expanded^42^ and novel^43^ families of specific and general TFs, and different features of long-range enhancers^44–46^. For instance, plant introns in close proximity to the TSS have been frequently identified as drivers of gene expression^47–49^. Specifically, research into the role of introns has highlighted a motif similar to the GATC motif^15,50,51^.

Our observations on the dependency of enhancer position relative to the TSS are consistent with several previous reports based on individual genes: intron-derived regulatory sequences became inactive when moved upstream of the TSS^15^, and strong upstream enhancers lost activity when moved into the transcribed region^14^. More generally, our results show that regulatory sequences function differently on either side of the TSS in plants, rather than exclusively on one side, as indicated by the lack of, rather than negative, correlation between the effects of the same fragment on either side (Fig. 2D). This may explain why testing enhancers by positioning them in the 3’ UTR of plants results in a strong enrichment of regions from transcribed regions^52,53^. Although this contradicts the common view of the role of the upstream region in controlling expression, the different ways in which enhancers are ‘read’ on either side of the TSS may account for these contrasting results.

One might expect that intragenic enhancers impede RNA polymerase II due to recruitment of DNA-binding TFs to the transcribed region. While the presence of nucleosomes at genes and intronic enhancers in animals indicate that the RNA polymerase can navigate proteins obstructing its path^54^, it remains unclear how enhancers might function differently depending on their positioning relative to the TSS. We propose that the distinct 3D genome architecture in plants, characterized by densely packed genes^55,56^, might create different local environments on either side of the TSS, but many other scenarios can be imagined as well.

Finally, the GATC-motif regulatory program exerts a widespread influence, modulating the gene expression of a significant proportion of genes throughout the plant body. The adaptive advantages this mechanism offers, and how it has evolved across different lineages, promises to be a fertile ground for future exploration.

## Acknowledgment

We thank J. Neuhold, M. Clavel, V. Nizhynska for technical assistance, A. Levy and D. Ben-Tov for help establishing the protoplast system, Y. Eshed and J. Bindics for sharing seeds, and The Plant Sciences and Next Generation Sequencing facilities at the Vienna BioCenter Core Facilities (VBCF). We also thank F. Berger, L. Dolan, A. Stark, RK. Papareddy, BP. de Almeida, K. Hanada, and FK. Lorbeer for fruitful discussions. This work was supported by core funding to MN from the Gregor Mendel Institute, ERA-CAPS grant 1001G+ to MN and DW, and postdoctoral fellowships to YV from the EU Horizon 2020 via the VIP^2^ program and the Marie Skłodowska-Curie individual fellowships (101028014).

## Data availability

Sequencing data have been deposited in the SRA database with accession number PRJNA1009032. Processed data are available in Tables S3 and S4.

## Code availability

Code for data cleaning and analysis is provided as part of the replication package. It is available at https://drive.google.com/drive/folders/1axGoSkiHc-TaO_oiDcxJH397Zo6kcABy?usp=sharing for review. It will be uploaded to the Zenodo repository once the paper has been conditionally accepted.

## References

1. Kumar, S. et al. TimeTree 5: An Expanded Resource for Species Divergence Times. Mol. Biol. Evol. 39, (2022).

2. Knoll, A. H. The Multiple Origins of Complex Multicellularity. Annu. Rev. Earth Planet. Sci. 39, 217–239 (2011).

3. Bonner, J. T. xThe origins of multicellularity. Integrative Biology: Issues, News, and Reviews doi:10.1002/(SICI)1520-6602(1998)1:1<27::AID-INBI4>3.0.CO;2-6.

4. Sebé-Pedrós, A. et al. The Dynamic Regulatory Genome of Capsaspora and the Origin of Animal Multicellularity. Cell 165, 1224–1237 (2016).

5. Meyerowitz, E. M. Plants, animals and the logic of development. Trends Cell Biol. 9, M65–8 (1999).

6. Meyerowitz, E. M. Plants compared to animals: the broadest comparative study of development. Science 295, 1482–1485 (2002).

7. Burgess, D. G., Xu, J. & Freeling, M. Advances in understanding cis regulation of the plant gene with an emphasis on comparative genomics. Curr. Opin. Plant Biol. 27, 141–147 (2015).

8. Kawakatsu, T. et al. Epigenomic Diversity in a Global Collection of Arabidopsis thaliana Accessions. Cell 166, 492–505 (2016).

9. 1001 Genomes Consortium. 1,135 Genomes Reveal the Global Pattern of Polymorphism in Arabidopsis thaliana. Cell 166, 481–491 (2016).

10. Veyrieras, J.-B. et al. High-resolution mapping of expression-QTLs yields insight into human gene regulation. PLoS Genet. 4, e1000214 (2008).

11. Newman, T. C., Ohme-Takagi, M., Taylor, C. B. & Green, P. J. DST sequences, highly conserved among plant SAUR genes, target reporter transcripts for rapid decay in tobacco. Plant Cell 5, 701–714 (1993).

12. Narsai, R. et al. Genome-wide analysis of mRNA decay rates and their determinants in Arabidopsis thaliana. Plant Cell 19, 3418–3436 (2007).

13. O’Malley, R. C. et al. Cistrome and Epicistrome Features Shape the Regulatory DNA Landscape. Cell 165, 1280–1292 (2016).

14. Jores, T. et al. Identification of Plant Enhancers and Their Constituent Elements by STARR-seq in Tobacco Leaves. Plant Cell 32, 2120–2131 (2020).

15. Gallegos, J. E. & Rose, A. B. Intron DNA Sequences Can Be More Important Than the Proximal Promoter in Determining the Site of Transcript Initiation. Plant Cell 29, 843–853 (2017).

16. Norris, S. R., Meyer, S. E. & Callis, J. The intron of Arabidopsis thaliana polyubiquitin genes is conserved in location and is a quantitative determinant of chimeric gene expression. Plant Mol. Biol. 21, 895–906 (1993).

17. Szabo, E. X. et al. Metabolic Labeling of RNAs Uncovers Hidden Features and Dynamics of the Arabidopsis Transcriptome. Plant Cell 32, 871–887 (2020).

18. Jores, T. et al. Synthetic promoter designs enabled by a comprehensive analysis of plant core promoters. Nat Plants 7, 842–855 (2021).

19. Ezer, D. et al. The G-Box Transcriptional Regulatory Code in Arabidopsis. Plant Physiol. 175, 628–640 (2017).

20. Schwechheimer, C., Schröder, P. M. & Blaby-Haas, C. E. Plant GATA Factors: Their Biology, Phylogeny, and Phylogenomics. Annu. Rev. Plant Biol. 73, 123–148 (2022).

21. Riechmann, J. L. et al. Arabidopsis transcription factors: genome-wide comparative analysis among eukaryotes. Science 290, 2105–2110 (2000).

22. Liu, Y. et al. PCSD: a plant chromatin state database. Nucleic Acids Res. 46, D1157–D1167 (2018).

23. Lee, T. A. & Bailey-Serres, J. Integrative Analysis from the Epigenome to Translatome Uncovers Patterns of Dominant Nuclear Regulation during Transient Stress. Plant Cell 31, 2573–2595 (2019).

24. Sidaway-Lee, K., Costa, M. J., Rand, D. A., Finkenstadt, B. & Penfield, S. Direct measurement of transcription rates reveals multiple mechanisms for configuration of the Arabidopsis ambient temperature response. Genome Biol. 15, R45 (2014).

25. Toufighi, K., Brady, S. M., Austin, R., Ly, E. & Provart, N. J. The Botany Array Resource: e-Northerns, Expression Angling, and promoter analyses. Plant J. 43, 153–163 (2005).

26. Klepikova, A. V., Kasianov, A. S., Gerasimov, E. S., Logacheva, M. D. & Penin, A. A. A high resolution map of the Arabidopsis thaliana developmental transcriptome based on RNA-seq profiling. Plant J. 88, 1058–1070 (2016).

27. Hofmann, F., Schon, M. A. & Nodine, M. D. The embryonic transcriptome of Arabidopsis thaliana. Plant Reprod. 32, 77–91 (2019).

28. Schmid, M. et al. A gene expression map of Arabidopsis thaliana development. Nat. Genet. 37, 501–506 (2005).

29. Schneider, A. et al. Potential targets of VIVIPAROUS1/ABI3-LIKE1 (VAL1) repression in developing Arabidopsis thaliana embryos. Plant J. 85, 305–319 (2016).

30. Shahan, R. et al. A single-cell Arabidopsis root atlas reveals developmental trajectories in wild-type and cell identity mutants. Dev. Cell 57, 543–560.e9 (2022).

31. Zhang, T.-Q., Chen, Y. & Wang, J.-W. A single-cell analysis of the Arabidopsis vegetative shoot apex. Dev. Cell 56, 1056–1074.e8 (2021).

32. Reyes, J. C., Muro-Pastor, M. I. & Florencio, F. J. The GATA family of transcription factors in Arabidopsis and rice. Plant Physiol. 134, 1718–1732 (2004).

33. Zhang, S. et al. Spatiotemporal transcriptome provides insights into early fruit development of tomato (Solanum lycopersicum). Sci. Rep. 6, 23173 (2016).

34. Stelpflug, S. C. et al. An Expanded Maize Gene Expression Atlas based on RNA Sequencing and its Use to Explore Root Development. Plant Genome 9, (2016).

35. Xia, L. et al. Rice Expression Database (RED): An integrated RNA-Seq-derived gene expression database for rice. J. Genet. Genomics 44, 235–241 (2017).

36. Perroud, P.-F. et al. The Physcomitrella patens gene atlas project: large-scale RNA-seq based expression data. Plant J. 95, 168–182 (2018).

37. Xiao, Y.-L. & Li, G.-S. Differential expression and co-localization of transcription factors during the indirect de novo shoot organogenesis in the fern Ceratopteris richardii. Research Square (2023) doi:10.21203/rs.3.rs-2531906/v1.

38. Niu, S. et al. The Chinese pine genome and methylome unveil key features of conifer evolution. Cell 185, 204–217.e14 (2022).

39. Huang, L. & Schiefelbein, J. Conserved Gene Expression Programs in Developing Roots from Diverse Plants. Plant Cell 27, 2119–2132 (2015).

40. Sharma, N., Bhalla, P. L. & Singh, M. B. Transcriptome-wide profiling and expression analysis of transcription factor families in a liverwort, Marchantia polymorpha. BMC Genomics 14, 915 (2013).

41. Kumari, S. & Ware, D. Genome-wide computational prediction and analysis of core promoter elements across plant monocots and dicots. PLoS One 8, e79011 (2013).

42. Shin-Han, S. A. D. M.-C. S. A. D. L. Transcription Factor Families Have Much Higher Expansion Rates in Plants than in Animals1. 139, 18–26 (2005).

43. Blanc-Mathieu, R., Dumas, R., Turchi, L., Lucas, J. & Parcy, F. Plant-TFClass: a structural classification for plant transcription factors. Trends Plant Sci. (2023) doi:10.1016/j.tplants.2023.06.023.

44. Weber, B., Zicola, J., Oka, R. & Stam, M. Plant Enhancers: A Call for Discovery. Trends Plant Sci. 21, 974–987 (2016).

45. Lu, Z. et al. The prevalence, evolution and chromatin signatures of plant regulatory elements. Nat Plants 5, 1250–1259 (2019).

46. Schmitz, R. J., Grotewold, E. & Stam, M. Cis-regulatory sequences in plants: Their importance, discovery, and future challenges. Plant Cell 34, 718–741 (2022).

47. Rose, A. B. Intron-mediated regulation of gene expression. Curr. Top. Microbiol. Immunol. 326, 277–290 (2008).

48. Meng, F. et al. Genomic editing of intronic enhancers unveils their role in fine-tuning tissue-specific gene expression in Arabidopsis thaliana. Plant Cell 33, 1997–2014 (2021).

49. Rose, A. B. Introns as Gene Regulators: A Brick on the Accelerator. Front. Genet. 9, 672 (2018).

50. Back, G. & Walther, D. Identification of cis-regulatory motifs in first introns and the prediction of intron-mediated enhancement of gene expression in Arabidopsis thaliana. BMC Genomics 22, 390 (2021).

51. Gallegos, J. E. & Rose, A. B. An intron-derived motif strongly increases gene expression from transcribed sequences through a splicing independent mechanism in Arabidopsis thaliana. Sci. Rep. 9, 13777 (2019).

52. Tan, Y. et al. Genome-wide enhancer identification by massively parallel reporter assay in Arabidopsis. Plant J. 116, 234–250 (2023).

53. Sun, J. et al. Global Quantitative Mapping of Enhancers in Rice by STARR-seq. Genomics Proteomics Bioinformatics 17, 140–153 (2019).

54. Zabidi, M. A. & Stark, A. Regulatory Enhancer-Core-Promoter Communication via Transcription Factors and Cofactors. Trends Genet. 32, 801–814 (2016).

55. Liu, C. et al. Genome-wide analysis of chromatin packing in Arabidopsis thaliana at single-gene resolution. Genome Res. 26, 1057–1068 (2016).

56. Lee, H. & Seo, P. J. Accessible gene borders establish a core structural unit for chromatin architecture in Arabidopsis. Nucleic Acids Res. gkad710 (2023).

