## Supplementary material and methods for "Widespread position-dependent transcriptional regulatory sequences in plants"

### Supplementary Materials:

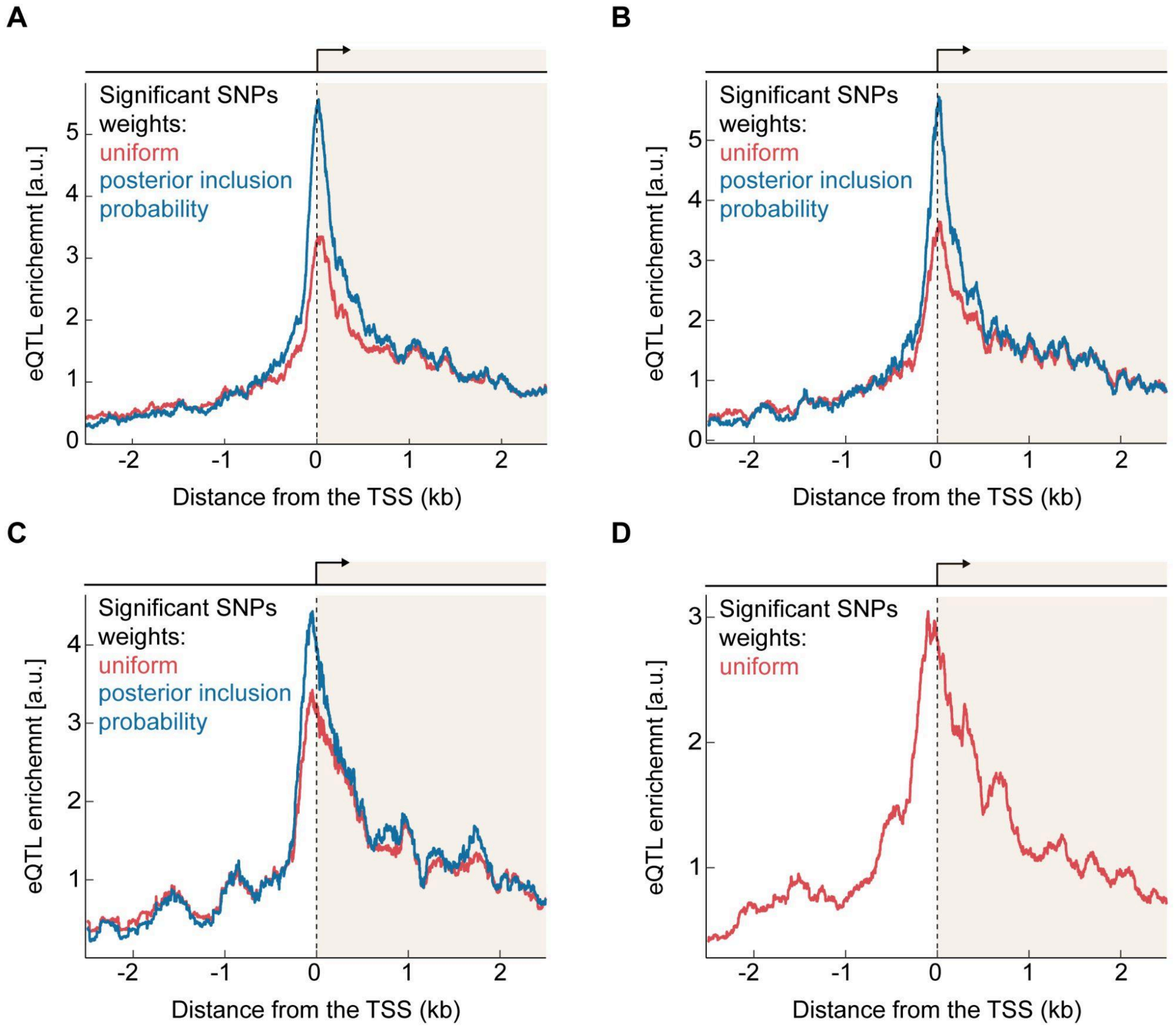

**Supplementary Figure S1: Consistent eQTL enrichment near the TSS of genes across multiple datasets**

eQTL enrichment near the transcription start site (TSS) of genes from various data sets: **(A)** Second batch from ref. (*1*) (4,259 genes with significant SNPs), **(B)** First batch from ref. (*1*) (2,760 genes with significant SNPs) **(C)** Ref. (*2*) (639 genes with significant SNPs), and **(D)** Ref. (*3*) (3,048 genes with significant SNPs). SNPs linked to the expression of the same gene were normalized to ensure equal contribution from each gene in the analysis. Weights were assigned either uniformly (red line), or based on the posterior inclusion probability (PIP, blue line), which accounts for linkage disequilibrium (LD) between SNPs. Incorporating LD using PIP consistently enhanced the enrichment in and downstream of the TSS across datasets. Data smoothed with a 100 bp (data from ref. (*1*)) or 200 bp (otherwise) rolling window.

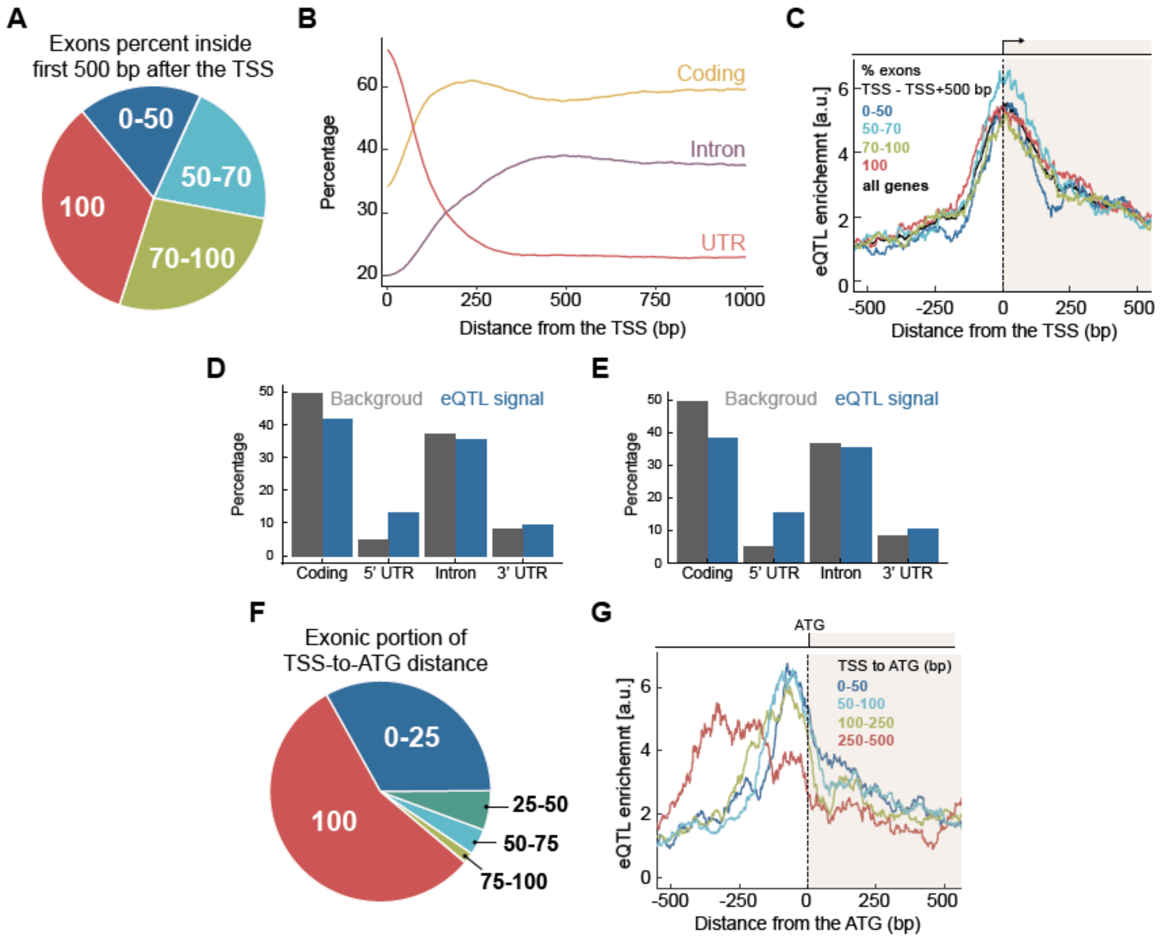


**Supplementary Figure S2: No association between exon-intron structure and eQTL enrichment near TSS**

**(A)** Genes grouped according to fraction of exonic sequence in the 500 bp following the TSS for *A. thaliana* genes. Four groups represent gene portions with different exon content: up to 50%, 50%-70%, 70%-100%, and 100%. **(B)** Percentage of genomic sequence coverage by different genomic features as a function of distance from the TSS **(C)** eQTL enrichment near TSS for genes with varying exonic fraction within the first 500 bp after TSS, shown as in Fig. 1A. Gene counts per group: 914 (0%-50%), 1,044 (50%-70%), 1,179 (70%-100%), 1,102 (100%). **(D-E)** The proportion of eQTL signals within transcripts, as determined by the posterior inclusion probability across various genomic features, compared to the total length of these features in genes where significant associations have been found. Plotted for first **(E)** or second **(F)** batch from ref. (*1*). **(F)** Genes grouped according to fraction of exonic (5' UTR) sequence in the TSS-to-ATG regions for *A. thaliana* genes. Five groups represent gene portions with different exonic fractions: 0%-25%, 25%-50%, 50%-75%, 75%-100%, and 100%. **(G)** eQTL enrichment for genes with different TSS-to-ATG distances, as in Fig. 1B, with data aligned to the ATG and not the TSS. In **A,C,** and **F**, groups exclude the upper limit, i.e., A%-B% represents A%≤x<B%.

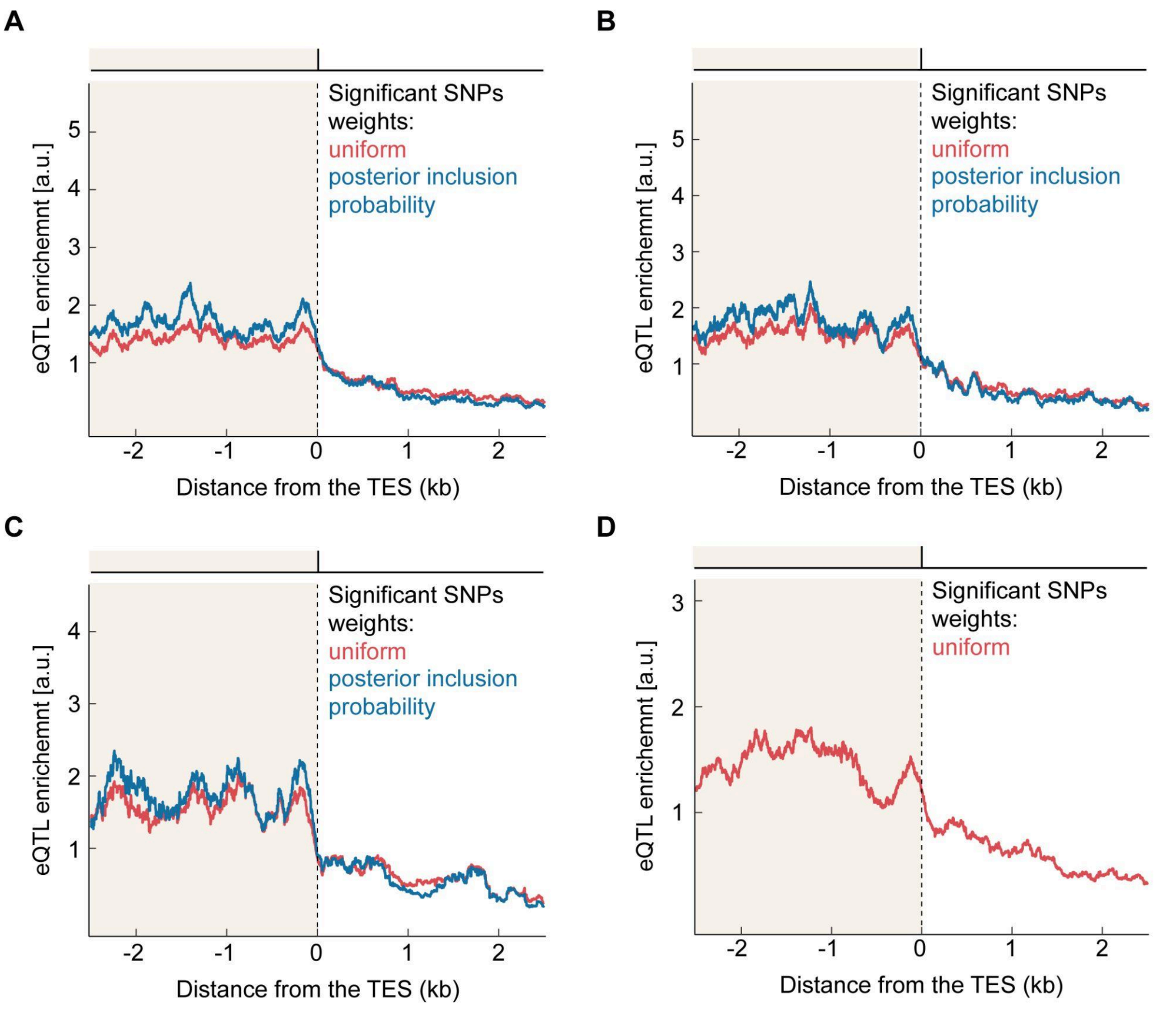


**Supplementary Figure S3: eQTL enrichment near the transcription end site of genes**

eQTL enrichment is depicted near the transcription end site (TES) of genes from various datasets, using the same methodology and details as described in Fig. S1. The y-axis scaling is the same as in the respective sub-plot in Fig. S1 to enable comparisons between the figures. The enrichment observed near the TSS is not found at the other end of genes, in the TES.

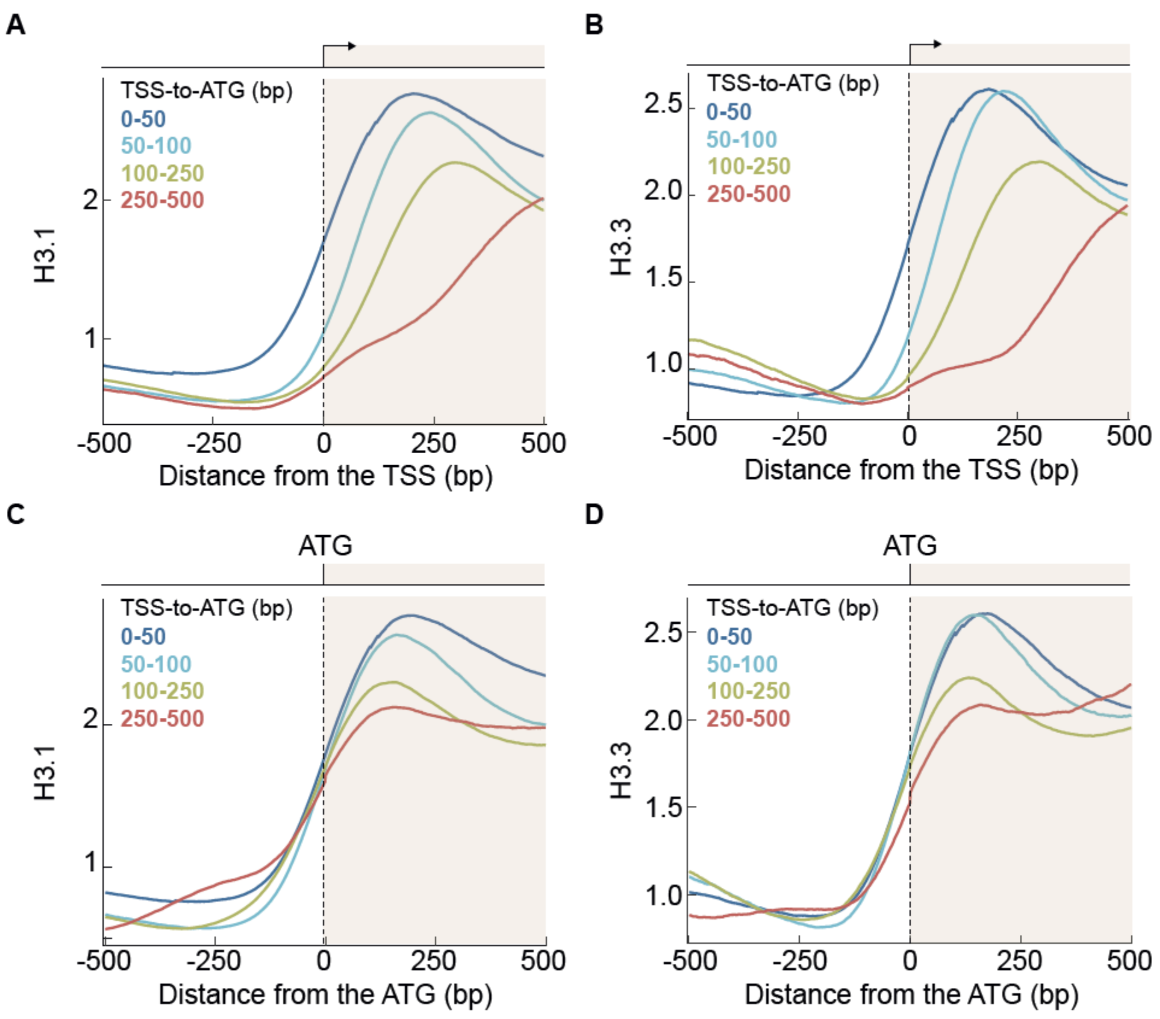


**Supplementary Figure S4: TSS-to-ATG distance influences histone H3 enrichment near the TSS**

Average enrichments of histones H3.1 **(A, C)** and H3.3 **(B, D)** near the TSS **(A-B)** or ATG **(C-D)** of genes with varying TSS-to-ATG distances. Processed data for enrichment of the two histones along the genome were retrieved from the plant chromatin state database (PCSD)(*4*), and data from replicates were averaged (*5*, *6*).

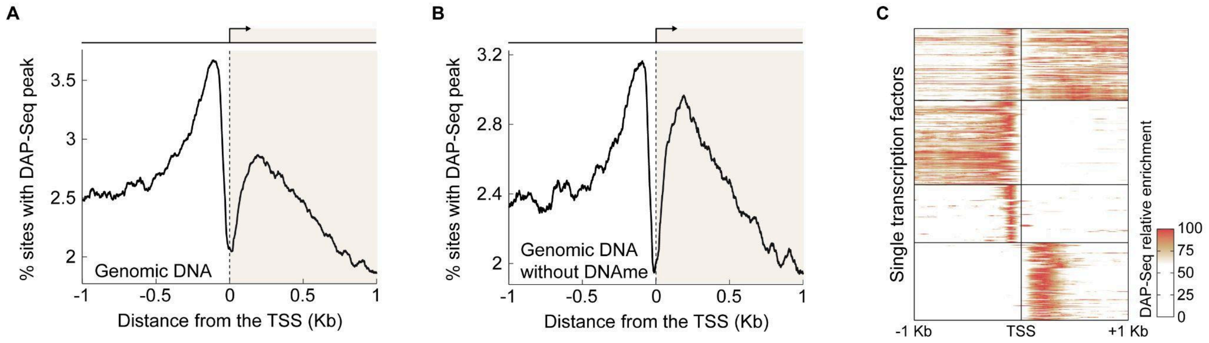


### Supplementary Figure S5: TFs bind bimodally both up- and downstream of the TSS (DAP-Seq)

**(A-B)** Fraction of sites with a DAP-Seq peak center for TFs, as in Fig. 1C, separately plotted for TFs binding purified genomic DNA **(A)** or to non-methylated genomic DNA **(B)**; data smoothed using a 100 bp rolling window. TFs binding downstream to the TSS were more sensitive to DNA methylation. **(C)** DAP-Seq peak enrichment, as in Fig. 1D, for each TF separately, maximum signal for each TF was scaled to 100, enrichment patterns were clustered using k-means (k = 4).

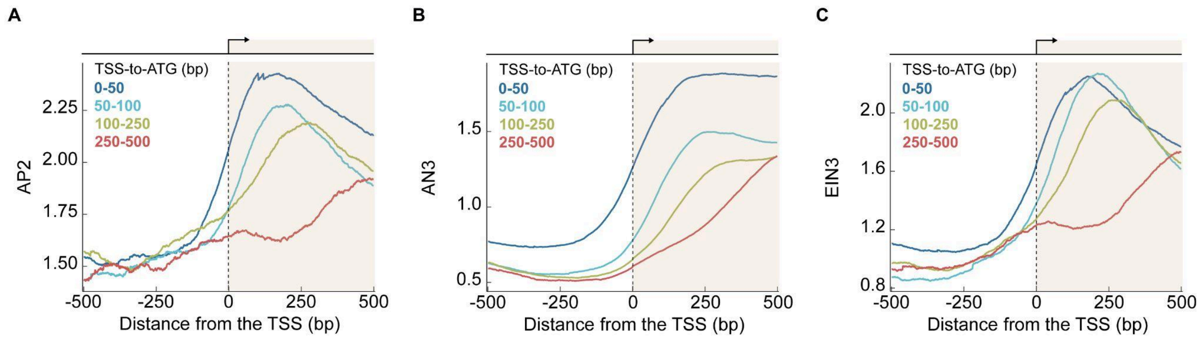


### Supplementary Figure S6: In vivo evidence of TFs binding downstream of the TSS

Average enrichment of **(A)** the DNA-binding protein APETALA 2 (AP2), **(B)** the transcriptional coactivator ANGUSTIFOLIA3 (AN3), and **(C)** the DNA-binding protein ETHYLENE INSENSITIVE3 (EIN3) near the TSS of genes with varying TSS-to-ATG distances. Data were retrieved from PCSD, averaged for replicates if available, and plotted as in Fig. S4A-B (*4*, *7*–*9*).

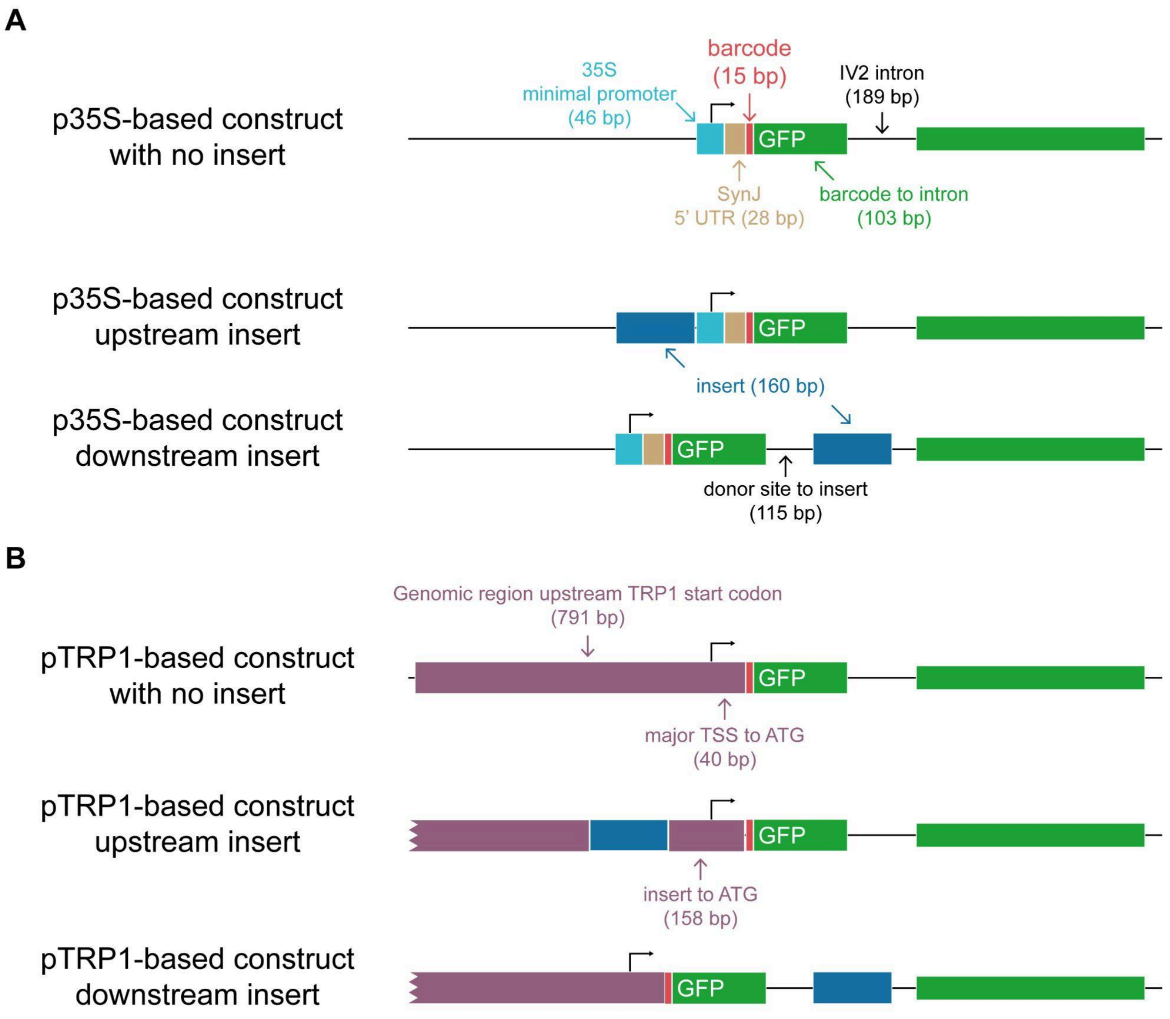


**Supplementary Figure S7: MPRA construct sequence layout**

The layout of the constructs used in the MPRA assay, based on a published design (*10*), for both p35S-based **(A)** and pTRP1-based **(B)** versions. 'No insertion' control constructs are displayed at the top, followed by constructs with upstream and downstream insertions (dark blue). Each construct includes a GFP coding region (green) with an IV2 intron, as well as a 15 bp barcode (red). Distances and lengths of genomic attributes are provided. **(A)** The p35S-based construct incorporates the minimal CaMV 35S core promoter (light blue), followed by the SynJ synthetic 5’ UTR (light brown) (*11*), with the upstream insertion placed just before the minimal core promoter. **(B)** The pTRP1-based construct consists of the 791 bp genomic sequence preceding the coding region of the TRP1 gene (purple), which includes upstream proximal promoter, the core promoter, and 5’ UTR. The upstream insertion site is situated within this sequence, 118 bp upstream of the major TSS and 40 bp upstream of the TSS annotation in the TAIR10 database.

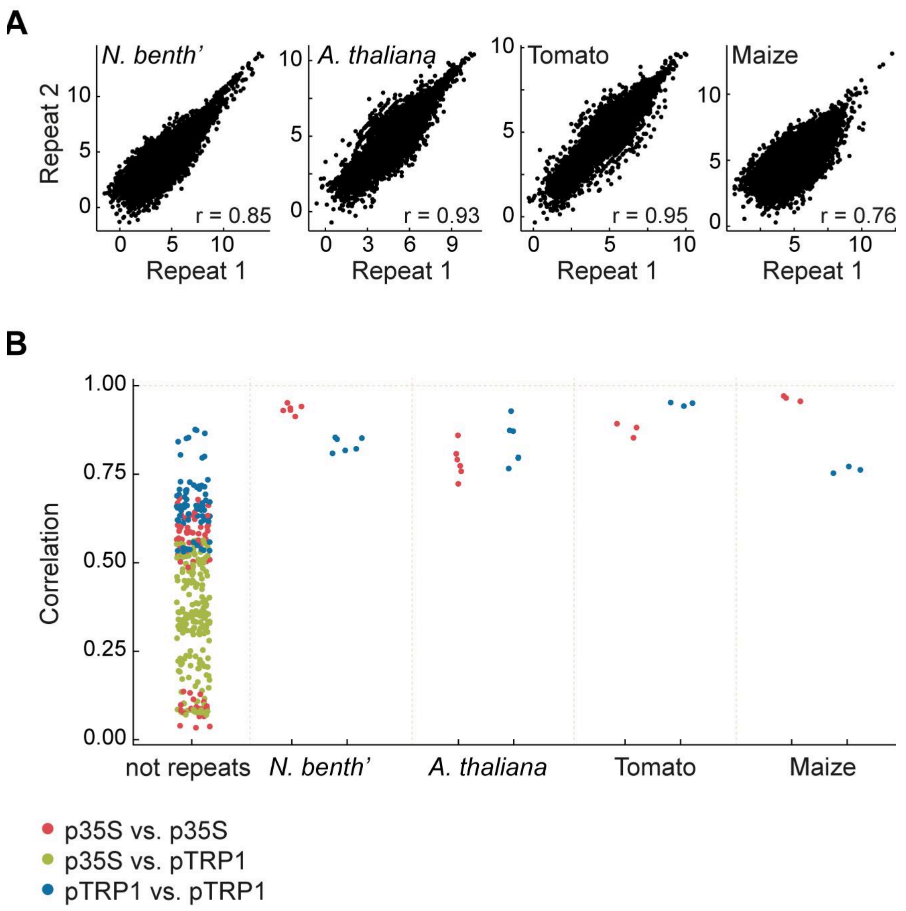


### Supplementary Figure S8: Reproducibility of MPRA across four flowering plants

MPRA experiments were conducted in four (*A. thaliana* and *N. benthamiana*) or three (tomato and maize) replicates, with both p35S- and pTRP1-based libraries. Pearson's correlation coefficients were calculated for comparison of all 24,000 constructs for all possible pairs within the 28 experiments. **(A)** Selected scatter plots are presented for pairs of pTRP1-based replicates for each of the four species, with the respective Pearson's correlation coefficients indicated (*r*). **(B)** The full array of Pearson's correlation coefficients for replicates plotted separately for each species, and compared to correlation values between pairs of non-replicates.

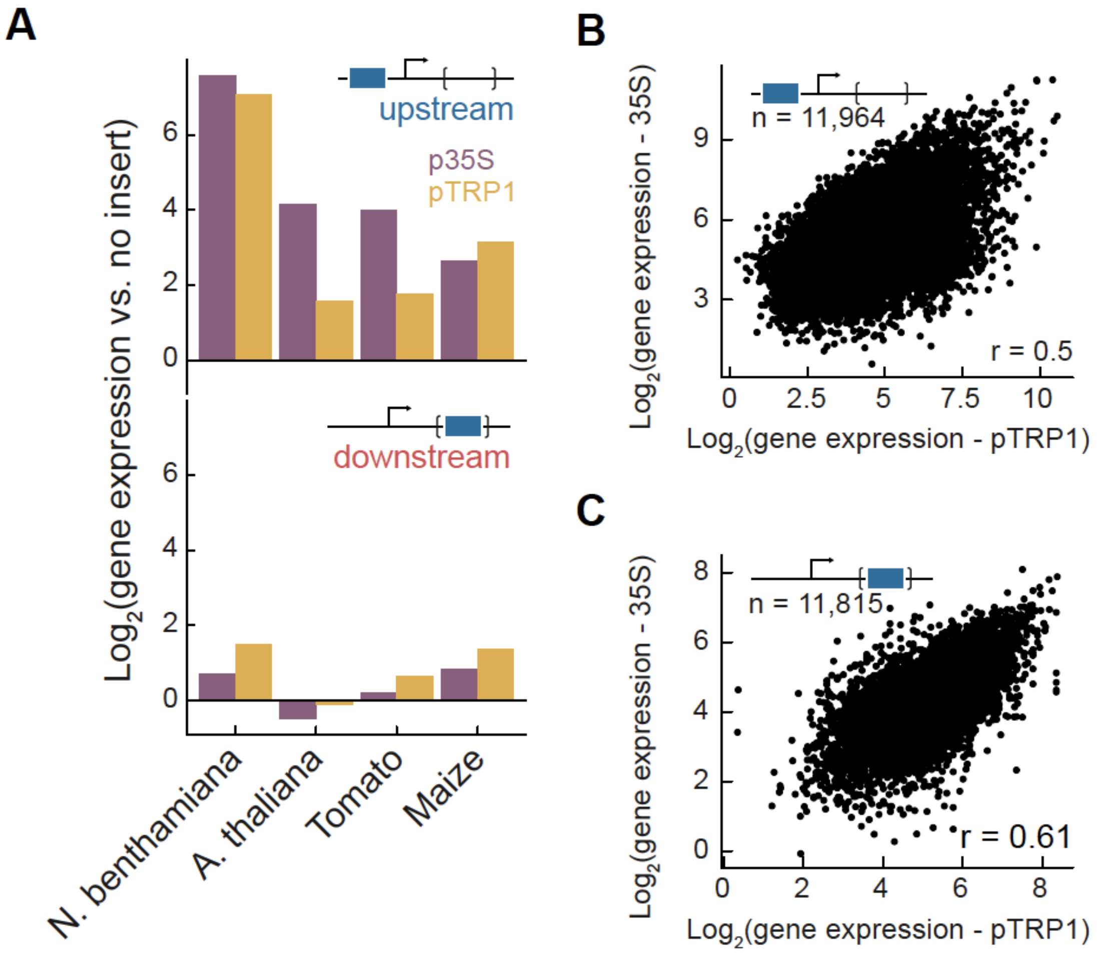


**Supplementary Figure S9: Comparison of MPRA results using two core promoters**

**(A)** Shown is the gene expression induced by 35S enhancer fragments vs. constructs lacking insertions in four species within the MPRA framework. Shown is the comparison of gene expression between the two core promoters used in the MPRA: 35S (purple) and the TRP1 gene promoter (orange). The effects are shown for insertion upstream of the TSS (top panel) and downstream of it within an intron (bottom panel). **(B-C)** Shown is the correlation in gene expression across all constructs, across the 12,000 inserted fragments, when comparing the use of the 35S core promoter vs. the TRP1 promoter within the Arabidopsis MPRA. The correlation is shown for insertions upstream **(B)** and downstream of the TSS within an intron **(C)**. Pearson's correlation (*r*) and number of constructs compared (*n*) are indicated in **B** and **C**.

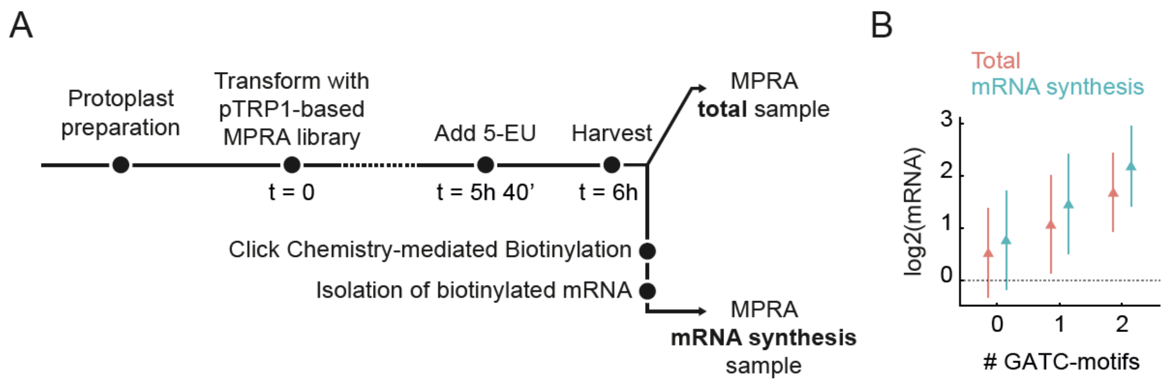


### Supplementary Figure S10: mRNA synthesis rate with MPRA

**(A)** Measurements of mRNA synthesis rate with MPRA, experimental setup: Arabidopsis protoplasts were transformed with pTRP1-based libraries containing 12,000 fragments positioned downstream of the TSS and incubated for 5 h 40 min at room temperature with constant light. After addition of 5-ethynyl uridine (5-EU) and 20 min incubation (*12*), total RNA was extracted. Total mRNA was used for regular MPRA. Newly synthesized mRNA was isolated by click reactions followed by selection of biotinylated mRNA using beads. The isolated mRNA was used to generate MPRA libraries. The experiment was performed in two repeats. **(B)** Relative activity of downstream fragments inserted into pTRP1-based constructs as a function of the number of YVGATCBR consensus motifs, as in Fig. 3C. Group sizes: 6,855 (no motif), 956 (1 motif), 119 (2 motifs). Data represent average signal from two replicates of mRNA synthesis MPRA experiments for total mRNA and newly synthesized mRNA. Error bars represent the mean ± 1 standard deviation.

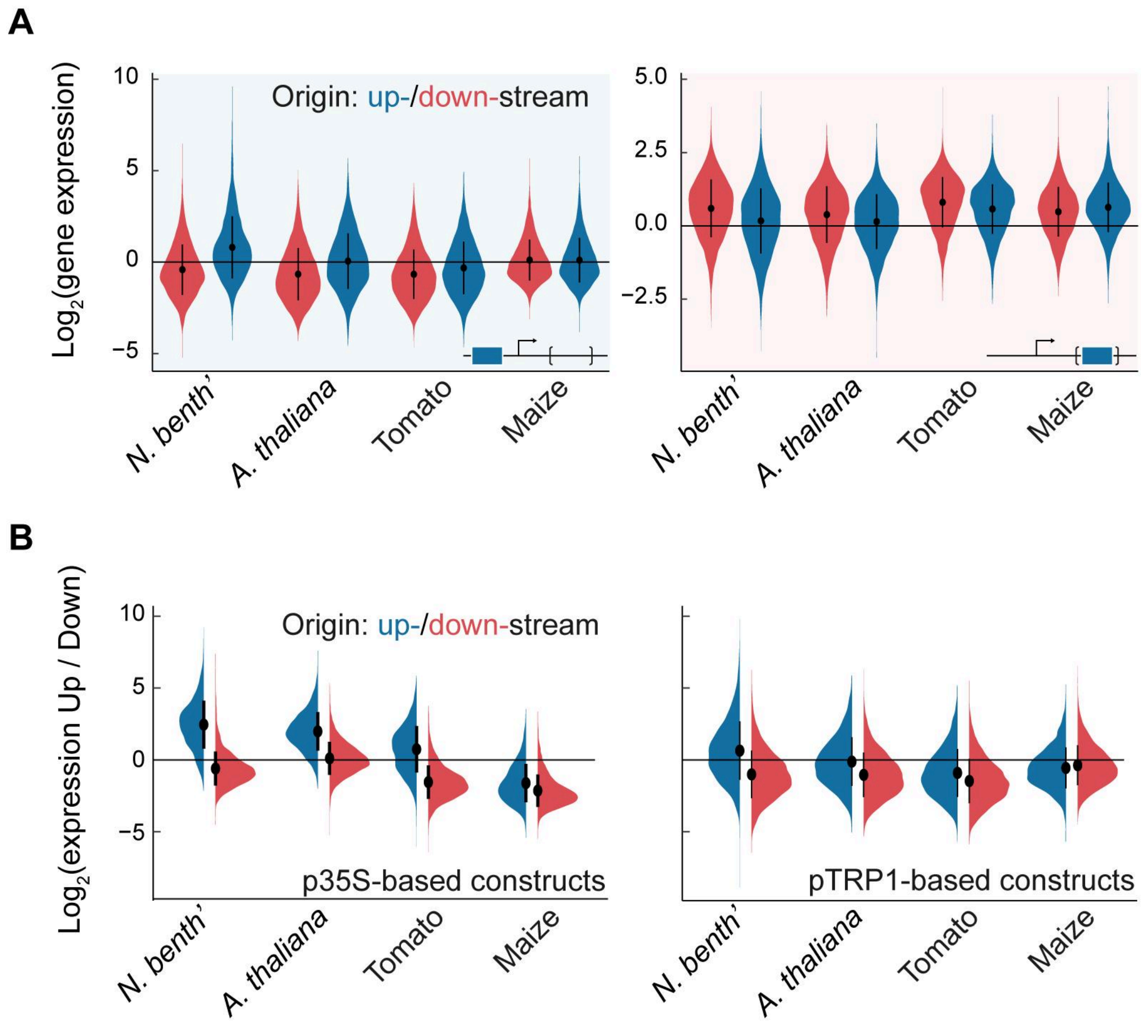


**Supplementary Figure S11: Comparative enhancement by upstream vs. downstream originating sequences**

**(A)** Same as Fig. 2F for pTRP1-based constructs. Expression from upstream- (blue, 3770±211 fragments) and downstream-derived (red, 7,914±26 fragments) fragments is compared against constructs lacking an insertion, with sequences situated upstream (left) or downstream (right). **(B)** Relative expression from identical enhancers due to enhancer position (upstream vs. downstream) separated by fragment genomic origin: upstream (blue, 3766±215 fragments) and downstream (red, 7906±33 fragments), in four different species, for both p35S-based (left) and pTRP1-based (right) libraries. In both A and B, error bars represent the mean and ± standard deviation.

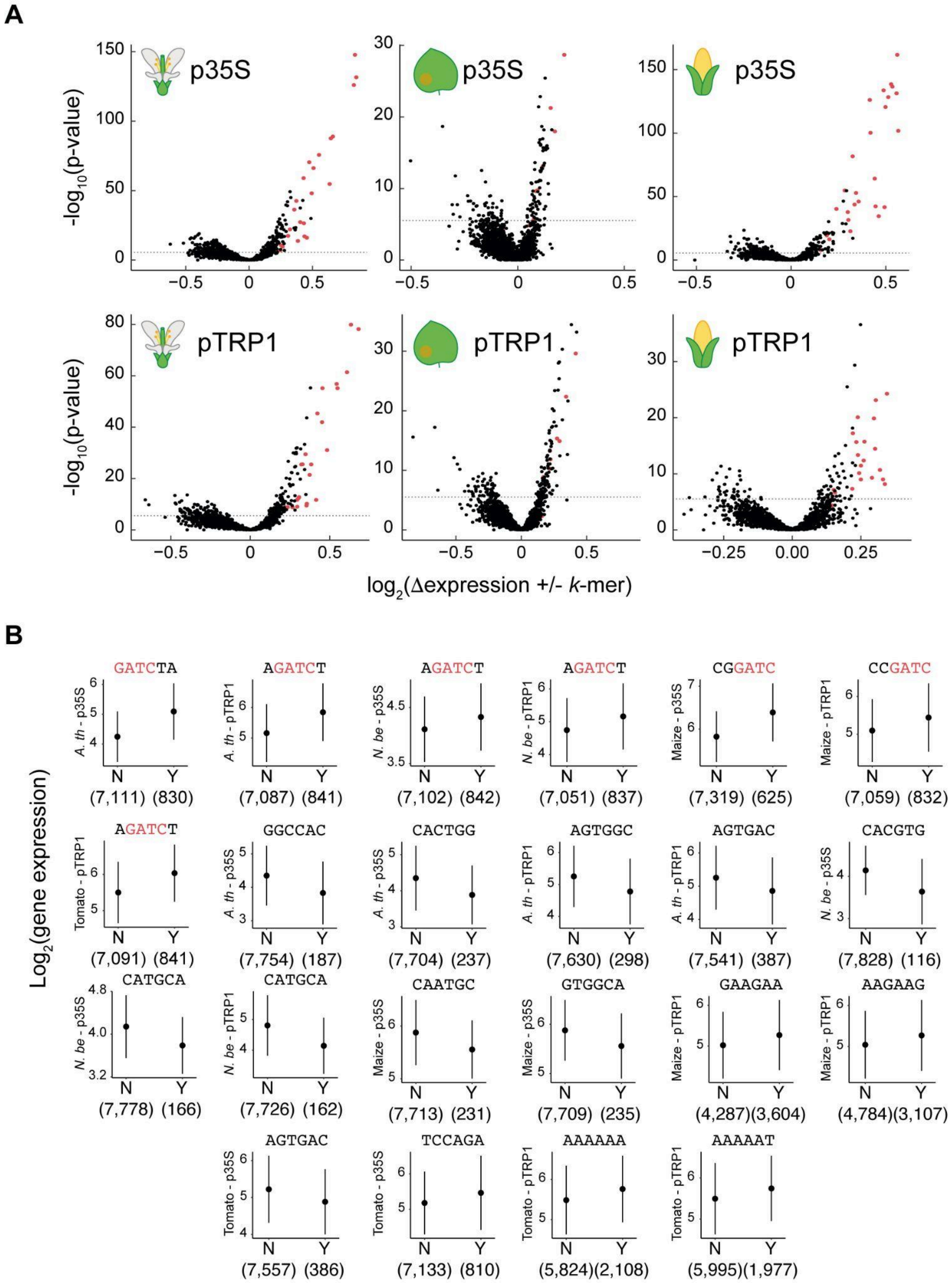


**Supplementary Figure S12: Identifying 6-mers linked to downstream MPRA expression**

**(A)** Score of 6-mers based on their influence on gene expression in the TSS downstream-MPRA, considering only sequences derived from downstream of the TSS in the Arabidopsis genome, as shown in Fig. 3A. For each of the 2,080 unique 6-mers, including reverse complements, downstream-derived sequences were divided into those containing or lacking the 6-mer. A $-\log_{10}$(p-value) from a Mann-Whitney U test comparing these two groups is plotted on the y-axis, versus the difference in average $\log_2$ expression between the groups containing and lacking the 6-mer on the x-axis. Bonferroni multiple testing 5% threshold (depicted by a horizontal dashed line) is defined as $-\log_{10}$(0.05 / # tests), where the number of tests includes all eight experimental setups. Points in red depict 6-mers containing the sequence GATC. The top row shows p35S-based backbones and the bottom row pTRP1-based ones. Host species from left to right are: *A. thaliana*, *N. benthamiana*, and maize, as indicated by icons. **(B)** Distributions of expression values for downstream-derived fragments with (Y) or without (N) the indicated 6-mer in downstream MPRA, with the number of fragments indicated below the x-axis. Effects of 22 6-mers for specific combinations of backbone x species are shown, similar to Fig. 3B. All 6-mers presented have a p-value smaller than $10^{-12}$. The GATC-containing 6-mer with the strongest effect, and the top two GATC-lacking 6-mers for each of the eight backbone-species combinations are displayed, comprising a set of 24 6-mers, which includes the 2 6-mers presented in Fig. 3B. Error bars represent the mean ± standard deviation. Species and backbone are indicated on the y-axis.

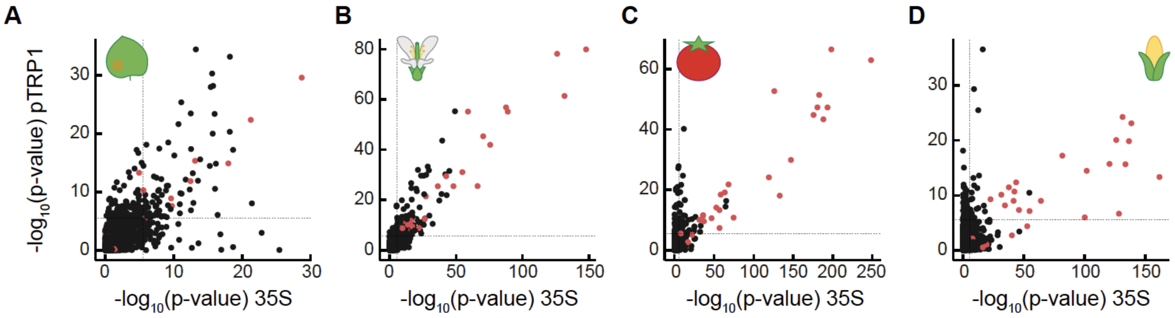


**Supplementary Figure S13: Comparison of 6-mers linked to downstream MPRA expression between p35S- and pTRP1-based libraries**

Comparison of (-log10) p-values for 6-mer associations with gene expression, as in Fig. S12A, between the p35S- and pTRP1-based libraries. The p-values are shown for MPRA done in the different species: *N. benthamiana* **(A)**, Arabidopsis **(B)**, tomato **(C)**, and maize **(D)**. A dashed line, as defined in Fig. S12A, represents the Bonferroni-corrected 5% significance threshold. Points highlighted in red represent 6-mers containing the sequence GATC.

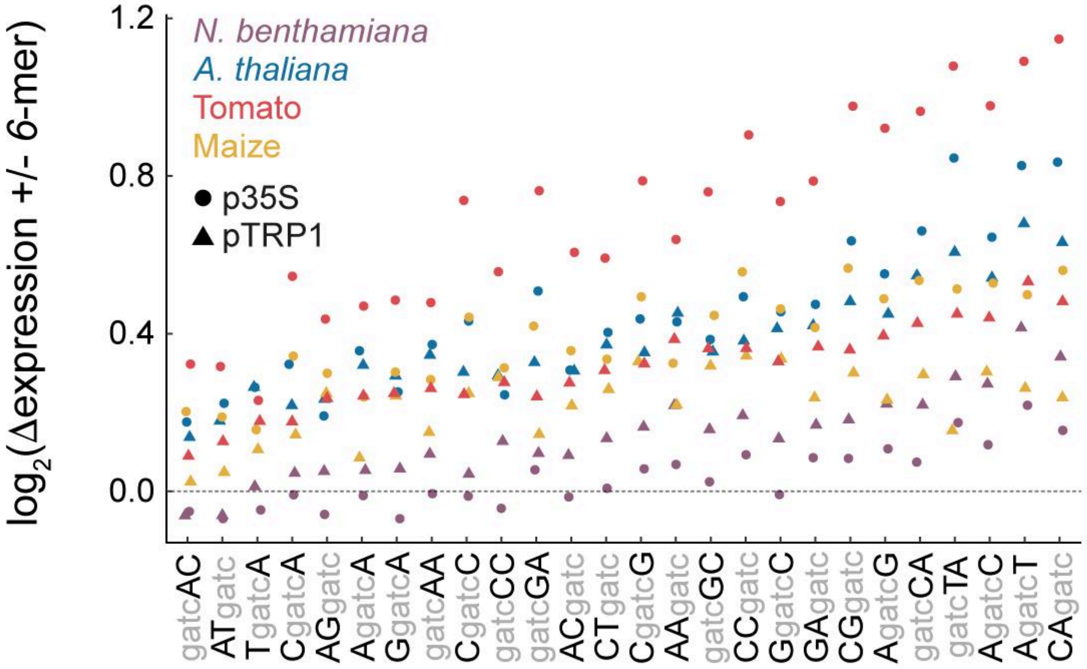


**Supplementary Figure S14: Influence of GATC containing-sequences on downstream-MPRA expression**

Examination of the effect of GATC motif's impact on MPRA expression levels using 6-mers. For all 26 unique 6-mer sequences containing a GATC, including their reverse complements, the average $\log_2$ expression difference between sequences having the 6-mer and those lacking the 6-mer is plotted. 6-mers are ordered based on their median effect across the eight experimental setups. Shape indicates the construct backbone (p35S or pTRP1), and color represents the host species. The six 6-mers (CAgatc, AgatcT, AgatcC, gatcTA, gatcCA, and AgatcG) with the highest median effect were used to establish the consensus YV**GATC**BR sequence defined as the GATC motif, which was then used in downstream analyses.

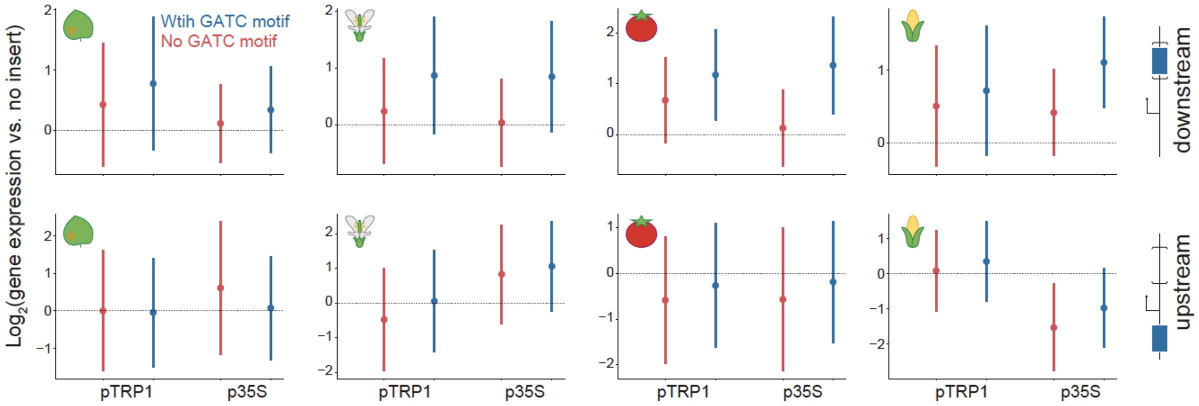


### Supplementary Figure S15: Comparison of GATC motif association with gene expression in the upstream and downstream MPRA

Relative activity of all fragments when inserted downstream (upper row) or upstream (bottom row), as a function of the presence of YVGATCBR consensus motifs in the tested fragments. Group sizes: 10,214-10,675 (No GATC motif, red) and 1,281-1,303 (With GATC motif, blue). Backbone and species are indicated. Error bars represent mean ± 1 standard deviation.

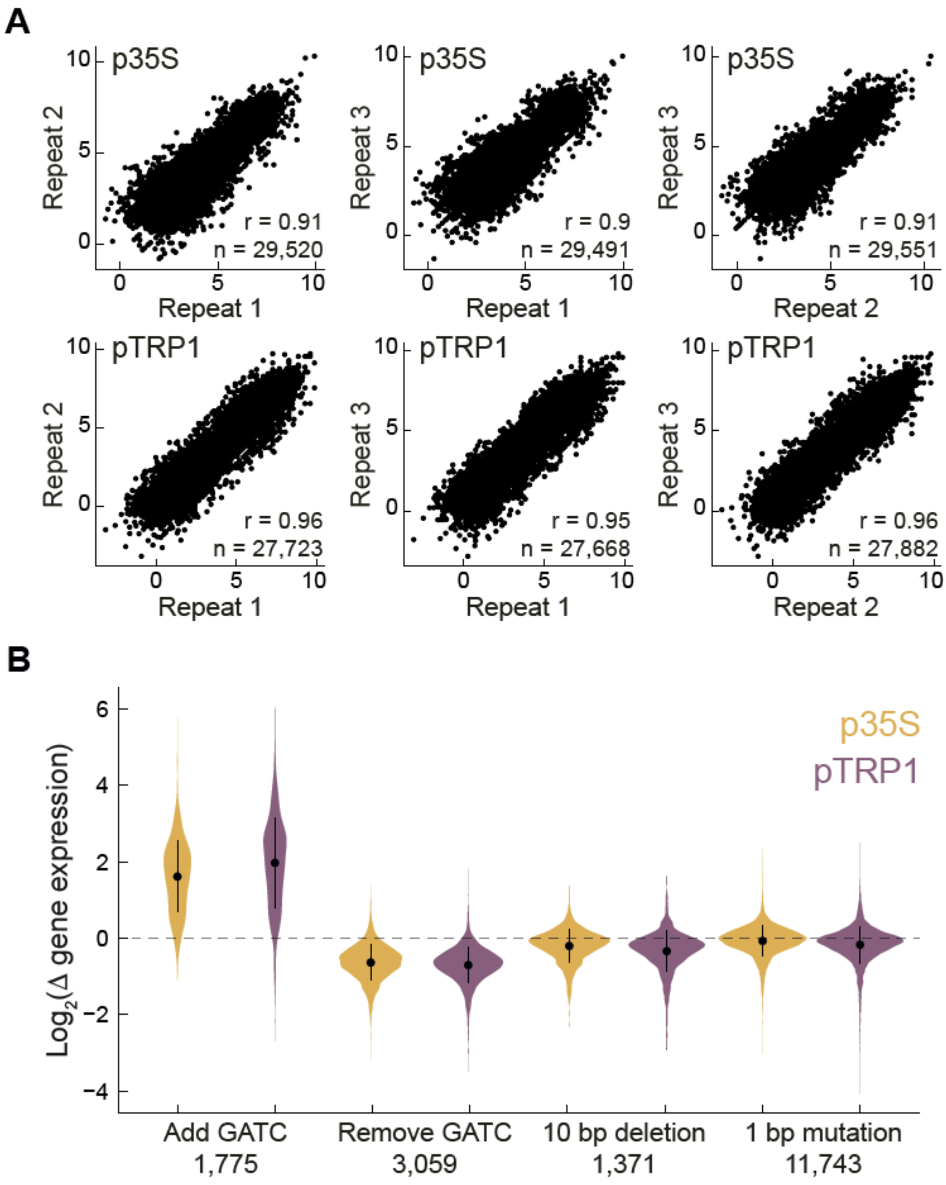


**Supplementary Figure S16: MPRA of mutated sequences**

MPRA experiments were performed in Arabidopsis protoplasts using synthetically mutated sequences inserted only in the downstream position. Both p35S- and pTRP1-based libraries were used. The libraries contained 30,000 fragments: 12,000 from the initial pool and an additional 18,000 fragments, each being a variant of one of the original fragments. **(A)** Comparison between each pair from the three replicates, displaying results with the p35S-based library at the top and the pTRP1-based library at the bottom. Pearson's correlation coefficients (*r*) and numbers of compared fragments (*n*) are indicated on each graph. **(B)** $Log_2$ expression ratio between mutated fragments and their original fragment. Mutations encompass: addition of a GATC motif, removal of a GATC motif, 10 bp deletions, and 1 bp changes, which include both deletions and nucleotide substitutions. Color represents the library type, either p35S- or pTRP1-based; numbers below the x-axis indicate the number of fragments in each category. Error bars depict the mean with ± standard deviation.

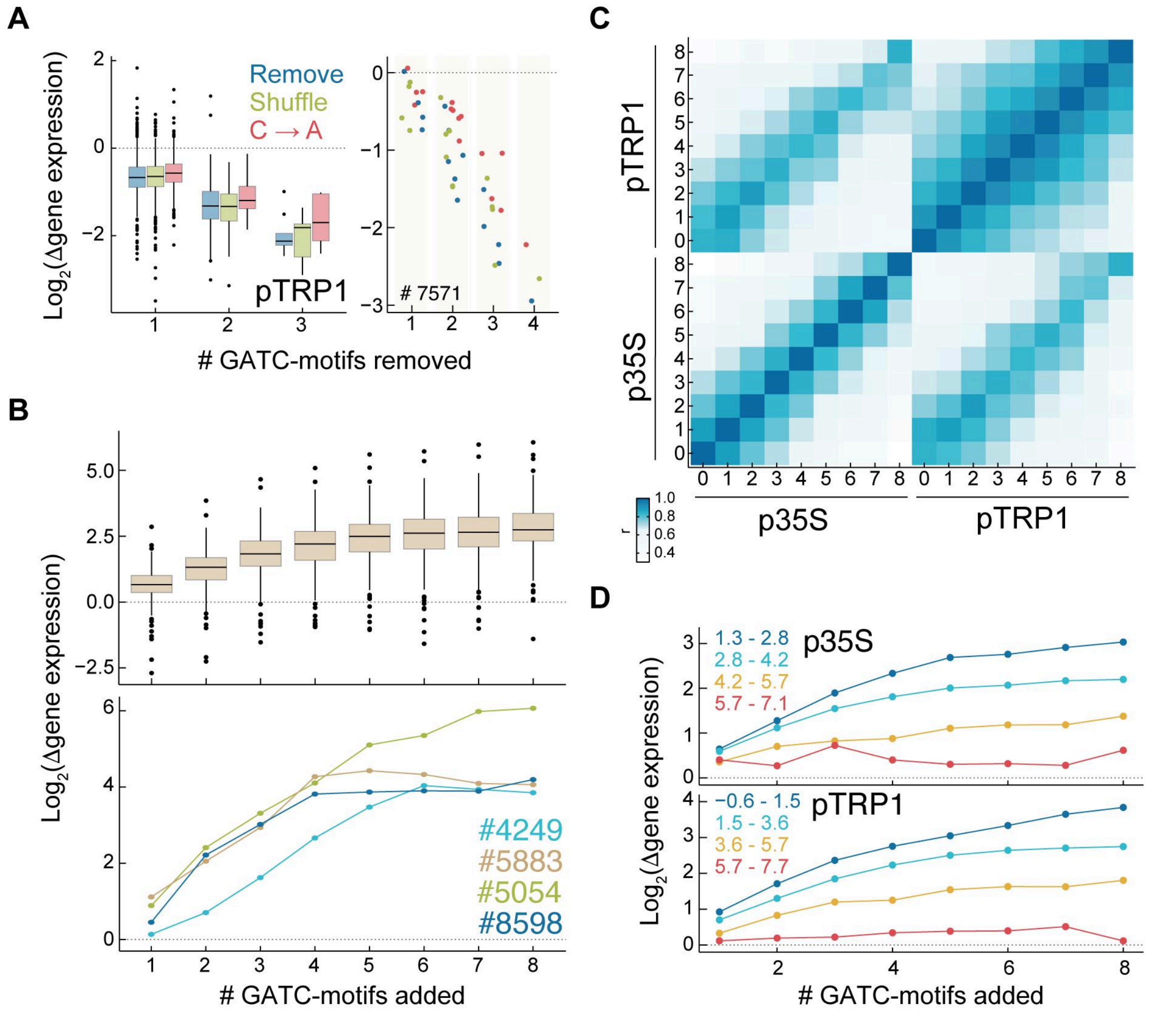


**Supplementary Figure S17: Impact of GATC-motif mutations on gene expression in MPRA**

**(A)** Expression change after removing GATC motifs in a pTRP1-based library, plotted as in Fig. 3D. Illustrated effects across 823 fragments (left) and an example fragment with originally 4 motifs (right). **(B)** Expression changes due to GATC-motifs additions in a pTRP1-based library, mirroring Fig. 3F. Effects across 221 fragments (top) and 4 specific examples (bottom). **(C)** Pearson's correlation coefficients between expression of 221 fragments with varying GATC-motif additions in p35S- and pTRP1-based libraries. **(D)** Influence of GATC-motif additions depends on initial sequence expression. Depicted is the average $\log_2$ expression difference upon motif addition relative to original fragments, for different copy numbers of the added motif. Color indicates original fragment ($\log_2$) expression level ranges, as indicated. Plotted for p35S- (top) and pTRP1-based libraries (bottom). Boxplots in **A-B** represent median, IQR, and 1.5x IQR, with outliers marked.

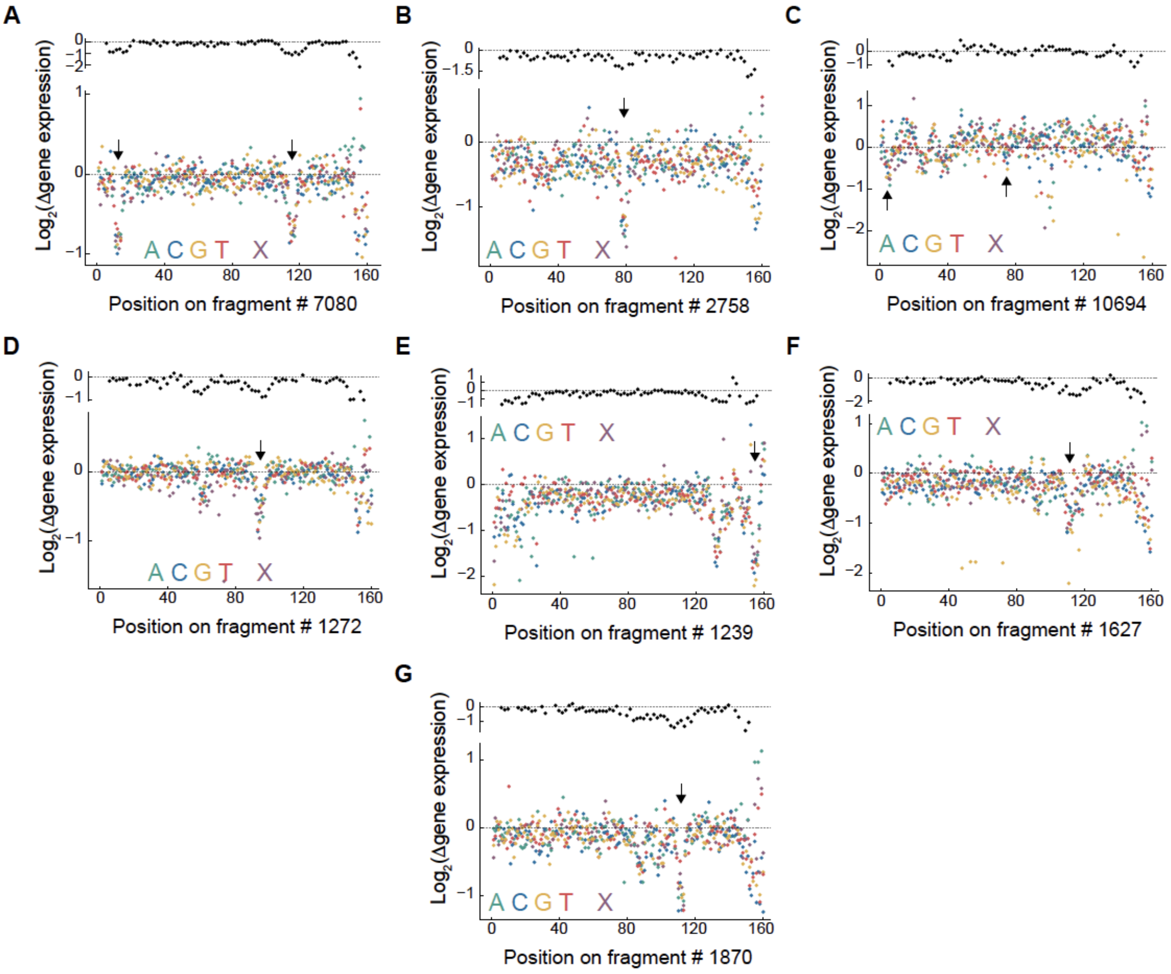


**Supplementary Figure S18: Examples of deep mutational scan of fragments in MPRA in pTRP1-based libraries**

Deep mutational scans of fragments as indicated in the x-axis using pTRP1-based libraries. All examples presented as in Fig. 3E.

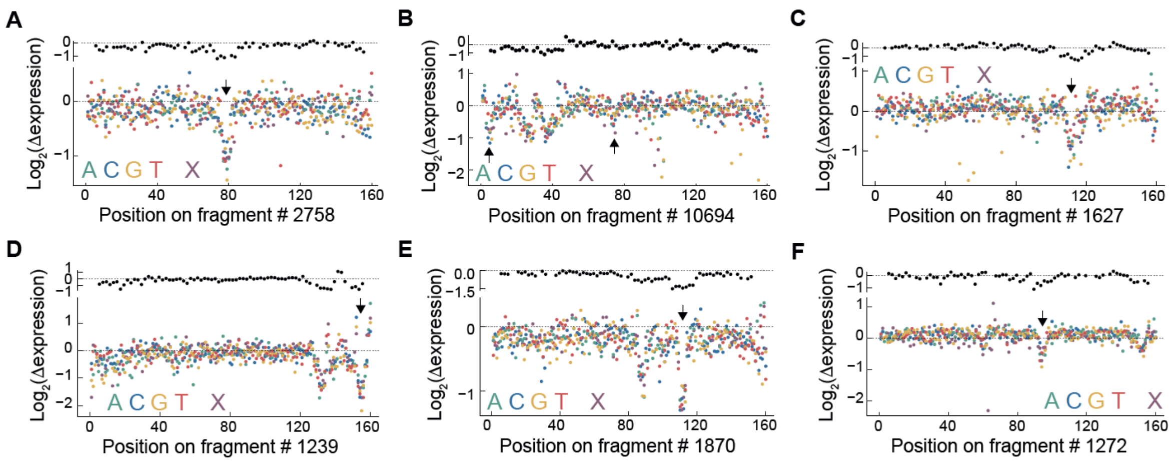


**Supplementary Figure S19: Examples of deep mutational scan of fragments in MPRA in p35S-based libraries**

Deep mutational scan of fragments as indicated in the x-axis using p35S-based libraries. All examples presented as in Fig. 3E for six additional fragments.

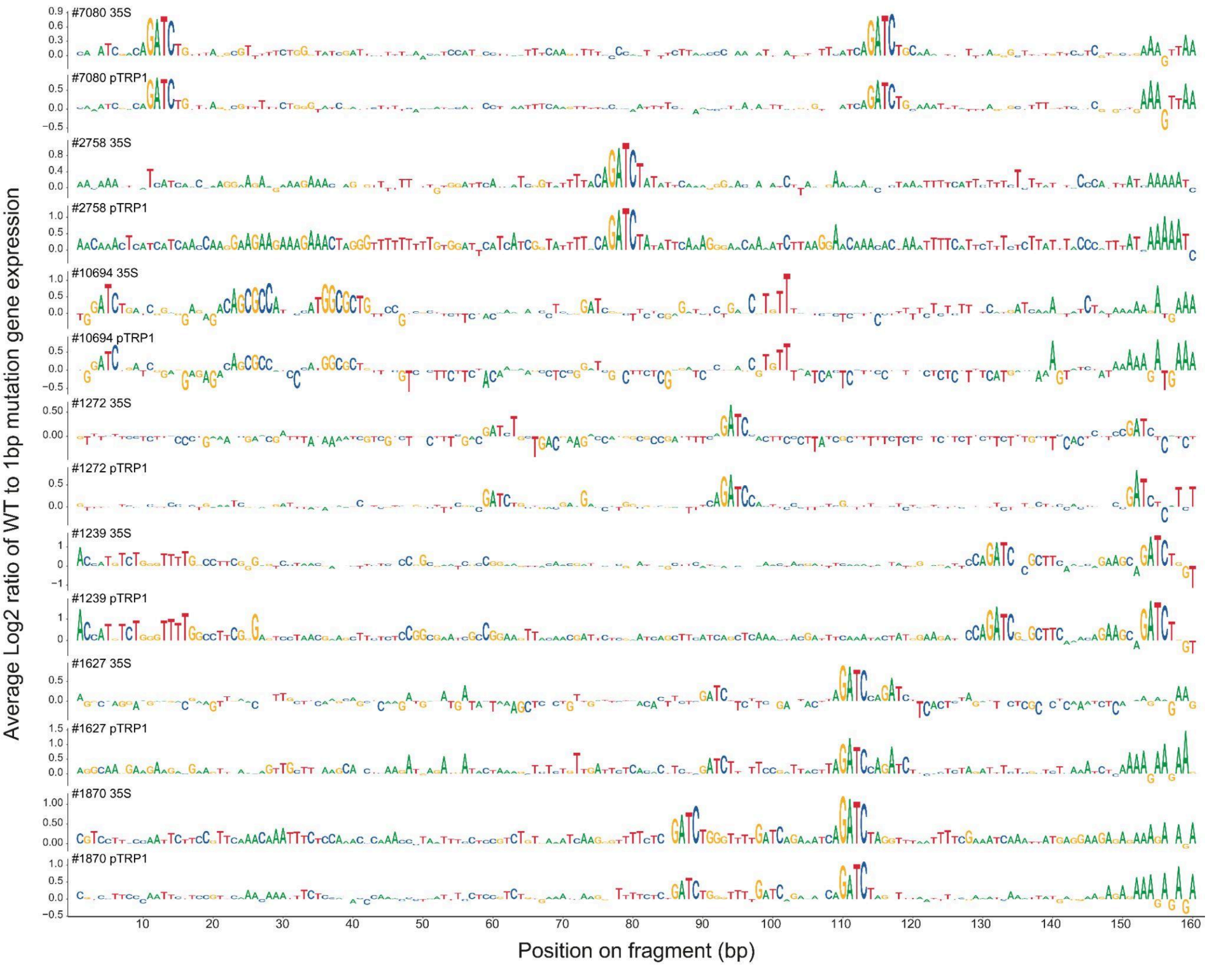


**Supplementary Figure S20: The per-nucleotide contribution to transcriptional regulation calculated from deep mutation scanning**

The impact of mutating each position within the fragments is shown by plotting the average log2 difference in gene expression between variants with the WT nucleotide and all four of their mutated counterparts. For each 1-bp mutation analyzed, we calculate this difference and plot the average change in expression as the height of the corresponding nucleotide in the original fragment. The plots include the fragments shown in Figs. 3E and S18-19. Each plot is labeled with the fragment index and the library core promoter used.

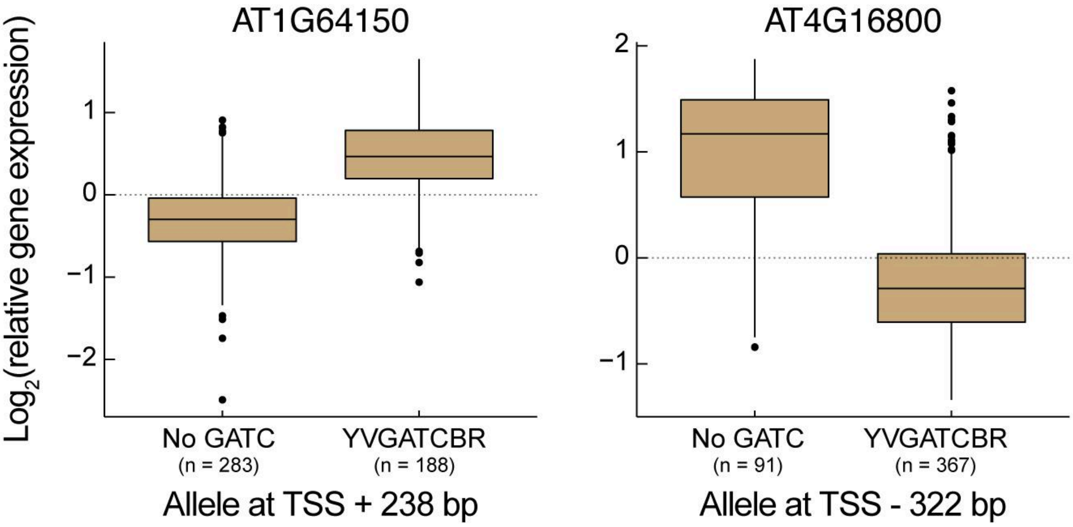


**Supplementary Figure S21: Examples of differential gene expression associations with alleles having or lacking a GATC motif in *A. thaliana* natural accessions**

Two representative examples from the summary statistics in Fig. 3G, illustrating the association between gene expression and the presence of a GATC motif (*1*, *13*). On the left, accessions with different alleles of AT1G64150 are grouped based on the presence of the GATC motif 238 bp downstream of the TSS or the complete absence of the 4 bp GATC sequence. Intermediate cases are omitted. Expression values are shown relative to the mean in the population. On the right, a similar graph is depicted for gene AT4G16800, with the motif 322 bp upstream of the TSS. Boxplots display the median, IQR, and 1.5x IQR, with outliers highlighted.

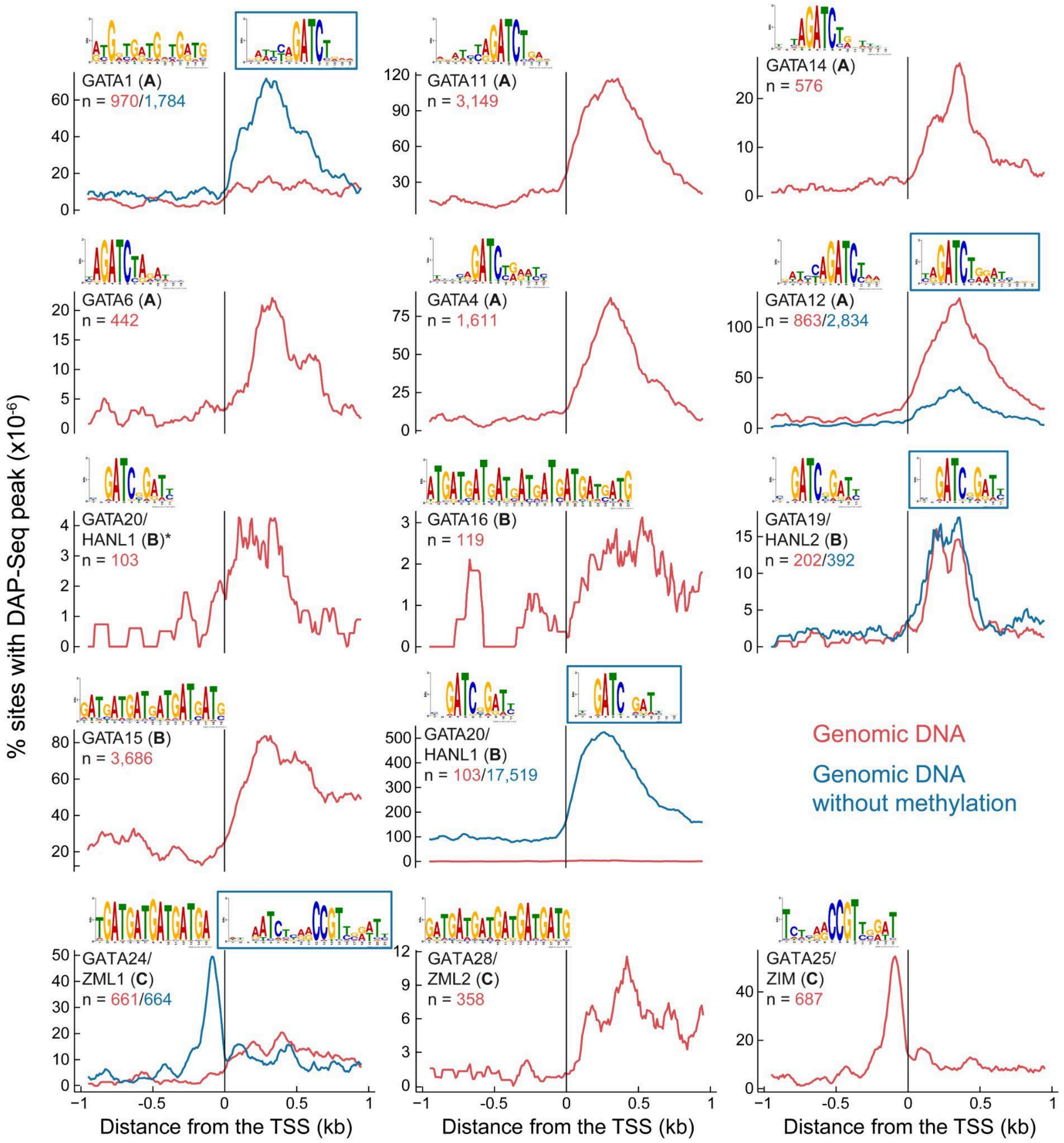


**Supplementary Figure S22: DAP-Seq data of GATA TFs from O'Malley et al.**

Enrichment relative to the TSS of the binding peaks of each of the 13 GATA TFs examined in ref' (*14*). Each GATA TF is plotted in a different subplot as indicated, the subfamily of the GATA is indicated in brackets (*15*, *16*). The number of identified peaks (*n*) is given below the TF name, if the TF was also assayed with non-methylated DNA (blue line), the number of binding peaks for this is also given in blue. Finally, the DNA motif identified in (*14*) is plotted above the graph, a blue frame is added if it was derived from the assay with non-methylated DNA. GATA20 is plotted twice, once with only the methylated DNA, due to the large difference in dynamic range between them. Note that the DNA motif associated with all of subfamily A is similar to the GATC motif identified in this work, while a few TFs in subfamily B also bind a motif with a GATC sequence, it also has a different affinity for “GAT” not inferred in the MPRA results of this work.

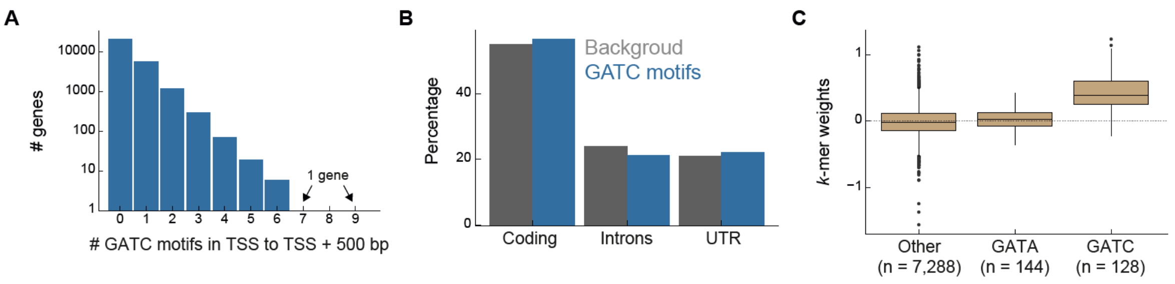


### Supplementary Figure S23: Genomic distribution of the GATC motif and its association with GATA factor binding

**(A)** Distribution of GATC motifs in the 500 bp region downstream of the TSS in Arabidopsis genes. The graph depicts the number of genes with different counts of GATC motifs within this window. **(B)** Distribution of GATC motifs in different genomic contexts within the 500 bp region downstream of the TSS, against the genomic background. **(C)** *k*-mer weights derived from an analysis of GATA12 ChIP-Seq in maize (*17*). A higher weight indicates stronger binding propensity of the transcription factor GATA12 to sequences containing that specific *k*-mer. The weights of *k*-mers containing either GATC or GATA are plotted along weights of all other *k*-mers. The weights serve as a measure of sequence preferences when modeling GATA12 TF binding based on the machine-learning model applied by the authors. The comparison to GATA-containing *k*-mers is highlighted due to the propensity of GATA factors to bind GATA sequences in other species (Supplementary Note S1). Boxplots display median, IQR, and 1.5x IQR with outliers as points, numbers of *k*-mers (*n*) are indicated.

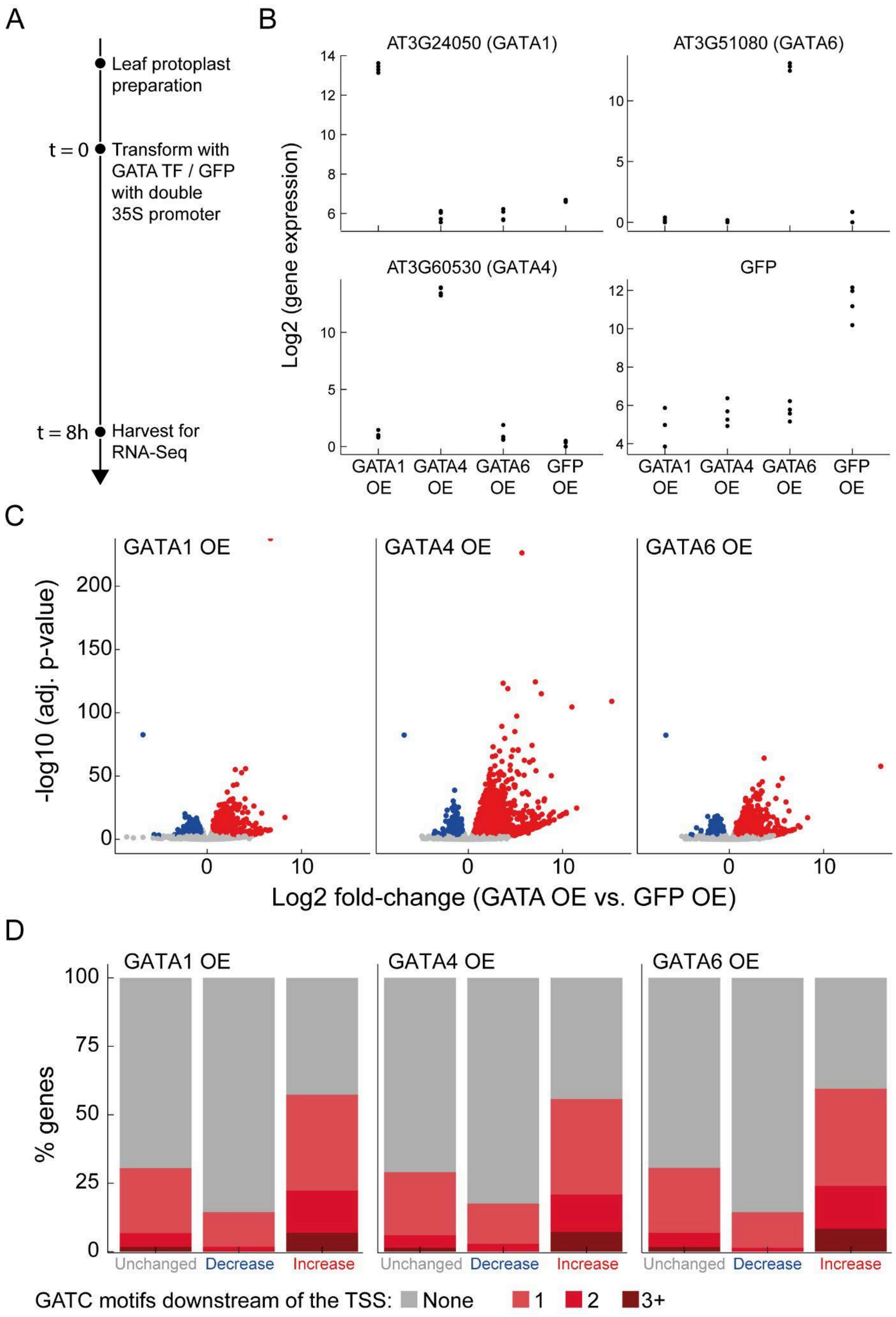
A
Leaf protoplast preparation
t = 0
Transform with GATA TF / GFP with double 35S promoter
t = 8h
Harvest for RNA-Seq
B
AT3G24050 (GATA1)
AT3G51080 (GATA6)
AT3G60530 (GATA4)
GFP
Log2 (gene expression)
GATA1 OE
GATA4 OE
GATA6 OE
GFP OE
C
GATA1 OE
GATA4 OE
GATA6 OE
-log10 (adj. p-value)
Log2 fold-change (GATA OE vs. GFP OE)
D
GATA1 OE
GATA4 OE
GATA6 OE
% genes
Unchanged
Decrease
Increase
GATC motifs downstream of the TSS: None 1 2 3+

### Supplementary Figure S24: Transient overexpression of GATA TFs promotes the expression of genes containing the GATC motif

**(A)** Experimental setup: Arabidopsis leaf protoplasts were isolated and transformed by PEG-mediated delivery with plasmids carrying GFP, GATA1, GATA4, or GATA6 under the control of a double 35S promoter. RNA was extracted for RNA-Seq analysis eight hours after transformation. Two biological repeats of protoplast isolation were done, with each plasmid transformed twice per isolation replicate, resulting in four replicates per plasmid. In each set of protoplast isolation, an additional sample was transformed with the GFP plasmid to estimate transformation efficiency, which was determined by imaging to be 59% and 69.7% at eight hours post-transformation. **(B)** Verification of overexpression (OE): Gene expression levels of GATA1, GATA4, GATA6, and GFP (as indicated in the panel title) are shown. Expression levels are log2-transformed and plotted for the four plasmids across all four replicates. Note the low level of GFP detection in the GATA TF samples is due to leaky expression of a YFP selection gene driven by a seed coat promoter also found in the plasmids; due to sequence similarities between YFP and GFP, there is cross-alignment of sequencing reads. **(C)** Differential expression analysis: A volcano plot shows the -log10 adjusted p-values relative to the log2 fold changes in gene expression following OE of each GATA TF compared to GFP OE. Calculations were performed using DESeq2 based on four replicates per experiment (*18*). Genes showing at least a 50% change in expression with an adjusted p-value < 0.001 were defined to be differentially expressed and are shown in blue (downregulated) and red (upregulated); all other genes are shown in gray. **(D)** The percentage of genes with a GATC motif within 500 bp downstream of the TSS is shown. These are categorized based on increased, decreased, or unchanged expression (labeled "Unchanged") as defined in **(C)**. Further separation is provided for genes with 1, 2, or 3 or more GATC motifs for each GATA TF OE experiment.

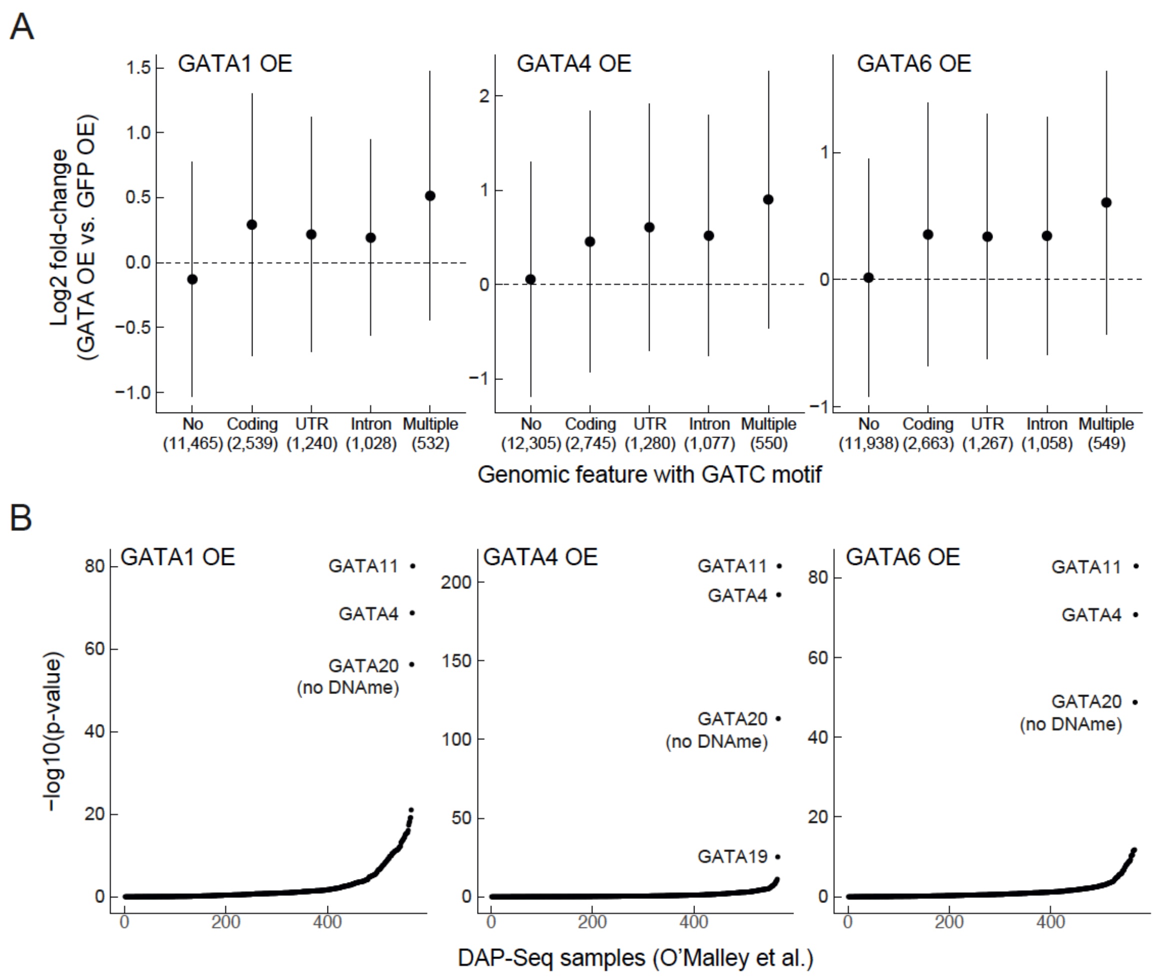


### Supplementary Figure S25: Transient overexpression of GATA TFs affects genes identified as GATA targets by DAP-Seq

**(A)** Shown is the log2-fold change in gene expression in response to GATA TFs OE (as indicated) relative to GFP OE. Data are plotted for genes without GATC motifs in the 500 bp downstream of the TSS, genes with GATC motifs only in coding regions, UTRs, introns, or those with motifs in more than one genomic feature. Error bars represent the mean ± 1 standard deviation; numbers of genes per category are indicated. **(B)** Enrichment analysis was performed to compare gene targets identified by DAP-Seq for all TFs assayed in the (*14*) study with genes upregulated (as defined in Fig. S24C) upon overexpression of the GATA TFs. Enrichment p-values were calculated using a hypergeometric test and are shown for each of the TFs samples previously assayed by DAP-Seq (*14*). For the three GATA overexpression experiments, the genes with increased expression are most enriched with the targets of GATA TFs in the DAP-Seq data.

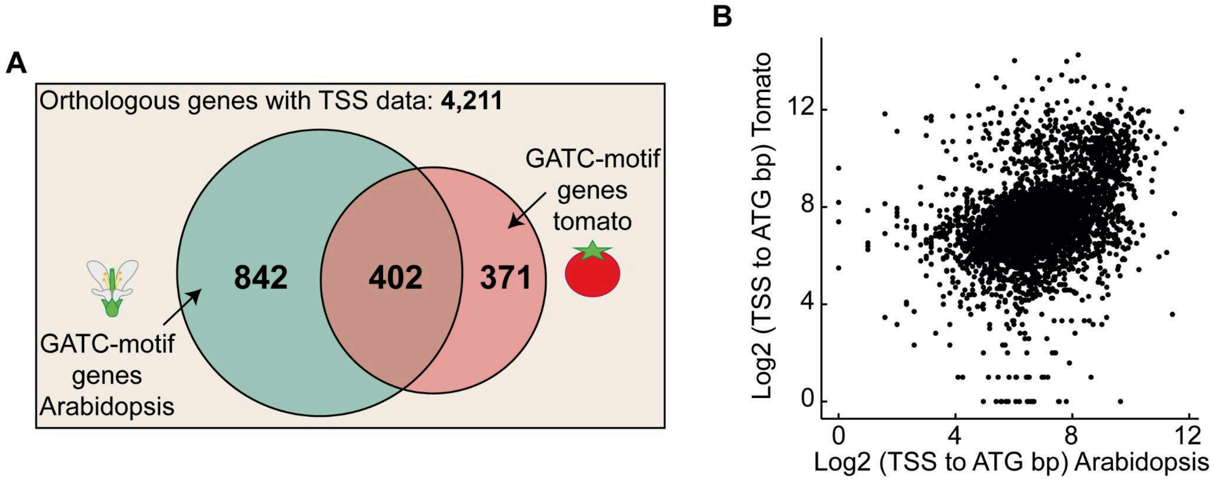


**Supplementary Figure S26: Conservation of GATC-motif genes between Arabidopsis and tomato**

One-to-one orthologous genes between Arabidopsis and tomato were identified using OrthoFinder (*19*). Out of 6,883 one-to-one pairs, 4,211 possessed a TSS upstream of the ATG in both species, forming the basis for comparison. **(A)** The conservation of genes containing a GATC motif within the 500 bp region downstream of the TSS is depicted. The Venn diagram illustrates orthologous genes and those harboring the GATC motif in one or both species. The intersection of genes possessing the motif is statistically significant (p-value $< 10^{-49}$) according to a hypergeometric enrichment test. **(B)** Analysis of the distance between the TSS and the ATG demonstrates conservation between Arabidopsis and tomato, with a Spearman correlation of 0.42 and a Pearson correlation of 0.26. Notably, this level of correlation, though slightly stronger here, parallels findings from an earlier study comparing the 5’ UTR length between *Candida albicans* and *Saccharomyces cerevisiae (20)*.

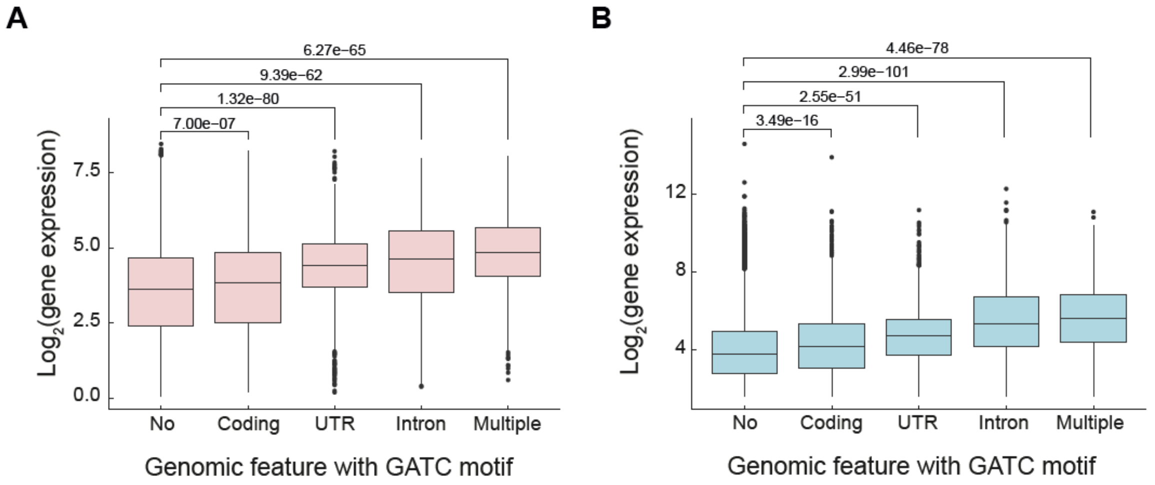


#### Supplementary Figure S27: Association between GATC-motifs presence and gene expression across different genomic contexts

Gene expression levels are displayed for the aerial parts of Arabidopsis seedlings (*21*), as depicted in Fig. 4A **(A)**, and in seedling roots (*22*), corresponding to Fig. 4F **(B)**. Genes are grouped according to the GATC motifs located within the 500 bp region downstream of the TSS, categorized by specific genomic features: no GATC motifs in this region (21,227 genes), motifs exclusively in the coding region (4,003 genes), in UTR regions (1,395 genes), in introns (1,271 genes), and genes with GATC motifs spanning more than one feature type, such as one motif in the coding region and another in a UTR (600 genes). p-values are shown for comparison of gene expression levels for genes with GATC motifs in any of the feature categories versus those without any GATC motif within the 500 bp region downstream of the TSS. Statistical significance was assessed using the Welch t-test.

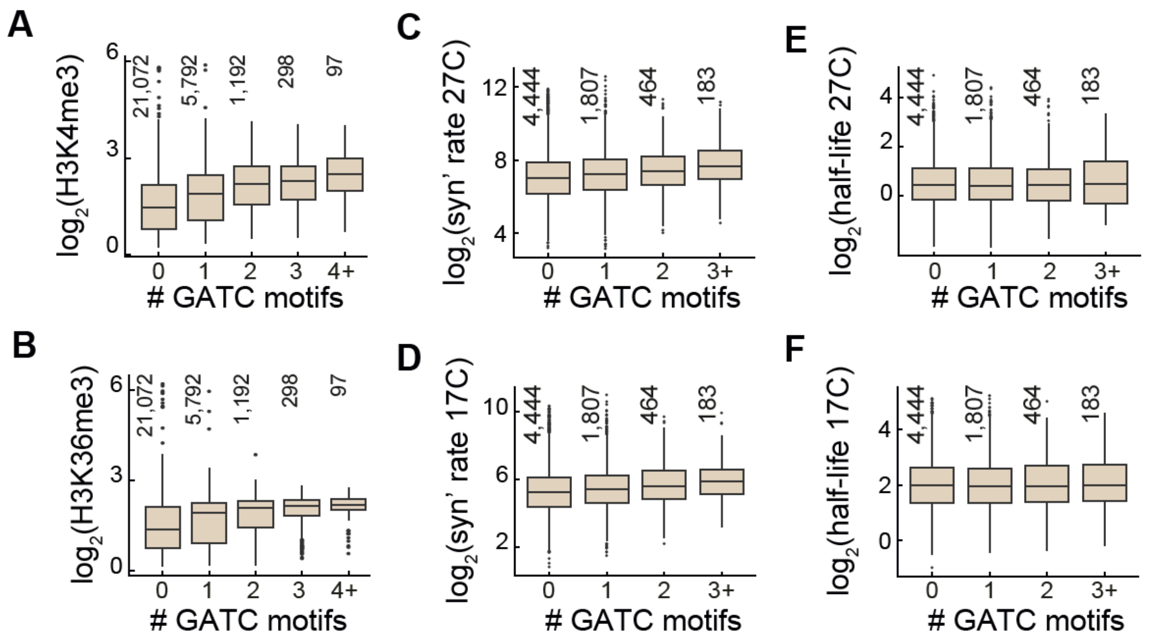


**Supplementary Figure S28: Association between GATC-motifs count and histone modification, mRNA synthesis rate, and mRNA half-life**

**(A-B)** Average $\log_2$ enrichment of H3K4me3 **(A)** and H3K36me3 **(B)** across genes (*4*, *23–25*) categorized by GATC-motif counts within 500 bp downstream of the TSS, as in Fig. 4A. H3K4me3 and H3K36me3 effect sizes are 0.27 (p-value: $3*10^{-300}$) and 0.2 (p-value: $4.4*10^{-194}$), respectively. **(C-D)** mRNA synthesis rates for 7,291 genes at 27°C **(C)** and 17°C **(D)**, plotted by GATC-motif counts (*26*). Genes with motif counts >=3 are grouped. Effect sizes are 0.17 (p-value: $2.9*10^{-16}$) at 17°C and 0.18 (p-value: $8.8*10^{-19}$) at 27°C. **(E-F)** mRNA half-lives at 27°C **(E)** and 17°C **(F)**, plotted similarly to **C-D**. No significant associations were found between GATC-motif and mRNA half-lives (*26*), with effect sizes of 0.01 (p-value: 0.4) at 17°C and -0.01 (p-value: 0.6) at 27°C. Boxplots display median, IQR, and 1.5x IQR with outliers as points. Number of genes per box plot is indicated.

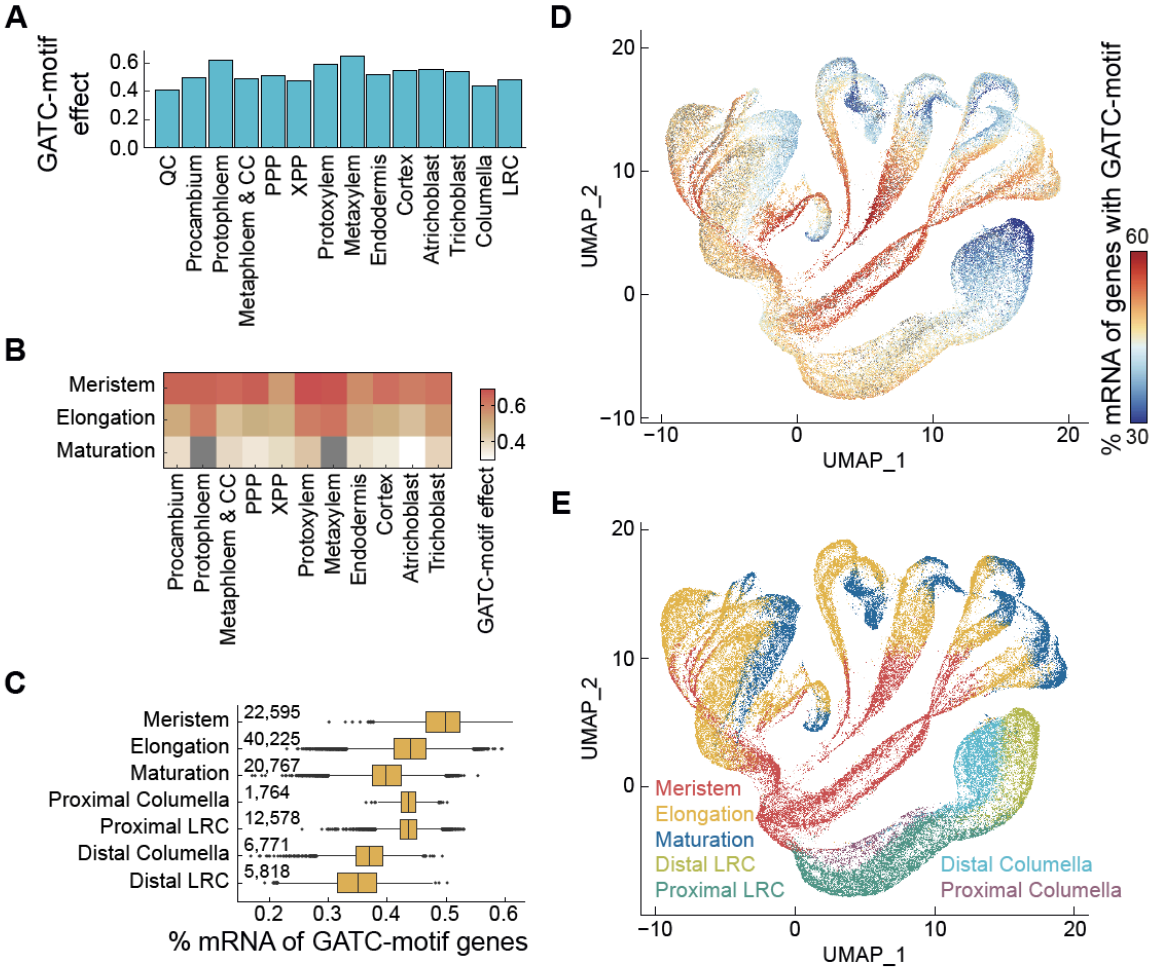


### Supplementary Figure S29: Effect of GATC motif on expression across root cell types

**(A)** GATC-effect sizes across root cell types, as quantified in Fig. 4G. Gene expression was averaged from scRNA-seq data based on published cell annotations (*27*). QC - Quiescent center; LRC - Lateral root cap; CC - companion cell; PPP - phloem pole pericycle; XPP - xylem pole pericycle. **(B)** GATC-effect sizes visualized for different root cell types (x-axis) across root developmental stages (y-axis). Analysis uses cases with at least 150 cells intersecting the two definitions. Gray rectangles indicate cases below this threshold. The quiescent center, defined only as meristematic, is excluded. **(C)** Given that each single cell only provides data on a subset of genes, we used an alternative method to quantify the GATC-motif effect at the single-cell level. For each cell, mRNA counts from genes with a GATC motif were divided by the total mRNA count. This approach is more robust for sparse data. The quantities plotted reveal significant changes in mRNA species across root developmental stages. Boxplots show median, IQR, and 1.5x IQR, with outliers marked. Number of cells per category is indicated. **(D)** The mRNA percentage from GATC-motif-containing genes is depicted for each of the 110,000 cells in the single-cell Arabidopsis root atlas map (*27*) **(E)** Root developmental stage definitions from ref. (*27*), presented as in their original study for comparison.

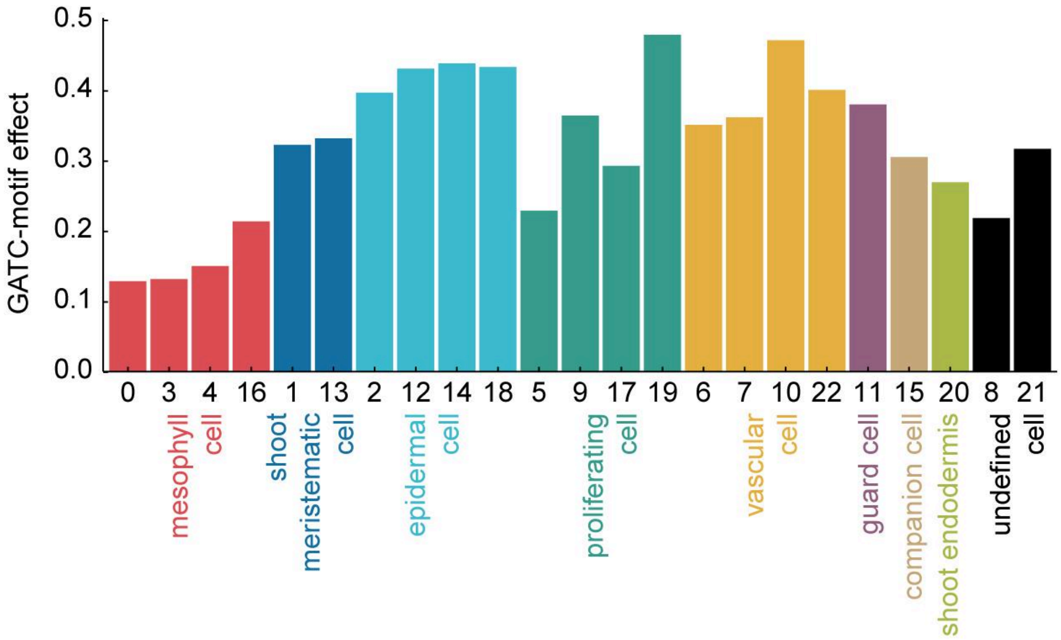


**Supplementary Figure S30: Effect of GATC motif on expression throughout the Arabidopsis vegetative shoot**

GATC-motif effect sizes across different cell types within the Arabidopsis vegetative shoot, as quantified in Fig. 4G. Gene expression was determined by averaging from scRNA-seq data, categorized based on the 23 clusters defined in ref. (*28*). Clusters are annotated as presented in the original study.

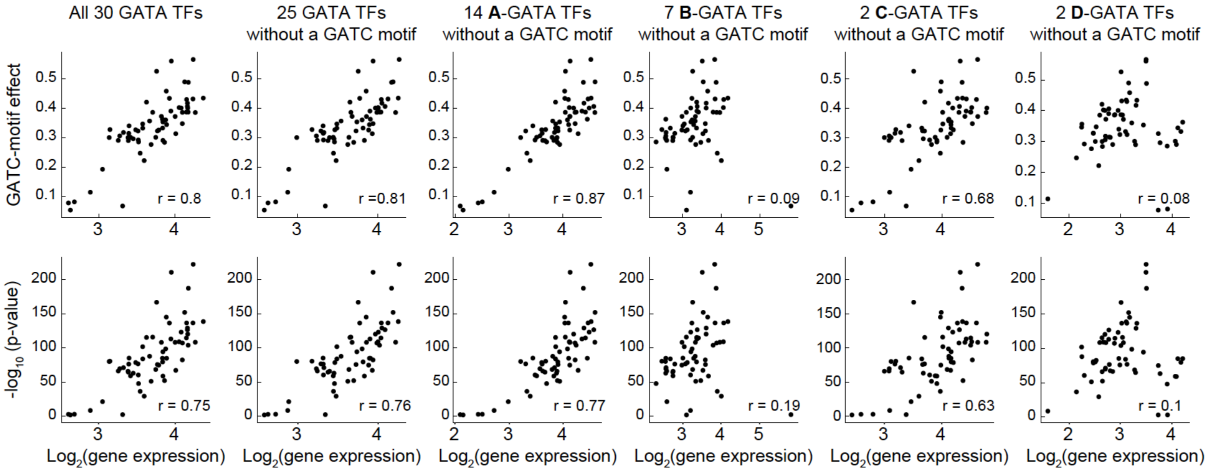


### Supplementary Figure S31: Correlation between GATC-motif effect and expression of GATA transcription factors

Shown is the relationship between the GATC-motif effect (upper row) and the associated fit p-values (lower row), as determined in Fig. 4D, across various tissues part of ref. (*22*) dataset vs. the mean expression levels of GATA TFs within the same tissues. The average expression is calculated across different groups of GATA TFs: all 30 GATA TFs (1st column), the subset of 25 GATA TFs lacking the GATC motif within 500 bp downstream of the TSS (2nd column), and GATA TFs excluding those with the GATC motif and categorized into subfamilies A (3rd column), B (4th column), C (5th column), and D (6th column), according to classifications by (*16*) and (*15*). The number of GATA TFs included in the average for each plot is noted in the column titles. The analysis of GATA TFs lacking the motif serves as a control to ensure that the correlation is not influenced by GATA TFs regulated by the GATC motif themselves. Pearson's correlation coefficients are displayed on each scatter plot (*r*). Notably, DAP-Seq analysis by ref. (*14*) has demonstrated that the A subfamily of GATA TFs binds to the GATC motif downstream of the TSS (Fig. S22).

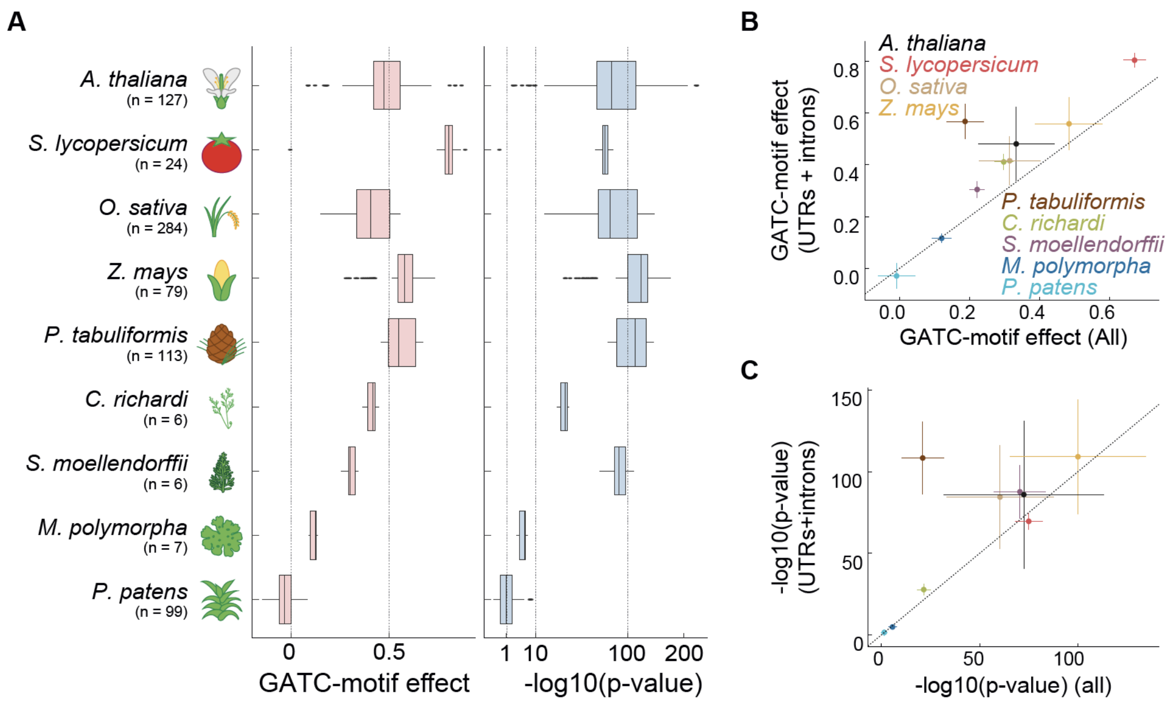


### Supplementary Figure S32: Downstream GATC motifs are associated with higher gene expression when located in introns or UTRs across species

**(A)** Association between GATC-motif downstream of the TSS and gene expression across land plant species is shown as in Fig. 5, counting only motifs found either in introns or UTRs in the 500 bp downstream of the TSS. The effect of the GATC motif on expression (slope, left panel) and the significance of the association (p-value, right panel) are shown. Box plots show median, IQR and 1.5x IQR with outliers. The right panel x-axis is square-root scaled. **(B-C)** Comparison of GATC motif effect size **(B)** and significance of associations **(C)** considering GATC motifs found in all genomic contexts (x-axis) vs. only ones in introns or UTRs (y-axis). Error bars represent one standard deviation around the mean. Points are coloured according to their species, as described in the legend of **(B)**. Dashed lines in **B** and **C** mark y=x.

**Supplementary Note S1 - Indications of plant GATA factors binding to the GATC Sequence**

GATA transcription factors were first studied in erythrocytes (*29*, *30*). Despite their name, derived from their ability to regulate transcription via G-A-T-A DNA sequences, some of these factors often bind non-GATA sequences. Sometimes with even higher affinity than their namesake sequences. Notably, a subset of GATA transcription factors have repeatedly been found to bind the G-A-T-C sequences.

Evidence of this can be seen in various species. Vertebrate GATA-1, GATA-2, and GATA-3 have demonstrated binding to GATC in addition to GATA sequences (*31–33*). In C. elegans, the ELT-1 GATA transcription factor showed a significantly stronger transcriptional activation effect on GATC sequences compared to GATA sequences (*34*). Similarly, in the mushroom *Coprinopsis cinerea*, the CcNsdD2 GATA TF showed a preference for the GATC sequence (*35*). Also in the well studied budding yeast (*S. cerevisiae*) four out of nine GATA TFs are associated with a GATC motif and not a GATA one (*36*).

In 2000, a study by Lowry & Atchley analyzed GATA protein sequences across multiple species (*37*). They found that GATA TFs from *A. thaliana* grouped separately from vertebrate GATAs. Subsequent large-scale studies investigating TF-binding motifs, both cross-kingdom and Arabidopsis-specific, profiled 19 GATA TFs from Arabidopsis (*14*, *38*, *39*). Interestingly, in these studies, none of the Arabidopsis GATA TFs showed binding to a GATA motif, but in most cases rather demonstrated a preference for the GATC sequence (Fig. S22).

Further support for the preference for the GATC sequence emerges from more focused studies in plants. ChIP-Seq analysis of the GNC and CGA1 GATA TFs in Arabidopsis, for instance, showed binding to the GATC motif (*40*). Similarly, ChIP-Seq in maize identified a preference for GATC, rather than GATA sequences, for the GATA12 TF (*17*) (Fig. S23C).

Evidence from other plant species also reflects this trend. In tobacco, the AGP1 GATA TF was found to induce expression by binding a GATC-containing motif (*41*). In *Catharanthus roseus*, the CrGATA1 GATA TF activated five light-responsive vindoline pathway genes via the GATC motif. Intriguingly, removing the GATA sequence did not affect this induced expression (*42*).

Taken together, these findings from both large-scale and gene-specific studies suggest that, in a majority of instances, GATA TFs in plants primarily interact with GATC motifs.

### Experimental Methods:

#### Construction of MPRA backbone plasmids

Plasmids pPSup_iGFP (id: 149420) and pPSint (id: 149421) were obtained from Addgene, and were modified to generate six new plasmids. Initially, BbsI sites were replaced with BsmBI sites and a 1 bp mutation was introduced to eliminate an extra BsmBI site in both plasmids. A ccdB lethal cassette was inserted between the two BsaI sites. These alterations produced pPSup_iGFP_v2 and pPSint_v2. Next, pPSint_v2_rmBsaI was created by removing the BsaI insertion sites with the *ccdB* cassette from pPSint_v2. Next, a 633 bp genomic region was cloned into both pPSup_iGFP_v2 and pPSint_v2. This region was lifted from the TRP1-gene's promoter, extending up to 40 bp upstream of the main transcription start site (TSS). The pTRP1 was positioned upstream of the BsaI site in pPSup_iGFP_v2 and upstream of the BsmBI site in pPSint_v2, generating pPSup_iGFP_v2_pTRP1 and pPSint_v2_pTRP1, respectively. Lastly, the BsaI and ccdB cassette were removed from pPSint_v2_pTRP1 to create pPSint_v2_pTRP1_rmBsaI.

#### Amplification of oligonucleotide pools

Two oligonucleotide pools were obtained from Twist Bioscience, consisting of 12,000 and 17,996 oligonucleotides of length 200 bp, referred to as OP1 and OP2, respectively. The design of these pools is detailed in the "Oligonucleotide pools design" section. Two separate amplification reactions were done for (1) OP1 and (2) an equimolar mix (in the level of single fragments) of OP1 and OP2 (2:3 ratio of full libraries) as follows: 20 reactions were performed, each containing 1 µl of OP1 or OP1+OP2 mix (1 ng/µl), 5 µl of primer P1 (10 µM, primers are listed in table S1), 5 µl of primer P2 (10 µM), 25 µl of KAPA HiFi HotStart ReadyMix (KK2601), and 14 µl of DDW. The reactions were run using the following thermal cycling protocol: 95°C 3', (98°C 20", 60°C 30", 72°C 20") x 15, 72°C 1'. The amplified oligonucleotide pool was then purified using 1.4x AMPure XP (A63880) beads and the amplification was verified with an Agilent Fragment Analyzer.

#### Cloning oligonucleotide pools and barcodes into plasmid backbones

Cloning of OP1 into pPSup_iGFP_v2, pPSint_v2, pPSup_iGFP_v2_pTRP1, or pPSint_v2_pTRP1, as well as the OP1+OP2 mixture into pPSint_v2 or pPSint_v2_pTRP1, was performed using 20 Golden-Gate reactions. Each reaction contained 1 µl of backbone plasmid (75 ng/µl), 1 µl of the amplified oligonucleotide pool (5 ng/µl), 2.5 µl of T4 DNA ligase (M0202T), 1.5 µl of BsaI-HFv2 (R3733S), and 18.5 µl of DDW. A control reaction was also carried out, with DDW replacing the oligonucleotide pool. The reactions were incubated in a thermal cycler using the following protocol: (37°C 5', 16°C 5') x 30, 60°C 5'. The reactions were then cleaned with 1.5x AMPure XP beads. The resulting reactions were transformed into MegaX DH10B T1R Electrocomp™ Cells (C640003) *E. coli*, according to ref. (*43*). A dilution series was plated on LB+Spectinomycin for both reaction and control in order to estimate cloning complexity. The rest of the transformation was grown overnight and purified with QIAGEN Plasmid Plus Midi Kit (12943).

A second set of Golden-Gate reactions was performed with the libraries containing the cloned oligonucleotide pools to add minimal promoter, 5' UTR as well as a 15 bp barcode (VNN x 5). Two different inserts were used. The first insert contained a CaMV 35S minimal promoter and the SynJ synthetic 5' UTR, as described (*10*, *11*), amplified using P3-P5 primers, diluted to 3.13 ng/µl, and inserted into pPSup_iGFP_v2, pPSint_v2, or pPSint_v2_rmBsaI. A second insert contained the minimal promoter and 5' UTR of the *TRP1* gene, amplified using P6-P8 primers, diluted to 5 ng/µl, and inserted into pPSup_iGFP_v2_pTRP1, pPSint_v2_pTRP1, or pPSint_v2_pTRP1_rmBsaI. 15 Golden-Gate reactions and 1 control were done for pPSup_iGFP_v2, pPSint_v2, pPSup_iGFP_v2_pTRP1, and pPSint_v2_pTRP1 as described above substituting the restriction enzyme with BsmBI-v2 (R0739L) and adding the corresponding insert to each reaction, with the thermocycler protocol: (42°C 5', 16°C 5') x 30, 60°C 5'. Transformation into *E. coli* and efficiency calculation followed the same procedure. For pPSint_v2_rmBsaI and pPSint_v2_pTRP1_rmBsaI, two modifications were made: only 10 Golden-Gate reactions and 1 control were performed, and in-house prepared *E. coli* competent cells were used.

This procedure resulted in 8 plasmid libraries: (1) pPSup_iGFP_v2 (p35S.SynJ-OP1), (2) pPSint_v2 (p35S.SynJ-OP1), (3) pPSint_v2_rmBsaI (p35S.SynJ), (4) pPSup_iGFP_v2_pTRP1 (pTRP1-OP1), (5) pPSint_v2_pTRP1 (pTRP1-OP1), (6) pPSint_v2_pTRP1_rmBsaI (pTRP1), (7) pPSint_v2 (p35S.SynJ-OP1+OP2), and (8) pPSint_v2_pTRP1 (pTRP1-OP1+OP2). Fig. S7 depicts the structure of the libraries.

**Mixing of input MPRA libraries**

Three mixes of libraries were made. The labeled MIX1 contained 6 libraries pPSup_iGFP_v2 (p35S.SynJ-OP1), pPSint_v2 (p35S.SynJ-OP1), pPSup_iGFP_v2_pTRP1 (pTRP1-OP1), pPSint_v2_pTRP1 (pTRP1-OP1), pPSint_v2_pTRP1_rmBsaI (pTRP1), and pPSint_v2_rmBsaI (p35S.SynJ) in 50:50:50:50:1:1 proportions, respectively. The second, labeled MIX2, with 4 libraries pPSint_v2 (p35S.SynJ-OP1+OP2), pPSint_v2_pTRP1 (pTRP1-OP1+OP2), pPSint_v2_rmBsaI (p35S.SynJ), and pPSint_v2_pTRP1_rmBsaI (pTRP1) in 100:100:1:1 proportions, respectively. The third, labeled MIX3, contained 2 libraries pPSint_v2_pTRP1 (pTRP1-OP1) and pPSint_v2_pTRP1_rmBsaI (pTRP1) in 100:1 proportions, respectively. These three mixes were transformed into MegaX DH10B T1R Electrocomp™ Cells, with efficiency of $>10^8$, and then purified with QIAGEN Plasmid Plus Giga Kit (12191).

**Sequencing of input MPRA libraries**

In order to connect barcodes to tested fragments and libraries, the relevant region from the cloned libraries was sequenced with next generation sequencing (NGS). To avoid the same initial bases in reads 1 and 2 in the Illumina run, due to amplification using the constant sequence around the barcode and enhancer, which is detrimental to the imaging analysis of sequence signal, primers were chosen to create variation in read start. Every forward or reverse primer used was a combination of 4 primers that each had a different, 0-3 nucleotide of shift before the constant sequence. Primers P9-12 & P13-16 were used for pPSup_iGFP_v2 (p35S.SynJ-OP1), P17-P20 & P21-24 used for pPSint_v2 (p35S.SynJ-OP1) and pPSint_v2 (p35S.SynJ-OP1+OP2), P17-P20 & P25-28 used for pPSint_v2_rmBsaI (p35S.SynJ), P13-16 & P29-32 used for pPSup_iGFP_v2_pTRP1 (pTRP1-OP1), P33-36 & P21-24 used for pPSint_v2_pTRP1 (pTRP1-OP1) and pPSint_v2_pTRP1 (pTRP1-OP1+OP2), and P33-36 & P25-28 used for pPSint_v2_pTRP1_rmBsaI (pTRP1). In order to estimate exact mixes' ratios, additional shotgun Tn5-based DNA-Seq libraries were prepared from each of the library mixes and sequenced in paired-end mode on an Illumina NovaSeq 6000 machine.

**Plant material and growth conditions**

*Arabidopsis thaliana* Col-0 seeds were sterilized with 70% [v/v] ethanol and 6.5% [v/v] bleach and sown on circular plates containing ½ MS, 0.05% [w/v] MES and 0.8% [w/v] agar. The plants were grown under long-day conditions (21°C, 85 μmol/m$^2$/s) for 23-26 days. Tomato (*S. lycopersicum*) cv. M82 ($sp^-/sp^-$) seeds were sterilized with 70% ethanol and 3.25% bleach and sown in Magenta boxes (6 cm x 6 cm x 9.5 cm) containing 62.5 ml 0.217% [w/v] Nitsch medium (Duchefa Biochemie, N0224.0050),

2% [w/v] sucrose and 0.9% agar. Tomato plants were grown under long-day conditions (21°C, 85 μmol/$m^2$/s) for 20 days. Maize (*Z. mays*) cv. B73 seeds were grown in soil (4:1 Klasmann Substrat 2:Perlite) in the greenhouse (long-day photoperiod, 23°C, 150 μmol/$m^2$/s) until 1-2 cm shoots were visible. The pots were covered and grown in the dark for 7-9 days. *Nicotiana benthamiana* cv. LAB seeds were sown on soil (4:1 Gramoflor 2006:Perlite), stratified for 4 days in the dark (4°C) and transferred to short-day conditions (21°C, 60% humidity) and grown for 23 days.

**MPRA assay in tomato and Arabidopsis protoplasts**

The protocol was adapted from ref. (*44*). Briefly, 5-6 leaves from 30-35 Arabidopsis seedlings or the true leaves from 20-25 tomato seedlings were cut into 0.5-1 mm wide strips using a razor blade and immediately submerged in 15 mL enzyme solution (0.4 M Mannitol, 20 mM MES, pH 5.7, 20 mM KCl, 1.5% [w/v] Cellulase R-10 (Duchefa Biochemie, C8001.0010), 0.4% [w/v] Macerozyme R-10 (Duchefa Biochemie, M8002.0005), 10 mM $CaCl_2$, 0.1% [w/v] BSA). The enzyme solution was incubated overnight at 25°C in the dark with gentle agitation (25 rpm). The solution was strained through a 100 μm filter and centrifuged at 100xg for 10 minutes at room temperature. This and all other centrifugation steps on protoplasts were performed with a soft start and end. The pellet was resuspended in 3 mL W5 solution (154 mM NaCl, 125 mM $CaCl_2$, 5 mM KCl, 2 mM MES, pH 5.7) and healthy protoplasts were isolated using a sucrose gradient (23% [w/v]) by centrifugation at 450xg for 3 minutes at room temperature. The protoplast fraction was resuspended in 14 mL W5 solution and counted on a hemocytometer. The suspension was centrifuged at 100xg for 10 minutes at room temperature and the pellet was resuspended in MMG solution (0.4 M Mannitol, 15 mM $MgCl_2$, 4 mM MES, pH 5.7) to a concentration of 1 million cells/mL.

From the suspension, two separate samples of 200 μL were taken, each containing 200,000 protoplasts. These were placed into two distinct 1.5 mL tubes. To the first tube, 10 μg of a plasmid, which codes for the Clover protein with a Nuclear localization signals (NLS) sequence expressed with the pUBI promoter, was added as a positive control. To the second tube, 10 μL of elution buffer was added, serving as the negative control. The rest of the suspension was split into 50 mL falcon tubes with 4-6 mL suspension each and mixed with 50 μg/million cells plasmid library. An equal volume of PEG solution (0.2 M Mannitol, 0.1 M $CaCl_2$, 40% [w/v] poly-ethylene glycol MW 4000) to the protoplast/DNA suspension was added to each tube and the protoplasts were incubated for 20 minutes in the dark at room temperature. 0.95 mL W5 solution was added to the controls and 4.75 mL/million cells to the samples. After 15 minutes incubation at room temperature, the protoplast were centrifuged at 450xg for 5 minutes at room temperature and the pellets were resuspended in 1 ml W1 solution (0.5 M Mannitol, 20 mM KCl, 4 mM MES, pH 5.7) for the controls and 5 ml/million cells for the samples. Each protoplast suspension was transferred to a separate sterile petri dish and incubated at 25°C under constant light (85 µmol/$m^2$/s) for 6 h (samples) or overnight (controls). After 6 h, the samples were centrifuged at 450xg for 5 minutes at room temperature and the pellets were flash-frozen in liquid nitrogen. The samples were stored at -70°C until RNA extraction. The controls were imaged under a microscope to check the transformation efficiency - typically around 70-80% in the positive control were successfully transformed both for Arabidopsis and tomato. Testing efficiency of transformation on a large scale, like done for the libraries, gave 60% efficiency. This experiment was done in four replicates for Arabidopsis, yielding 10, 14, 10 and 14 million protoplasts, and in three replicates for tomato, yielding 9, 24 and 16 million protoplasts.

**MPRA assay in maize protoplasts**

The protocol was adapted from ref. (*45*). The middle parts (6-8 cm) of the second leaf from 30 maize plants were used. Each leaf was cut in half and both halves were placed on top of each other, followed by cutting into 0.5-1 mm wide strips perpendicular to the veins using a razor blade. Strips were immediately submerged in 60 mL enzyme solution (0.6 M Mannitol, 10 mM MES, pH 5.7, 1.5% [w/v] Cellulase R-10 (Duchefa Biochemie, C8001.0010), 0.3% [w/v] Macerozyme R-10 (Duchefa Biochemie, M8002.0005), 1 mM $CaCl_2$, 0.1% [w/v] BSA, 5 mM β-mercaptoethanol) split into 4 petri dishes with 15 mL solution each. The enzyme solutions were covered and vacuum infiltrated for 1 h at room temperature and then incubated for 2 h in the dark at room temperature with gentle agitation (40 rpm). The protoplasts were released by shaking at 80 rpm for 10 minutes. The solutions were strained through a 100 µm filter and combined in two tubes with 30 mL solution each. The tubes were centrifuged (all centrifugations with protoplasts were done with soft start/end) at 70xg for 3 minutes at room temperature and the pellets were resuspended in 10 mL 0.6 M Mannitol. The two protoplast suspensions were combined in one tube and counted on a hemocytometer. The suspension was centrifuged at 70xg for 3 minutes at room temperature and the pellet was resuspended in MMG solution (0.6 M Mannitol, 15 mM $MgCl_2$, 4 mM MES, pH 5.7) to a concentration of 1 million cells/mL.

The protoplast suspension was split into several 50 mL falcon tubes with 4-6 mL suspension each and mixed with 200 µg/million cells plasmid library. An equal volume of PEG solution (0.6 M Mannitol, 0.1 M $CaCl_2$, 40% [w/v] poly-ethylene glycol MW 4000) to the protoplast/DNA suspension was added to each tube and the protoplasts were incubated for 15 minutes in the dark at room temperature. 5 mL/million cells W5 solution (same as for Arabidopsis and tomato protoplasts) was added. The samples were centrifuged at 70xg for 3 minutes at room temperature and the pellets were resuspended in 5 ml/million cells incubation solution (0.6 M Mannitol, 4 mM KCl, 4 mM MES, pH 5.7). Each suspension was transferred to a separate sterile petri dish and incubated at 25°C in the dark for 12 h. After 12 h, 1 mL aliquot was saved for imaging and the rest of the samples were centrifuged at 70xg for 3 minutes at room temperature. The pellets were flash-frozen in liquid nitrogen and stored at -70°C until RNA extraction. The aliquot was imaged under a microscope to check the transformation efficiency: 77% and 93% protoplasts were successfully transformed in the first and second replicate, respectively, while the third replicate was not quantified. The protoplast counts for the experiments were: 21, 18 and 20 million.

**MPRA assay in *N. benthamiana***

Libraries MIX1 were transformed into *Agrobacterium tumefaciens* (ACC-110, GV3101 (pSoup), Lifeasible). In a cold room, 25 µL of ACC-110 competent cells were mixed with 1 µL of plasmid, transferred to a pre-chilled 0.1 cm cuvette (BioRad), and electroporated (1,800 V, 25 µF, 200 Ω). Immediately after, 1 mL of pre-warmed (30°C) LB media was added and the mixture was incubated at 30°C for 160 minutes at 200 RPM. Agrobacterium was then plated in a dilution series on LB plates containing Spectinomycin, Gentamicin, and Rifampicin to estimate transformation efficiency. The remaining Agrobacterium were grown overnight in 50 mL LB with the same antibiotics. The next day, Agrobacterium was centrifuged for 5 minutes at 3,000xg, resuspended to an OD600 of 0.5 in infiltration media (50 mM $NaH_2PO_4$, 2 mM MES, 0.5% [w/v] glucose, 150 µM acetosyringone, pH 5.6-5.7), and incubated for 1-2 h in the dark at room temperature with gentle agitation. Agrobacterium was then infiltrated into 17-20 plants (two leaves per plant, using a needless syringe). Plants were not watered for two days prior to infiltration and were watered immediately after. To increase humidity, plants were covered with a transparent cover for 24 h, and infiltrated leaves were harvested in liquid nitrogen 48 h post-infiltration. This experiment was done in four replicates.

**RNA extraction from protoplasts**

Total RNA from Arabidopsis, tomato and maize protoplasts was extracted using the Monarch® Total RNA Miniprep Kit (New England Biolabs, T2010S). The frozen pellets were thawed shortly on ice and then the protocol for "Cultured Mammalian Cells" in part 1 of the kit manual was followed by resuspending each sample with 400-600 µL Lysis buffer and proceeding to part 2. The Arabidopsis and tomato samples were treated with DNase I as recommended in the kit manual. Due to the larger amount of plasmid used for transformation of maize protoplasts, DNase I treatment was repeated three times for maize samples.

**RNA extraction from *N. benthamiana* tissue**

Harvested samples were ground using a mortar and pestle in liquid nitrogen, then mixed with TRIzol™ LS (up to a 1:3 tissue-to-TRIzol™ volume ratio), vortexed for 10 minutes at room temperature, and centrifuged for 5 minutes at 12,000xg and 4°C. The clear fraction was transferred to a new tube, and 0.2 mL of chloroform per 1 mL of TRIzol™ LS was added for lysis. Samples were mixed by shaking for 15 seconds, incubated for 3 minutes at room temperature and then centrifuged for 15 minutes at 16,000xg and 4°C. The aqueous phase was combined (1:1) with chloroform, shaken, incubated for 3 minutes, and centrifuged for 15 minutes at 16,000xg and 4°C. The resulting aqueous phase was mixed with 0.5 mL of isopropanol per 1 mL of TRIzol™ used for lysis, vortexed, incubated for 10 minutes on ice, and centrifuged for 60 minutes at 21,000xg and 4°C. The supernatant was discarded, and the sample was washed twice with 70% ethanol (1:1 ratio of TRIzol™ used for lysis), vortexed, and centrifuged for 5 minutes at 7,500xg and 4°C. Ethanol was removed and residual ethanol was let to evaporate for 7 minutes at room temperature. Finally, RNA was eluted by adding 500 µL DDW and incubating for 10 minutes at 42°C.

**mRNA/poly(A)+ RNA isolation with Dynabeads Oligo (dT)$_{25}$**

mRNA was isolated from total RNA of protoplasts and *N. benthamiana* tissue with Dynabeads Oligo (dT)$_{25}$ (ThermoFisher Scientific, 61005) following the STARR-Seq protocol of ref. (*43*). Briefly, total RNA was heated to 65°C for 7 minutes, followed by incubation on ice for 3 minutes and at room temperature for 1 minute. Two volumes of beads were used for each volume of total RNA. For preparation, the beads were placed on a magnetic separator and washed twice with the same volume of 2x binding buffer (*43*), followed by resuspension in ½ volume of 2x binding buffer. Total RNA was mixed with the washed beads and incubated for 10 minutes on a rolling shaker. The tubes were placed on a magnetic separator and washed twice with the same volume of washing buffer as the starting volume of the beads. For mRNA elution, the beads were resuspended in 60 µL 10 mM Tris·HCl (pH 7.5) and incubated at 80°C for 3 minutes at 750 rpm. The tubes were placed immediately on a magnetic separator and incubated for >1 minute. The eluted mRNA was transferred to a new RNase-free tube. The beads were re-eluted with new 30 µL 10 mM Tris·Cl (pH 7.5) buffer and the two eluates were pooled together.

**MPRA libraries construction and sequencing**

The protocol was adapted from ref. (*43*) and ref. (*10*). From each replicate, ten reactions with 11 µL mRNA each and a construct-specific primer (P37-P40) were prepared for cDNA synthesis using SuperScript IV reverse transcriptase (Thermo Fisher Scientific, 18090010). One of the reactions was used as a no reverse transcription control, in which the RT enzyme was replaced with RNAse-free water. After the cDNA synthesis, 1 µL RNase A (200 µg/mL) was added to each reaction and the samples were incubated at 37°C for 1 h. The nine RT reactions were pooled and purified with 1.8 volumes of AMPure XP beads (Beckman Coulter, A63880) to each volume of cDNA. The no RT control was processed separately in the same way as the RT reactions. In the final step, the RT reactions and the no RT control were eluted in 146 µL and 49 µL 10 mM Tris·HCl (pH 8) buffer, respectively. The purified cDNA was split in two 72 µL aliquots and each aliquot was used for preparation of three PCR reactions amplifying the p35S-based transcripts (P17-20 + P41) and pTRP1-based transcripts (P33-36 + P41), respectively. The no RT control was also split in two 24 µL aliquots and each aliquot was used for one PCR reaction of the p35S and pTRP1 transcripts, respectively. All PCR reactions were prepared with 24 µL template, 25 µL KAPA HiFi HotStart ReadyMix (Roche Molecular Systems, KK2601), 0.5 µL forward primer (100 µM) and 0.5 µL reverse primer (100 µM). The samples from *N. benthamiana* tissue were amplified with 21 cycles, whereas the ones from Arabidopsis, tomato and maize protoplasts - with 24 cycles. The three reactions from each library (p35S and pTRP1) were pooled and purified with an equal volume of AMPure XP beads. The two no RT reactions (p35S and pTRP1) were processed separately in the same way. Finally, all samples were analyzed with 5200

Fragment Analyzer (Agilent, M5310AA) and sequenced on an Illumina NovaSeq 6000 or NextSeq 2000 in paired-end configuration.

**mRNA synthesis rate measurements in MPRA assay**

The protocol was adapted from ref. (*12*) and Click-iT® Nascent RNA Capture Kit from the Invitrogen™ manual. Arabidopsis protoplasts were transformed with MIX3 following the same procedure as for the MPRA in Arabidopsis protoplasts. After transformation, the samples were incubated for 5h and 40 minutes at 25°C under constant light (85 µmol/m$^2$/s). 200 mM 5-ethynyl uridine (5-EU) stock solution (Click-iT™ Nascent RNA Capture Kit, Invitrogen™, C10365) was added to each sample to a final concentration of 200 µM and the samples were incubated for additional 20 minutes before harvesting. Total RNA was extracted from the frozen pellets as described above. DNase I treatment was repeated three times for each sample. mRNA was isolated from the total RNA as described above. In the elution step, the beads (provided in the kit) were resuspended in 55 µL 10 mM Tris·HCl (pH 7.5) and then re-eluted with the first eluate. 5-8 µL of mRNA was set aside for preparation of libraries from total mRNA, whereas the remaining mRNA was split in three aliquots and each aliquot was used for one Click reaction with 0.25 mM biotin azide following the Click-iT™ kit manual. Biotinylated mRNA was precipitated from each Click reaction following the manual and resuspended in 25 µL RNase-free water. 3 µL bead suspension was added to each aliquot of biotinylated mRNA and incubated for 30 minutes at room temperature rotating at 30 rpm. After incubation, the three aliquots were pooled and washed five times with wash buffer 1 and five times with wash buffer 2 (wash buffers from the Click-iT™ kit). The bead suspension was resuspended in 50 µL wash buffer 2 and used immediately for cDNA synthesis.

Libraries were prepared from the total mRNA and 5-EU-labeled mRNA bead suspension samples following the procedure for MPRA libraries with a few modifications. The total mRNA samples were diluted to 50 uL final volume with RNase-free water. From each sample, four reactions with 11 µL mRNA each and a construct-specific primer (P37-P40) were prepared for cDNA synthesis using SuperScript IV reverse transcriptase (Thermo Fisher Scientific, 18090010). An additional reaction with the remaining mRNA (5.5 uL) was prepared as a no reverse transcription control, in which the RT enzyme was replaced with RNAse-free water. The cDNA reactions with 5-EU-labeled mRNA were incubated on a shaker at 1500 rpm to prevent settling of the beads on the bottom of the tube, whereas the reactions with total mRNA were incubated without mixing. Following the final step of cDNA synthesis, the reactions with 5-EU-labeled mRNA were immediately put on a magnetic rack and the supernatants were transferred to new tubes. After this step, the total mRNA and 5-EU-labeled mRNA samples were handled identically. RNase A treatment and purification with AMPure XP beads was performed as described above. In the final step, the RT reactions and the no RT control were eluted in 73 µL and 24.5 µL 10 mM Tris·HCl (pH 8) buffer, respectively. The purified cDNA from the RT reactions was used for preparation of three PCR reactions amplifying the pTRP1-based transcripts (P33-36 + P41). The no RT control was used for one PCR reaction of the pTRP1 transcripts. All PCR reactions were incubated for 24 cycles. The second purification with AMPure XP beads was performed as described above. Finally, all samples were analyzed with 5200 Fragment Analyzer (Agilent, M5310AA) and sequenced on an Illumina NovaSeq 6000 or NextSeq 2000 in a paired-end configuration. This experiment was done in two replicates, yielding 19 and 15 million protoplasts.

To estimate the specificity of the Click-iT™ Nascent RNA Capture Kit in our system, Arabidopsis protoplasts were transformed with a plasmid encoding the Clover protein under the control of UBI4 promoter. In the final step, the protoplast suspension was split into three samples of 4.3 million protoplasts and the samples were incubated at 25°C under constant light (85 µmol/m$^2$/s) for 6 h. After 4 h, 200 mM 5-EU stock solution was added to one of the samples ("2h 5-EU") to a final concentration of 200 µM. After 5h and 40 minutes, the same amount of 5-EU stock solution was added to the second sample ("20 min 5-EU") and an identical volume of DMSO was added to the third sample ("No 5-EU") which served as a negative control. After 6h incubation, all three samples were harvested and stored at -70°C. Total RNA was extracted from the samples as described above. 5-EU-labeled RNA was isolated from the total RNA as described above. 0.5 mM biotin azide was used for each Click reaction. cDNA was prepared from both the total RNA and 5-EU-labeled RNA samples using SuperScript™ VILO™ cDNA Synthesis Kit (Invitrogen™, 11754050). No reverse transcription controls were prepared for each sample, in which the enzyme mix was heat-inactivated at 65°C for 10 minutes following the kit manual. qPCR reactions were prepared from all samples using an in-house qPCR mix and oligonucleotides specific for *ACTIN2* mRNA and *Clover* mRNA. The results were analyzed using Lightcycler® 96 System (Roche Diagnostics). Using the "No 5-EU" samples, the estimated amount of non-labeled mRNA in the "20 min 5-EU" and "2h

5-EU” samples was less than 9% and 2% for *ACTIN2* mRNA respectively, and less than 6% and 1% for *Clover* mRNA respectively.

#### Transient overexpression of GATA TFs in Arabidopsis protoplasts

Four plasmids were constructed using the GreenGate reaction (*46*) by combining the following plasmids: the double 35S promoter, N-terminal tag dummy sequence (addgene id: 48821), coding sequence of one of the GATA1, GATA4, and GATA6 TFs from *A. thaliana* or GFP with nuclear localization signal (addgene id: 48826), C-terminal tag dummy sequence (addgene id: 48834), rbcS terminator (addgene id: 48839), selection cassette containing the Venus protein under the seed-specific At2S3 promoter (*47*) and a destination vector. The plasmids containing the coding sequences of the three GATA TFs were synthesized by Twist Bioscience. The CDS of GATA1 was modified to remove the internal BsaI restriction site with a synonymous mutation at Gly 46 (GGT -> GGA). The four final plasmids were transformed into DH5α competent *E. coli* and purified using the QIAGEN Plasmid Plus Midi Kit (12943).

10 µg of a plasmid containing GATA TF or GFP was transformed into 200,000 Arabidopsis leaf protoplasts. Each construct was transformed in two replicates into two independent protoplast preparations, resulting in four replicates per construct. Additional two samples, per protoplast production, were transformed with 10 µg GFP control and 10 µL elution buffer, serving as a positive and negative imaging control. After 8h incubation at 25°C under constant light (85 µmol/$m^2$/s), the samples were centrifuged at 450xg for 5 minutes at room temperature and the pellets were flash-frozen in liquid nitrogen and stored at -70°C. The controls were imaged shortly before harvesting under a microscope to check the transformation efficiency - 60% and 70% protoplasts were successfully transformed in the first and second replicate, respectively.

#### RNA sequencing (RNA-Seq)

Total RNA from the overexpression experiment and polyA-selected RNA from the MPRA assay in Arabidopsis using MIX1, were used to construct RNA-Seq libraries. These libraries were prepared using the Smart-seq3 protocol (*48*), with each library constructed in multiple technical replicates and sequenced on a NovaSeqX or NovaSeq 6000 with paired-end configuration.

### Computational Methods:

#### Processing gene expression data from the *A. thaliana* accessions

Raw RNA-Seq data from ref. (*1*) were downloaded from NCBI's Sequence Read Archive (SRA) database, accession SRP074107. The data were separated into two main batches, done more than a year apart, as communicated by the authors of the original study. To verify accession identity, single nucleotide polymorphisms (SNPs) were called for each RNA-Seq sample against the reference genome (TAIR10), and compared to the SNPs of the 1,001 Genomes Project (*13*). In a few cases a mismatch between the indicated accession and the identified accession was found. Most of these cases intersected with known mix-ups in the 1001 Genomes Project (*13*, *49*). These mix-ups were corrected, or in ambiguous cases, data were not used. Reads were trimmed using Trim Galore with default parameters (*50*). Next, gene expression was quantified for all accessions which are part of the 1,001 Genomes collection, leaving out 44 accessions unique only to the gene-expression dataset. To quantify gene expression, a pseudo-genome was created for each accession by incorporating SNPs from the 1,001 Genomes vcf file, using bcftools consensus option on the reference genome (TAIR10) (*51*). Gene expression per sample was quantified using STAR against the accession's pseudo-genome using the Araport11 annotations (*52*, *53*). 72 samples with fewer than $4*10^6$ sequencing reads were omitted from further analysis. After discarding the chloroplast and mitochondria genes, gene expression was calculated by dividing the read count by gene length and normalizing to a total signal of $10^6$. Then, genes with signal less than 2 were discarded, and gene expression was $log_2$ transformed.

In a subset of the RNA-Seq libraries, a gene length bias was identified, where coverage was reduced at the 5' end of the genes. To address potential biases associated with gene length, a normalization strategy using a custom R script was implemented. Each batch was processed individually and the difference in gene expression between all possible sample pairs was calculated and correlated to the $log_2$-transformed lengths of genes. Subsequently, for each sample, the average absolute correlation across all pairs involving that sample was computed. The 10% of samples with the lowest average correlation were designated as the background group, as they were least affected by the gene length bias. These samples were left unchanged. For each non-background sample, the linear fit for the differences relative to the $log_2$-transformed gene lengths were determined when compared with each background sample. The average slope from these linear fits served as the correction factor. By subtracting the product of the correction factor and the $log_2$-transformed gene lengths to the respective samples, the gene length bias was corrected. Lastly, the total gene expression per sample was adjusted to be $10^6$ after the correction.

Raw RNA-Seq data from ref. (*2*) were downloaded from NCBI's SRA, accession SRP036643. Data of 12 samples (6 accessions x 2 conditions) were not part of the SNPs 1,001 Genomes dataset, and were omitted, as well as 46 samples with less than $10^6$ reads. Gene expression was quantified as above, first by trimming the sequence reads using Trim Galore and then quantifying using STAR against the pseudo-genomes, calculating TPMs per gene.

**eQTLs analysis**

The 1,001 Genomes vcf file was filtered for minor allele count >=5 using vcftools (--min-alleles 2 --mac 5) and converted to Plink binary format (*54*) using plink (v1.9). The Kinship matrix was calculated according to EMMA, using the *k-mers*-GWAS implementation with default parameters (*55*, *56*). For running the eQTL analysis on data from ref. (*1*), the analysis was conducted on the two batches separately. Only genes with values for at least 200 accessions were used. Gene expression ($log_2$ transformed, as described above) was normalized by subtracting the average signal of the gene in the batch, averaged over repeats if present, and transforming using a Box-Cox transformation by the MASS R library (*57*). Transformed values were used to run Genome-wide associations (GWA) with linear-mixed-models using the kinship matrix by GEMMA (v0.98.5, -lmm 2, -maf 0.05). Only SNPs up to 10 Kb downstream or upstream of the gene were used.

The eQTL analysis for data from ref. (*2*) was done for the two conditions (10°C and 16°C) separately. For each one, a gene was used if it had values for at least 120 accessions. Normalization of gene expression and following GWA was done as for the dataset from ref. (*1*). data. For each of the 4 eQTL analyses done, a threshold for significant SNPs was defined as 0.05 divided by the total number of SNPs used in the analysis, on all genes. These thresholds varied between $1.59 * 10^{-8}$ to $1.16 * 10^{-8}$, or 7.80 to 7.93 in $-log_{10}$, between the four analyses. 2760, 4259, 335, and 304 genes had at least one SNP that passed the threshold for batch1, batch2 of the data from ref. (*1*) and 10°C and 16°C conditions for data from ref. (*2*), respectively. Analysis presented in the main text (Fig. 1A-B) are from the second batch of ref. (*1*) as well as eQTL enrichment for different groups of genes (TSS-to-ATG distances or exonic fraction in 500 bp downstream of TSS), results from the other eQTL analysis is presented in the supplementary materials (Fig. S1-3).

For plotting the average enrichment of eQTLs relative to the TSS or the transcription end site (TES), each gene with significant associations had the same total contribution to the analysis, regardless of the number of associated SNPs. Two methods to weight the different associated SNPs per gene were used. First, each significant SNP got equal weight. Second, significant associated SNPs were scored according to the posterior inclusion probabilities (PIP), which take into account the linkage disequilibrium (LD) between them. To calculate the PIP, fine map analysis was conducted using the SusieR library (*58*). For the fine-map analysis a previously calculated imputed matrix of the 1,001 Genomes SNP matrix was used (*59*). For each GWA run, imputed genotypic information for the significant SNPs were extracted and PIP were calculated using the phenotype as used for the GWA.

For eQTL analysis of data from ref. (*3*) the eQTL analysis results from the original study were used. The p-values from the common effect were used with the same threshold used ($10^{-7}$). Significant SNPs per gene were weighted equally.

**Metaplots of genomic profiles around the TSS of genes**

To create a plot for genomic data around the TSS of genes, the following strategy was employed to account for overlapping and nearby genes. First, each genomic position was used at most once. For tail-to-tail genes, every upstream position was assigned to the nearest TSS. If a position was both upstream of a gene and within another gene, it was allocated to the gene it was part of, and therefore considered downstream of the TSS. If a position was located inside multiple genes, it was assigned to the TSS that was closest. The relationship between genomic positions and their corresponding TSS was stored and subsequently used to generate an average signal plot surrounding the TSS for a selected group of genes. Genomic annotations were taken from TAIR10. Analysis and plotting of genomic information was done using ‘misha’ and ‘tidyverse’ R packages (*60*). Plotting genomic data around the start codon (ATG) of genes was done using the same procedure.

Nucleotide diversity (pi) per position in the genome was calculated using vcftools with --site-pi parameter on all the SNPs table of the 1,001 Genomes project (*61*).

ChIP-Seq data from the plant chromatin state database (PCSD) was downloaded in bigwig format and imported into a misha database in R (*4*).

ChIP-Seq data for CGA1 was obtained in two independent repeats from the SRA database (*40*). Using Bowtie2, these reads were aligned to the TAIR10 reference genome with standard parameters (*62*). Subsequent peak calling was performed with MACS2 (v2.2.7.1), employing the 'callpeak' option and parameters set to '-B -q 0.01', allowing for the generation of a genomic profile (*63*). For downstream analyses, these genomic profiles were loaded into the misha database.

In the analysis of DNA Affinity Purification and Sequencing (DAP-Seq) data, narrowPeak files were used to identify the genomic positions of TF-binding peaks. These files were from the NCBI's GEO database, accession number GSE60143 (*14*). Using this data, various genomic tracks were constructed, each capturing the central points of the respective peaks. Distinct tracks were generated for each Transcription Factor (TF), done separately for each type of DNA source - genomic DNA and genomic DNA without DNA methylation. In addition, tracks were created that encompassed all peaks associated with a given TF family. This method was only applied to TF families with a minimum of 10 members. Furthermore, tracks that included all peaks linked to any TF were assembled, separate tracks were prepared for each DNA source, and one for the full dataset combining both DNA sources.

A genomic track for the GATC-motif (YVGATCBR) was constructed within a misha database. In this track, all central positions of the motif were assigned a value of 1, while all other positions were set to 0. This genomic track was used to generate the meta-plot surrounding the TSS and to define genes with a GATC motif within the initial 500 bp downstream of the TSS.

**Design of oligonucleotide pools**

Every fragment in oligonucleotide pool 1 or 2 (OP1 / OP2) was designed to have a core sequence, not longer than 160 bp surrounded by AGTTCAAACGGTCTCCACTC and AGGACGAGACCAATGTGAAC in the 5' and 3' ends, respectively.

Oligonucleotide pool 1 (OP1) was designed to incorporate fragments near the TSS of specific genes, along with control fragments. The upstream control fragments were based on the 35S, AB80, and RbcS_E9 enhancers; the fragments were taken from ref. (*10*). The downstream control fragment was designed according to the intron of the UBQ10 gene, from 3 base pairs (bp) past the donor site to 3 bp before the acceptor site. Fifty oligonucleotides of 160 bp each were designed to cover each control fragment, with an overlap of 150 bp between every two consecutive oligos (for instance, 1-160, 11-170, 21-180, and so on).

The oligos drawn from near the TSS of genes were designed in the following way: Genes of *Arabidopsis thaliana* were selected according to the TAIR10 annotation (*64*) under certain criteria: (1) excluding genes on the mitochondria or chloroplast genomes, (2) ensuring the closest upstream gene is at least 50 bp away from the gene's TSS, (3) only considering genes with a single TSS, (4) focusing on genes of at least 500 bp in length, (5) only including protein-coding genes, and (6) selecting highly expressed genes with expression in Col-0 of at least 10 in $\log_2$(TPM) in ref. (*1*).

Three fragments were obtained from the genome for each of the 7,775 genes that passed this filtering process: 200 bp to 41 bp upstream of the TSS, 41 bp to 200 bp and 201 bp to 360 bp downstream the TSS. If any of these fragments contained a donor or an acceptor splicing site, the sequences surrounding it, 5 bp (-1 to +3) for donor sites and 3 bp (-1 to +1) for acceptor sites, were removed. Any genes with a BsmbI or BsaI recognition site within the three TSS-proximate sequences were then removed, reducing the gene count to 4,884. Finally, 3,991 genes with the highest expression were chosen, and the three fragments around each of these genes' TSS were included in the oligonucleotide pool.

Oligonucleotide pool 2 (OP2) was constructed to contain altered versions of fragments from OP1, which were split into three distinct sets. The goal of Set 1 was to mutate the GATC motifs. 823 fragments originating downstream of the TSS in OP1 were chosen, each with at least one GATC motif, adding up to a total of 917 GATC elements. New fragments were created for each of these fragments by either of the following: (1) removing the GATC motif, resulting in shorter fragments, (2) rearranging / shuffling the 8 bp of the GATC element to ensure disruption of the original GATC sequence, and (3) changing the 6th nucleotide in the element, from "C", to an "A".

If a single fragment contained more than one GATC motif, each of these modifications was applied to every possible subset of elements. For instance, if a fragment had three GATC motifs, it would yield seven subsets ($2^3$-1), and each of the three transformations would be applied to each of these, resulting in 21 mutated fragments. In total, 3,216 fragments were created for Set 1.

Set 2 was designed with the objective of introducing GATC motifs into the fragments. For this purpose, 221 fragments which can be detected consistently in our MPRA libraries were arbitrarily selected from OP1, with 75% originating downstream and 25% originating upstream of the TSS. For each chosen fragment, an incremental series of 8 fragments was created, with each successive fragment incorporating an additional CAGATCTG sequence. There was a minimum of 2 bp between two adjacent GATC motifs. As a result, a total of 1,768 fragments (221 * 8) were designed for Set 2.

Set 3 was designed for a deep mutational screening of a small subset of fragments. To this end, 20 fragments from OP1, all originating from downstream of the TSS, were selected. The chosen fragments were among the top 100 enhancing fragments from the six libraries of tomato, *A. thaliana*, and *N. benthamiana*, in each of the two backbones when positioned downstream the TSS. Among these 20 fragments, seven were devoid of any GATC elements, from which two fragments lacked any GATC sequences (only the 4 bp). Out of the 13 fragments that did have a GATC element, two of them had two such elements. In the scope of this paper, only these 13 are analyzed, while all data can be found in the supplementary tables.

Two series of modifications were created for each of these fragments. First, every alternating 10 bp sequence, that is positions 1-10, 3-12, 5-14,..., and 151-160, was removed, leading to 76 fragments each of 150 base pairs in length. Second, each single bp in the fragment was substituted with one of the other three potential nucleotides, or it was completely removed to yield a 159 base pair fragment. This second set of alterations resulted in 640 fragments (160 * 4) from each original fragment.

Duplicate fragments in the OP2 design were subsequently identified and removed. These duplicates could have been the result of the removal of a single bp within a region composed of the same nucleotides in Set 3, or they could have been due to overlaps between Set 1 and Set 3.

**Connecting barcode to enhancer fragment and plasmid backbone**

Paired-end sequence reads were used to connect barcodes to inserted fragments. In all 8 constructed libraries, the barcode was in R2 read and the inserted fragment (except in the two *_rmBsaI backbones) in R1. The sequence of R2, for example, in pPSup_iGFP_v2 based library begins with:

{BBD}AGCTCCTCGCCCTTGCTCACNNBNNBNNBNNBNNBCATGGT, where {BBD} can be either BBD, BD, D, or nothing. The underlined sequences are constant and the NNBx5 is the barcode. In all libraries, R2 starts with the same form, with different constant sequences and different degenerative nucleotides. R1 starts, for example, with the following form in pPSup_iGFP_v2-based library: {BBH}CTTGATATCGAATTCCACTCNNNNNNN…. also here the {BBH} represent up to 3 degenerative bp and the constant sequences is underlined. The NNN… represent the sequence of the inserted fragment. R1 sequences from all libraries have a similar form, with the exception of the two *_rmBsaI-based libraries, where there is no cloned sequence and thus the constant sequence is longer. For pPSint_v2 and pPSint_v2_pTRP1 cloned with OP1+OP2, R1 sequences were of length 210 bp and the read ended with more constant sequences after the up to 160 bp of the synthesized-inserted fragments.

Paired-end reads were filtered to ones that have a maximum of 1 bp mutation in each of the three constant sequences: preceding the inserted fragments in R1, prior to the barcode in R2, or within the 10 bp following the barcode in R2. This means that a read can have 1 bp mutation in all three sequences and still be used. Following the filtering steps the barcode and beginning of the enhancer fragment are extracted. Barcodes that do not fit the VNNx5 pattern are filtered out.

The subsequent processing was carried out independently for OP1+OP2-based libraries and all other libraries. For the latter group containing enhancer fragments, the extracted enhancer fragments were aligned to the 12,000 synthesized oligos using Bowtie (with parameters -v 1 -a --best --strata) (*65*). Barcodes associated with more than one enhancer fragment were flagged, and were not used in following analysis. Due to varying sequence coverage and library complexities (particularly between the *_rmBsaI libraries and others), the libraries were downsampled, targeting for 135 appearances of the 50th most frequent barcode.

In the case of OP1+OP2-based libraries, the full 160 bp of the inserted fragment had to be utilized, as OP2 contained sets of all 1 bp mutations derived from specific fragments. Subsequently, paired-reads that passed the initial filtering (i.e., <= 1 bp mutation in constant regions) underwent further filtering to detect the presence of the expected constant sequence ("GAGTAATTGC") after the enhancer, eliminating less than 5% of reads. All enhancer fragments flanked by constant sequences connected to the same barcode were processed together, to get a mapping of each barcode. To link barcodes to a unique enhancer fragment, the following criteria were applied: (i) The barcode appeared at least in two reads, (ii) The most abundant connected sequence matched one of the synthesized fragments (OP1+OP2) exactly, (iii) None of the other connected sequences with more than 2 bp mismatches to the most abundant sequence were found in the list of synthesized-fragment enhancers (OP1+OP2), (iv) The total number of sequences with up to 2 bp mutations from the most abundant sequence, including the most abundant sequence itself, constituted at least 80% of all sequences connected to the barcode. If a barcode appeared at least twice but couldn't be linked to a single enhancer it was flagged, and was not used in the following analysis.

For the MPRA experiment, groups of libraries were mixed and assayed together. Barcodes from these libraries have to be linked back both to the enhancer and the original library, and if the same barcode appeared in multiple such libraries it cannot be used. To this end, the barcodes to enhancer-fragment dictionaries from each mix of libraries were combined, and barcodes appearing in multiple libraries were marked. Finally, for each of the two mixes of libraries employed in the MPRA experiment, a Tn5-derived DNA sequencing library was generated and sequenced. Barcodes were extracted from the reads based on the following patterns: ACCATG(VNNx5)GTGAGC or GTGATG(VNNx5)GTGAGC. The extracted barcodes were subsequently traced back to their original libraries by searching the corresponding barcode-to-enhancer-fragment dictionaries for each mix, allowing for the inference of actual library-mix ratios.

This procedure produced (1) a dictionary between enhancer X position to barcodes, (2) the normalization weights of how many times each barcode and enhancer appeared in the transformed mix.

**MPRA quantification of gene expression**

Paired-end reads from the MPRA RNA-Seq libraries were used. The R1 structure from both p35S- and pTRP1-based libraries shared the same structure: {DDHHBDBDHDV}GAACTTGTGGCCGTTTACG. In this sequence, the underlined sections represent constant regions, while the degenerate sequence is a unique molecular identifier (UMI), of length 8 bp up to 11 bp, incorporated during the RT step (*66*). In the case of R2, the p35S-based library started with: {HBB}TTCTAGTATACTAAACCATGVNNVNNVNNVNNVNNGTGAGCAAGG whereas the pTRP1-based library started with:

{HHV}TGAGCAATCGAGTGATGVNNVNNVNNVNNVNNGTGAGCAAGG. As described in the previous section, the underlined sequences are constant regions, {HBB} and {HHV} indicate a possible shift of 0-3 bp comprising a random sequence, and VNNx5 represents the barcode sequence. Subsequently, the read-pairs were filtered to include only those exhibiting a maximum of 2-bp mismatch in each of the constant sequences, and having the barcode and UMI in the right lengths. Following this, barcodes and UMIs were extracted from the filtered reads.

The Barcode-UMI pairs were processed in several stages. Firstly, these pairs were collapsed to retain only unique combinations, and the frequency of each unique pair was recorded.

Both the barcode and the UMI contained variable sequences, where specific positions could take on one of three or all four possible nucleotides. This variability allows the estimation of the total sequencing errors in the barcodes or UMIs. With the UMIs used for quantifying expression levels, sequencing errors could potentially lead to inaccurate gene expression estimates.

Given that the MPRA RNA-Seq libraries were sequenced to high effective coverage (sometimes exceeding 20X), it was plausible that a UMI could occasionally be read with a sequencing error. Consequently, the percentage of pairs with a UMI that did not adhere to the variable sequence constraints and a barcode that did was calculated. This calculation was performed at different thresholds of the minimal frequency of barcode-UMI pairs.

The expectation was that setting a higher threshold for the frequency of barcode-UMI pair appearance would also reduce the number of UMIs with sequencing errors. Owing to the degenerate sequence's structure, only one out of three possible errors could be detected for a single base mutation. Therefore, the actual number of UMI sequence errors was assumed to be three times the percentage of detected errors for each frequency threshold.

However, since UMIs with sequences that did not fit the variable sequence could be removed, the real error in estimation was up to double the percentage of detected error sequences. A threshold of less than 0.75% error UMI sequences was set, leading to the potential of up to 1.5% of UMIs being incorrect. It was, however, expected to be lower due to the RT primers initially containing a UMI that didn't fit the pattern due to errors in the synthesis.

The outcome threshold for appearances from this procedure ranged from 2 to 4 in the various experiments. This approach enabled the minimization of errors while still retaining a significant portion of the data for analysis.

Subsequently, only the barcodes included in the associated mix dictionary were retained. The RNA count for each enhancer X position was determined by the total number of unique barcode-UMI pairs linked to it. To obtain the gene expression level for each construct, this RNA count was normalized by the corresponding normalized weight for the enhancer X position in the mix. Lastly, the total signal for each experiment was normalized to a value of one million, providing a measure parallel to the Transcripts Per Million (TPM) score.

In the majority of the analyses presented in this study, the mean expression of the enhancer, derived from the three or four experimental replicates, is used. For Fig. 2C, the expression of each control enhancer or iUBQ10 was calculated by taking the average gene expression from all its constituent fragments.

**Determining overrepresented *k*-mers in downstream enhancers**

To uncover overrepresented 6-mers (*k*-mers composed of 6 base pairs) in active downstream enhancers, each species and backbone combination was evaluated separately. Only fragments derived downstream of the TSS were taken into account, and expression levels from downstream positioning were used. For each unique 6-mer (considering reverse complement as well), the enhancers were classified into two groups: those that contained the 6-mer or its complement, and those that did not. A 6-mer was considered in the analysis only if it was present in more than 20 fragments across both groups. The expression levels of the constructs in the MPRA setup were compared between the groups using the Mann-Whitney U test, yielding a p-value. Moreover, the average $\log_2$ expression for each group was determined, and the difference between the groups was assessed. A significance threshold was established at $-\log_{10}(0.05/\#tests)$, where #tests signifies the total number of tests executed for all 6-mers across the eight combinations. This threshold is demonstrated in Figs. 3A, S12, and S13.

**Associations of GATC variation to gene expression in the 1,001G**

To detect variations in GATC motifs across the 1,001 Genomes data set, genomic coordinates having a GATC 4-bp sequence within the 1,001 Genomes pseudo-genomes were assembled. The list was filtered down to coordinates within 1 kb of the TSS of genes used in the eQTLs analysis of ref. (*1*) dataset. For each relevant position, an 8-bp motif-allele was extracted, which includes an extra 2 bp flanking the GATC 4-bp sequence from all pseudo-genomes. Alleles at each site were designated as GATC+ if they aligned with the YVGATCBR motif, and as GATC- if devoid of the GATC sequence. Alleles with a GATC not matching the YVGATCBR motif were excluded. Subsequently, for each identified position, expression values from the two batches from ref. (*1*) were compared between the GATC+ and GATC- accessions. A statistical analysis was conducted using the Mann-Whitney U test in R for positions with at least 10 elements in both comparison groups. Positions were considered significant if they had a p-value less than 0.05 divided by the total number of tests and showed an average $\log_2$(expression) difference of at least 0.5. Such significant positions were then grouped based on their proximity to the TSS and whether the GATC motif amplified the expression. In total, 111 associations meeting the established criteria were identified. Of the 111 associations identified, distributions for GATC- and GATC+ were: 13 each for -1000 to -500 from the TSS; 17 GATC- and 11 GATC+ for -500 to TSS; 7 GATC- and 18 GATC+ for TSS to +500; and 20 GATC- against 12 GATC+ for +500 to +1000.

**Evaluation of effect sizes of the GATC motif or other 6-mers**

The influence of the GATC motif or 6-mers (as depicted in Fig. 4B) on gene expression and other genomic datasets was determined by analyzing the slope of a linear relationship (defined as the effect size) and the accompanying p-value from the fit. A linear regression, using R's 'lm' function, was performed to relate the frequency of motif occurrences to the genomic database.

**Analysis of gene expression and other genomic measurements from published *A. thaliana* studies**

Meta analysis of different tissue expression in Arabidopsis was done with data from ref. (*22*), processed as described above, and on a compendium of tissue-specific gene expression data from AtGenExpress, produced with the Affymetrix ATH1 microarray platform, from the Bio-Analytic Resource for Plant Biology (BAR) (*21*, *67*). Due to concerns raised regarding contamination of embryo samples in gene expression datasets (*68*), the embryo related samples from the latter dataset were replaced with the full compendium of gene expression data collected in a newer dataset (*69*). These data also included data from refs. (*70*, *71*) which were used in Fig. 4E. In all cases, values of replicates from the same experiment were averaged.

Root expression data at the single-cell level was downloaded from NCBI's GEO database using the accession number GSE152766 (*27*). Additionally, single-cell data for the vegetative shoot apex was obtained directly via personal correspondence (*28*). Both datasets, retaining their initial cell annotations, were processed using Seurat (v4.3.0) (*72*).

mRNA synthesis rate and mRNA half-lives per gene were used as reported (*26*).

Data on H3K4me3 and H3K36me3 were from ref. (*4*). Using the misha R package, the average signal per gene was calculated and averaged over repeats. RNA polymerase binding information was taken from ref. (*73*), specifically from the control dataset (NS), which was downloaded from the NCBI's GEO database under the accession number GSE122804 (*73*).

**Evolutionary analysis of land plant genomes and transcriptomes**

For the evolutionary examination of the GATC motif's impact on genome-wide gene expression, genomes and annotations from the following sources were employed: Arabidopsis - TAIR10 (*64*); *S. lycopersicum* - ITAG4.0 (*74*); *O. sativa* - v7 (*75*); *Z. mays* - B73 NAM-5.0.55 (*76*); *P. tabuliformis* - v1.0 (*77*); *C. richardii* - v2.1 (*78*); *S. moellendorffii* - v1.0 (*79*) annotations from NCBI Annotation Release 100; *M. polymorpha* - v3.1 (*80*); *P. patens* - v3.3 (*81*).

Gene expression data were sourced and processed as described below. For *A. thaliana*, raw data were obtained from the SRA database under the accession PRJNA314076 (*22*). These 138 samples underwent processing through the Nextflow (v22.10.7.5854) tool using the nf-core RNA-Seq pipeline (v3.6) set to default parameters (*82*, *83*). In the case of tomato, maize, rice, and *P. patens*, processed data were directly downloaded from respective studies (*84–87*). For *C. richardii*, the processed dataset was from ref. (*88*) under the NCBI's GEO database accession GSE212819.

For *P. tabuliformis*, raw RNA-Seq data were accessed from SRA under accession PRJNA173457 (*77*). Due to the large size of chromosomes (>$2*10^9$ bp), chromosomes were segmented into pseudo-chromosomes no longer than $10^8$ bp. Any intersecting features between two consecutive pseudo-chromosomes were discarded. Using GffRead (v0.11.8) (*89*), a fasta file comprising transcripts was generated from the modified chromosome and annotations. Subsequent quantification of gene expression across the 136 samples employed Salmon (v1.10) (*90*).

For *S. moellendorffii* and *M. polymorpha*, RNA-Seq datasets were accessed from ref. (*91*) (SRR1740446-SRR1740451) and ref. (*92*) (DRR130762-DRR130768) in the SRA, respectively. These samples were processed using the nf-core RNA-Seq pipeline similarly to the Arabidopsis samples.

For every gene across each species, a 500 bp sequence downstream of the TSS was extracted, followed by counting the occurrences of the YVGATCBR motif. The effect size and the p-value were calculated using a linear fit as explained above.

**Comparing orthologs between Arabidopsis and tomato**

Protein sequences from Arabidopsis (TAIR10) (*64*) and *Solanum lycopersicum* (ITAG4.0) (*74*) were utilized to conduct orthology analysis using OrthoFinder (version 2.5.4) (*19*). Prior to analysis, one variant per gene was selected using the primary_transcript.py tool provided by OrthoFinder. The identification of one-to-one orthologous genes was carried out using the Phylogenetic Hierarchical Orthogroups generated by OrthoFinder, and these orthologs were subsequently employed for further analysis.

**Processing of RNA-Seq**

RNA-Seq libraries were prepared using the Smart-seq3 protocol (*48*) , which generates two types of sequence fragments. The first type consists of reads derived from the 5' end of mRNAs, starting with an 11 bp constant sequence (ATTGCGCAATG) followed by a UMI and "GGG". These reads, which represent more than 80% of our library, were used for analysis. The second type, originating from the middle of the mRNAs, lacks a UMI and is thus more susceptible to PCR amplification bias. For UMI processing, UMIs were extracted, and reads (following the “GGG” in R1) that shared identical UMIs and had up to 2 bp mismatches in read R1 were collapsed using the clumpify.sh script from the BBMap suite (*93*). The collapsed reads were then trimmed using Trim Galore with default settings (*50*). Gene expression quantification was performed using Salmon (v1.5.2) with the parameters

--validateMappings -l A (*90*)against transcripts from TAIR10 with manual addition of the GFP gene.

For the MPRA libraries of Arabidopsis with MIX1, the same collapsed and trimmed reads were used to assess splicing efficiency.

**Assessing splicing efficiency and pTRP1- vs p35S-based total RNA levels**

The 5' reads of the SMART-Seq3 library from the full RNA-Seq of the Arabidopsis MPRA experiment with MIX1 were used to assess splicing efficiency. After removing the constant sequence and the UMIs, the reads were filtered to those containing a portion of the sequence between the barcode and the start of the intron (see Fig. S7). This was done by taking all 15 bp *k*-mers from this sequence and filtering out all 5' clean reads from the RNA-Seq that had at least one of these *k*-mers, resulting in a total of 20,972 sequences across all four repeats, ranging from 2,735 to 7,092 across the four repeats. When these sequences were further filtered to include sequences from the region upstream of the barcode that differed between the pTRP1 and 35S-based libraries, 90.7% of the reads were uniquely assigned to one of the two libraries. Using the ratio between the number of reads assigned to each library, we estimated the ratio between the backbone without insert of the 35S-based library to be higher than the pTRP1-based by log2 of 3.13, 1.77, 2.3, 2.25 respectively in the four repeats.

The 20,972 5'-end cleaned sequence reads were used to assess splicing efficiency within the MPRA setup. Among these reads, 7,789 R2 sequences contained the sequence "CTTCGCCCTC", which is located just upstream of the splice site and at least 6 bp downstream of this sequence. Of these sequences, 7,405 represented exon-exon junctions and 100 represented exon-intron junctions. Therefore, the estimated splicing efficiency is 98.7% for the four libraries, or 99.39%, 98.91%, 98.55%, and 98.34% for the four repeats, respectively.

**References:**

1. T. Kawakatsu, S.-S. C. Huang, F. Jupe, E. Sasaki, R. J. Schmitz, M. A. Urich, R. Castanon, J. R. Nery, C. Barragan, Y. He, H. Chen, M. Dubin, C.-R. Lee, C. Wang, F. Bemm, C. Becker, R. O'Neil, R. C. O'Malley, D. X. Quarless, 1001 Genomes Consortium, N. J. Schork, D. Weigel, M. Nordborg, J. R. Ecker, Epigenomic Diversity in a Global Collection of Arabidopsis thaliana Accessions. *Cell* **166**, 492–505 (2016).

2. M. J. Dubin, P. Zhang, D. Meng, M.-S. Remigereau, E. J. Osborne, F. Paolo Casale, P. Drewe, A. Kahles, G. Jean, B. Vilhjálmsson, J. Jagoda, S. Irez, V. Voronin, Q. Song, Q. Long, G. Rätsch, O. Stegle, R. M. Clark, M. Nordborg, DNA methylation in Arabidopsis has a genetic basis and shows evidence of local adaptation. *Elife* **4**, e05255 (2015).

3. P. Clauw, F. Coppens, A. Korte, D. Herman, B. Slabbinck, S. Dhondt, T. Van Daele, L. De Milde, M. Vermeersch, K. Maleux, S. Maere, N. Gonzalez, D. Inzé, Leaf Growth Response to Mild Drought: Natural Variation in Arabidopsis Sheds Light on Trait Architecture. *Plant Cell* **28**, 2417–2434 (2016).

4. Y. Liu, T. Tian, K. Zhang, Q. You, H. Yan, N. Zhao, X. Yi, W. Xu, Z. Su, PCSD: a plant chromatin state database. *Nucleic Acids Res.* **46**, D1157–D1167 (2018).

5. H. Stroud, S. Otero, B. Desvoyes, E. Ramírez-Parra, S. E. Jacobsen, C. Gutierrez, Genome-wide analysis of histone H3.1 and H3.3 variants in Arabidopsis thaliana. *Proc. Natl. Acad. Sci. U. S. A.* **109**, 5370–5375 (2012).

6. H. Wollmann, S. Holec, K. Alden, N. D. Clarke, P.-É. Jacques, F. Berger, Dynamic deposition of histone variant H3.3 accompanies developmental remodeling of the Arabidopsis transcriptome. *PLoS Genet.* **8**, e1002658 (2012).

7. K. N. Chang, S. Zhong, M. T. Weirauch, G. Hon, M. Pelizzola, H. Li, S.-S. C. Huang, R. J. Schmitz, M. A. Urich, D. Kuo, J. R. Nery, H. Qiao, A. Yang, A. Jamali, H. Chen, T. Ideker, B. Ren, Z. Bar-Joseph, T. R. Hughes, J. R. Ecker, Temporal transcriptional response to ethylene gas drives growth hormone cross-regulation in Arabidopsis. *Elife* **2**, e00675 (2013).

8. L. Vercruyssen, A. Verkest, N. Gonzalez, K. S. Heyndrickx, D. Eeckhout, S.-K. Han, T. Jégu, R. Archacki, J. Van Leene, M. Andriankaja, S. De Bodt, T. Abeel, F. Coppens, S. Dhondt, L. De Milde, M. Vermeersch, K. Maleux, K. Gevaert, A. Jerzmanowski, M. Benhamed, D. Wagner, K. Vandepoele, G. De Jaeger, D. Inzé, ANGUSTIFOLIA3 binds to SWI/SNF chromatin remodeling complexes to regulate transcription during Arabidopsis leaf development. *Plant Cell* **26**, 210–229 (2014).

9. L. Yant, J. Mathieu, T. T. Dinh, F. Ott, C. Lanz, H. Wollmann, X. Chen, M. Schmid, Orchestration of the floral transition and floral development in Arabidopsis by the bifunctional transcription factor APETALA2. *Plant Cell* **22**, 2156–2170 (2010).

10. T. Jores, J. Tonnies, M. W. Dorrity, J. T. Cuperus, S. Fields, C. Queitsch, Identification of Plant Enhancers and Their Constituent Elements by STARR-seq in Tobacco Leaves. *Plant Cell* **32**, 2120–2131 (2020).

11. S. Kanoria, P. K. Burma, A 28 nt long synthetic 5'UTR (synJ) as an enhancer of transgene expression in dicotyledonous plants. *BMC Biotechnol.* **12**, 85 (2012).

12. E. X. Szabo, P. Reichert, M.-K. Lehniger, M. Ohmer, M. de Francisco Amorim, U. Gowik, C. Schmitz-Linneweber, S. Laubinger, Metabolic Labeling of RNAs Uncovers Hidden Features and Dynamics of the Arabidopsis Transcriptome. *Plant Cell* **32**, 871–887 (2020).

13. 1001 Genomes Consortium, 1,135 Genomes Reveal the Global Pattern of Polymorphism in Arabidopsis thaliana. *Cell* **166**, 481–491 (2016).

14. R. C. O'Malley, S.-S. C. Huang, L. Song, M. G. Lewsey, A. Bartlett, J. R. Nery, M. Galli, A. Gallavotti, J. R. Ecker, Cistrome and Epicistrome Features Shape the Regulatory DNA Landscape. *Cell* **165**, 1280–1292 (2016).

15. J. C. Reyes, M. I. Muro-Pastor, F. J. Florencio, The GATA family of transcription factors in Arabidopsis and rice. *Plant Physiol.* **134**, 1718–1732 (2004).

16. C. Schwechheimer, P. M. Schröder, C. E. Blaby-Haas, Plant GATA Factors: Their Biology, Phylogeny, and Phylogenomics. *Annu. Rev. Plant Biol.* **73**, 123–148 (2022).

17. X. Tu, M. K. Mejía-Guerra, J. A. Valdes Franco, D. Tzeng, P.-Y. Chu, W. Shen, Y. Wei, X. Dai, P. Li, E. S. Buckler, S. Zhong, Reconstructing the maize leaf regulatory network using ChIP-seq data of 104 transcription factors. *Nat. Commun.* **11**, 5089 (2020).

18. M. I. Love, W. Huber, S. Anders, Moderated estimation of fold change and dispersion for RNA-seq data with DESeq2. *Genome Biol.* **15**, 550 (2014).

19. D. M. Emms, S. Kelly, OrthoFinder: phylogenetic orthology inference for comparative genomics. *Genome Biol.* **20**, 238 (2019).

20. Z. Lin, W.-H. Li, Evolution of 5' untranslated region length and gene expression reprogramming in yeasts. *Mol. Biol. Evol.* **29**, 81–89 (2012).

21. M. Schmid, T. S. Davison, S. R. Henz, U. J. Pape, M. Demar, M. Vingron, B. Schölkopf, D. Weigel, J. U. Lohmann, A gene expression map of Arabidopsis thaliana development. *Nat. Genet.* **37**, 501–506 (2005).

22. A. V. Klepikova, A. S. Kasianov, E. S. Gerasimov, M. D. Logacheva, A. A. Penin, A high resolution map of the Arabidopsis thaliana developmental transcriptome based on RNA-seq profiling. *Plant J.* **88**, 1058–1070 (2016).

23. A. J. Bewick, L. Ji, C. E. Niederhuth, E.-M. Willing, B. T. Hofmeister, X. Shi, L. Wang, Z. Lu, N. A. Rohr, B. Hartwig, C. Kiefer, R. B. Deal, J. Schmutz, J. Grimwood, H. Stroud, S. E. Jacobsen, K. Schneeberger, X. Zhang, R. J. Schmitz, On the origin and evolutionary consequences of gene body DNA methylation. *Proc. Natl. Acad. Sci. U. S. A.* **113**, 9111–9116 (2016).

24. S. Inagaki, M. Takahashi, A. Hosaka, T. Ito, A. Toyoda, A. Fujiyama, Y. Tarutani, T. Kakutani, Gene-body chromatin modification dynamics mediate epigenome

differentiation in Arabidopsis. *EMBO J.* **36**, 970–980 (2017).

25. M. V. C. Greenberg, A. Deleris, C. J. Hale, A. Liu, S. Feng, S. E. Jacobsen, Interplay between active chromatin marks and RNA-directed DNA methylation in Arabidopsis thaliana. *PLoS Genet.* **9**, e1003946 (2013).

26. K. Sidaway-Lee, M. J. Costa, D. A. Rand, B. Finkenstadt, S. Penfield, Direct measurement of transcription rates reveals multiple mechanisms for configuration of the Arabidopsis ambient temperature response. *Genome Biol.* **15**, R45 (2014).

27. R. Shahan, C.-W. Hsu, T. M. Nolan, B. J. Cole, I. W. Taylor, L. Greenstreet, S. Zhang, A. Afanassiev, A. H. C. Vlot, G. Schiebinger, P. N. Benfey, U. Ohler, A single-cell Arabidopsis root atlas reveals developmental trajectories in wild-type and cell identity mutants. *Dev. Cell* **57**, 543–560.e9 (2022).

28. T.-Q. Zhang, Y. Chen, J.-W. Wang, A single-cell analysis of the Arabidopsis vegetative shoot apex. *Dev. Cell* **56**, 1056–1074.e8 (2021).

29. S. H. Orkin, Globin gene regulation and switching: circa 1990. *Cell* **63**, 665–672 (1990).

30. M. Merika, S. H. Orkin, DNA-binding specificity of GATA family transcription factors. *Mol. Cell. Biol.* **13**, 3999–4010 (1993).

31. P. V. Pedone, J. G. Omichinski, P. Nony, C. Trainor, A. M. Gronenborn, G. M. Clore, G. Felsenfeld, The N-terminal fingers of chicken GATA-2 and GATA–3 are independent sequence-specific DNA binding domains. *EMBO J.* **16**, 2874–2882 (1997).

32. A. Newton, J. Mackay, M. Crossley, The N-terminal zinc finger of the erythroid transcription factor GATA-1 binds GATC motifs in DNA. *J. Biol. Chem.* **276**, 35794–35801 (2001).

33. L. J. Ko, J. D. Engel, DNA-binding specificities of the GATA transcription factor family. *Mol. Cell. Biol.* **13**, 4011–4022 (1993).

34. Y.-H. Shim, J. J. Bonner, T. Blumenthal, Activity of aC. elegansGATA Transcription Factor, ELT-1, Expressed in Yeast. *J. Mol. Biol.* **253**, 665–676 (1995).

35. C. Liu, L. Kang, M. Lin, J. Bi, Z. Liu, S. Yuan, Molecular Mechanism by Which the GATA Transcription Factor CcNsdD2 Regulates the Developmental Fate of Coprinopsis cinerea under Dark or Light Conditions. *MBio* **13**, e0362621 (2021).

36. J. A. Castro-Mondragon, R. Riudavets-Puig, I. Rauluseviciute, R. B. Lemma, L. Turchi, R. Blanc-Mathieu, J. Lucas, P. Boddie, A. Khan, N. Manosalva Pérez, O. Fornes, T. Y. Leung, A. Aguirre, F. Hammal, D. Schmelter, D. Baranasic, B. Ballester, A. Sandelin, B. Lenhard, K. Vandepoele, W. W. Wasserman, F. Parcy, A. Mathelier, JASPAR 2022: the 9th release of the open-access database of

transcription factor binding profiles. *Nucleic Acids Res.* **50**, D165–D173 (2022).

37. J. A. Lowry, W. R. Atchley, Molecular evolution of the GATA family of transcription factors: conservation within the DNA-binding domain. *J. Mol. Evol.* **50**, 103–115 (2000).

38. J. M. Franco-Zorrilla, I. López-Vidriero, J. L. Carrasco, M. Godoy, P. Vera, R. Solano, DNA-binding specificities of plant transcription factors and their potential to define target genes. *Proc. Natl. Acad. Sci. U. S. A.* **111**, 2367–2372 (2014).

39. M. T. Weirauch, A. Yang, M. Albu, A. G. Cote, A. Montenegro-Montero, P. Drewe, H. S. Najafabadi, S. A. Lambert, I. Mann, K. Cook, H. Zheng, A. Goity, H. van Bakel, J.-C. Lozano, M. Galli, M. G. Lewsey, E. Huang, T. Mukherjee, X. Chen, J. S. Reece-Hoyes, S. Govindarajan, G. Shaulsky, A. J. M. Walhout, F.-Y. Bouget, G. Ratsch, L. F. Larrondo, J. R. Ecker, T. R. Hughes, Determination and inference of eukaryotic transcription factor sequence specificity. *Cell* **158**, 1431–1443 (2014).

40. Z. Xu, J. A. Casaretto, Y.-M. Bi, S. J. Rothstein, Genome-wide binding analysis of AtGNC and AtCGA1 demonstrates their cross-regulation and common and specific functions. *Plant Direct* **1**, e00016 (2017).

41. K. Sugimoto, S. Takeda, H. Hirochika, Transcriptional activation mediated by binding of a plant GATA-type zinc finger protein AGP1 to the AG-motif (AGATCCAA) of the wound-inducible Myb gene NtMyb2. *Plant J.* **36**, 550–564 (2003).

42. Y. Liu, B. Patra, S. Pattanaik, Y. Wang, L. Yuan, GATA and Phytochrome Interacting Factor Transcription Factors Regulate Light-Induced Vindoline Biosynthesis in Catharanthus roseus. *Plant Physiol.* **180**, 1336–1350 (2019).

43. C. Neumayr, M. Pagani, A. Stark, C. D. Arnold, STARR-seq and UMI-STARR-seq: Assessing Enhancer Activities for Genome-Wide-, High-, and Low-Complexity Candidate Libraries. *Curr. Protoc. Mol. Biol.* **128**, e105 (2019).

44. D. Ben-Tov, F. Mafessoni, A. Cucuy, A. Honig, C. Melamed-Bessudo, A. A. Levy, Uncovering the Dynamics of Precise Repair at CRISPR/Cas9-induced Double-Strand Breaks, *bioRxiv* (2023)p. 2023.01.10.523377.

45. J. Cao, D. Yao, F. Lin, M. Jiang, PEG-mediated transient gene expression and silencing system in maize mesophyll protoplasts: a valuable tool for signal transduction study in maize. *Acta Physiol. Plant* **36**, 1271–1281 (2014).

46. A. Lampropoulos, Z. Sutikovic, C. Wenzl, I. Maegele, J. U. Lohmann, J. Forner, GreenGate---a novel, versatile, and efficient cloning system for plant transgenesis. *PLoS One* **8**, e83043 (2013).

47. S. Bensmihen, A. To, G. Lambert, T. Kroj, J. Giraudat, F. Parcy, Analysis of an activated ABI5 allele using a new selection method for transgenic Arabidopsis seeds. *FEBS Lett.* **561**, 127–131 (2004).

48. M. Hagemann-Jensen, C. Ziegenhain, P. Chen, D. Ramsköld, G.-J. Hendriks, A. J. M. Larsson, O. R. Faridani, R. Sandberg, Single-cell RNA counting at allele and isoform resolution using Smart-seq3. *Nat. Biotechnol.* **38**, 708–714 (2020).

49. R. Pisupati, I. Reichardt, Ü. Seren, P. Korte, V. Nizhynska, E. Kerdaffrec, K. Uzunova, F. A. Rabanal, D. L. Filiault, M. Nordborg, Verification of Arabidopsis stock collections using SNPmatch, a tool for genotyping high-plexed samples. *Sci Data* **4**, 170184 (2017).

50. F. Krueger, F. James, P. Ewels, E. Afyounian, M. Weinstein, B. Schuster-Boeckler, G. Hulselmans, sclamons, *FelixKrueger/TrimGalore: v0.6.10 - Add Default Decompression Path* (2023; https://zenodo.org/record/7598955).

51. P. Danecek, J. K. Bonfield, J. Liddle, J. Marshall, V. Ohan, M. O. Pollard, A. Whitwham, T. Keane, S. A. McCarthy, R. M. Davies, H. Li, Twelve years of SAMtools and BCFtools. *Gigascience* **10** (2021).

52. C.-Y. Cheng, V. Krishnakumar, A. P. Chan, F. Thibaud-Nissen, S. Schobel, C. D. Town, Araport11: a complete reannotation of the Arabidopsis thaliana reference genome. *Plant J.* **89**, 789–804 (2017).

53. A. Dobin, C. A. Davis, F. Schlesinger, J. Drenkow, C. Zaleski, S. Jha, P. Batut, M. Chaisson, T. R. Gingeras, STAR: ultrafast universal RNA-seq aligner. *Bioinformatics* **29**, 15–21 (2013).

54. S. Purcell, PLINK : a toolset for whole-genome association and population-based linkage analysis. *Am. J. Hum. Genet.* **81**, 559–575 (2007).

55. H. M. Kang, N. A. Zaitlen, C. M. Wade, A. Kirby, D. Heckerman, M. J. Daly, E. Eskin, Efficient control of population structure in model organism association mapping. *Genetics* **178**, 1709–1723 (2008).

56. Y. Voichek, D. Weigel, Identifying genetic variants underlying phenotypic variation in plants without complete genomes. *Nat. Genet.* **52**, 534–540 (2020).

57. W. N. Venables, B. D. Ripley, Modern Applied Statistics with S, Springer, New York: ISBN 0-387-95457-0. [Preprint] (2002).

58. Y. Zou, P. Carbonetto, G. Wang, M. Stephens, Fine-mapping from summary data with the "Sum of Single Effects" model. *PLoS Genet.* **18**, e1010299 (2022).

59. M. Togninalli, Ü. Seren, D. Meng, J. Fitz, M. Nordborg, D. Weigel, K. Borgwardt, A. Korte, D. G. Grimm, The AraGWAS Catalog: a curated and standardized Arabidopsis thaliana GWAS catalog. *Nucleic Acids Res.* **46**, D1150–D1156 (2018).

60. H. Wickham, M. Averick, J. Bryan, W. Chang, L. McGowan, R. François, G. Grolemund, A. Hayes, L. Henry, J. Hester, M. Kuhn, T. Pedersen, E. Miller, S. Bache, K. Müller, J. Ooms, D. Robinson, D. Seidel, V. Spinu, K. Takahashi, D. Vaughan, C. Wilke, K. Woo, H. Yutani, Welcome to the tidyverse. *J. Open Source Softw.* **4**, 1686 (2019).

61. P. Danecek, A. Auton, G. Abecasis, C. A. Albers, E. Banks, M. A. DePristo, R. E. Handsaker, G. Lunter, G. T. Marth, S. T. Sherry, G. McVean, R. Durbin, 1000 Genomes Project Analysis Group, The variant call format and VCFtools. *Bioinformatics* **27**, 2156–2158 (2011).

62. B. Langmead, S. L. Salzberg, Fast gapped-read alignment with Bowtie 2. *Nat. Methods* **9**, 357–359 (2012).

63. Y. Zhang, T. Liu, C. A. Meyer, J. Eeckhoute, D. S. Johnson, B. E. Bernstein, C. Nusbaum, R. M. Myers, M. Brown, W. Li, X. S. Liu, Model-based analysis of ChIP-Seq (MACS). *Genome Biol.* **9**, R137 (2008).

64. T. Z. Berardini, L. Reiser, D. Li, Y. Mezheritsky, R. Muller, E. Strait, E. Huala, The Arabidopsis information resource: Making and mining the “gold standard” annotated reference plant genome. *Genesis* **53**, 474–485 (2015).

65. Applied Research Applied Research Press, *Ultrafast and Memory-Efficient Alignment of Short DNA Sequences to the Human Genome* (CreateSpace Independent Publishing Platform, 2015).

66. T. Kivioja, A. Vähärautio, K. Karlsson, M. Bonke, M. Enge, S. Linnarsson, J. Taipale, Counting absolute numbers of molecules using unique molecular identifiers. *Nat. Methods* **9**, 72–74 (2011).

67. K. Toufighi, S. M. Brady, R. Austin, E. Ly, N. J. Provart, The Botany Array Resource: e-Northerns, Expression Angling, and promoter analyses. *Plant J.* **43**, 153–163 (2005).

68. M. A. Schon, M. D. Nodine, Widespread Contamination of Arabidopsis Embryo and Endosperm Transcriptome Data Sets. *Plant Cell* **29**, 608–617 (2017).

69. F. Hofmann, M. A. Schon, M. D. Nodine, The embryonic transcriptome of Arabidopsis thaliana. *Plant Reprod.* **32**, 77–91 (2019).

70. A. Schneider, D. Aghamirzaie, H. Elmarakeby, A. N. Poudel, A. J. Koo, L. S. Heath, R. Grene, E. Collakova, Potential targets of VIVIPAROUS1/ABI3-LIKE1 (VAL1) repression in developing Arabidopsis thaliana embryos. *Plant J.* **85**, 305–319 (2016).

71. M. D. Nodine, D. P. Bartel, Maternal and paternal genomes contribute equally to the transcriptome of early plant embryos. *Nature* **482**, 94–97 (2012).

72. Y. Hao, S. Hao, E. Andersen-Nissen, W. M. Mauck 3rd, S. Zheng, A. Butler, M. J.

Lee, A. J. Wilk, C. Darby, M. Zager, P. Hoffman, M. Stoeckius, E. Papalexi, E. P. Mimitou, J. Jain, A. Srivastava, T. Stuart, L. M. Fleming, B. Yeung, A. J. Rogers, J. M. McElrath, C. A. Blish, R. Gottardo, P. Smibert, R. Satija, Integrated analysis of multimodal single-cell data. *Cell* **184**, 3573–3587.e29 (2021).

73. T. A. Lee, J. Bailey-Serres, Integrative Analysis from the Epigenome to Translatome Uncovers Patterns of Dominant Nuclear Regulation during Transient Stress. *Plant Cell* **31**, 2573–2595 (2019).

74. P. S. Hosmani, M. Flores-Gonzalez, H. van de Geest, F. Maumus, L. V. Bakker, E. Schijlen, J. van Haarst, J. Cordewener, G. Sanchez-Perez, S. Peters, Z. Fei, J. J. Giovannoni, L. A. Mueller, S. Saha, An improved de novo assembly and annotation of the tomato reference genome using single-molecule sequencing, Hi-C proximity ligation and optical maps, *bioRxiv* (2019)p. 767764.

75. S. Ouyang, W. Zhu, J. Hamilton, H. Lin, M. Campbell, K. Childs, F. Thibaud-Nissen, R. L. Malek, Y. Lee, L. Zheng, J. Orvis, B. Haas, J. Wortman, C. R. Buell, The TIGR Rice Genome Annotation Resource: improvements and new features. *Nucleic Acids Res.* **35**, D883–7 (2007).

76. M. B. Hufford, A. S. Seetharam, M. R. Woodhouse, K. M. Chougule, S. Ou, J. Liu, W. A. Ricci, T. Guo, A. Olson, Y. Qiu, R. Della Coletta, S. Tittes, A. I. Hudson, A. P. Marand, S. Wei, Z. Lu, B. Wang, M. K. Tello-Ruiz, R. D. Piri, N. Wang, D. W. Kim, Y. Zeng, C. H. O'Connor, X. Li, A. M. Gilbert, E. Baggs, K. V. Krasileva, J. L. Portwood 2nd, E. K. S. Cannon, C. M. Andorf, N. Manchanda, S. J. Snodgrass, D. E. Hufnagel, Q. Jiang, S. Pedersen, M. L. Syring, D. A. Kudrna, V. Llaca, K. Fengler, R. J. Schmitz, J. Ross-Ibarra, J. Yu, J. I. Gent, C. N. Hirsch, D. Ware, R. K. Dawe, De novo assembly, annotation, and comparative analysis of 26 diverse maize genomes. *Science* **373**, 655–662 (2021).

77. S. Niu, J. Li, W. Bo, W. Yang, A. Zuccolo, S. Giacomello, X. Chen, F. Han, J. Yang, Y. Song, Y. Nie, B. Zhou, P. Wang, Q. Zuo, H. Zhang, J. Ma, J. Wang, L. Wang, Q. Zhu, H. Zhao, Z. Liu, X. Zhang, T. Liu, S. Pei, Z. Li, Y. Hu, Y. Yang, W. Li, Y. Zan, L. Zhou, J. Lin, T. Yuan, W. Li, Y. Li, H. Wei, H. X. Wu, The Chinese pine genome and methylome unveil key features of conifer evolution. *Cell* **185**, 204–217.e14 (2022).

78. D. B. Marchant, G. Chen, S. Cai, F. Chen, P. Schafran, J. Jenkins, S. Shu, C. Plott, J. Webber, J. T. Lovell, G. He, L. Sandor, M. Williams, S. Rajasekar, A. Healey, K. Barry, Y. Zhang, E. Sessa, R. R. Dhakal, P. G. Wolf, A. Harkess, F.-W. Li, C. Rössner, A. Becker, L. Gramzow, D. Xue, Y. Wu, T. Tong, Y. Wang, F. Dai, S. Hua, H. Wang, S. Xu, F. Xu, H. Duan, G. Theißen, M. R. McKain, Z. Li, M. T. W. McKibben, M. S. Barker, R. J. Schmitz, D. W. Stevenson, C. Zumajo-Cardona, B. A. Ambrose, J. H. Leebens-Mack, J. Grimwood, J. Schmutz, P. S. Soltis, D. E. Soltis, Z.-H. Chen, Dynamic genome evolution in a model fern. *Nat Plants* **8**, 1038–1051 (2022).

79. J. A. Banks, T. Nishiyama, M. Hasebe, J. L. Bowman, M. Gribskov, C. dePamphilis, V. A. Albert, N. Aono, T. Aoyama, B. A. Ambrose, N. W. Ashton, M. J.

Axtell, E. Barker, M. S. Barker, J. L. Bennetzen, N. D. Bonawitz, C. Chapple, C. Cheng, L. G. G. Correa, M. Dacre, J. DeBarry, I. Dreyer, M. Elias, E. M. Engstrom, M. Estelle, L. Feng, C. Finet, S. K. Floyd, W. B. Frommer, T. Fujita, L. Gramzow, M. Gutensohn, J. Harholt, M. Hattori, A. Heyl, T. Hirai, Y. Hiwatashi, M. Ishikawa, M. Iwata, K. G. Karol, B. Koehler, U. Kolukisaoglu, M. Kubo, T. Kurata, S. Lalonde, K. Li, Y. Li, A. Litt, E. Lyons, G. Manning, T. Maruyama, T. P. Michael, K. Mikami, S. Miyazaki, S.-I. Morinaga, T. Murata, B. Mueller-Roeber, D. R. Nelson, M. Obara, Y. Oguri, R. G. Olmstead, N. Onodera, B. L. Petersen, B. Pils, M. Prigge, S. A. Rensing, D. M. Riaño-Pachón, A. W. Roberts, Y. Sato, H. V. Scheller, B. Schulz, C. Schulz, E. V. Shakirov, N. Shibagaki, N. Shinohara, D. E. Shippen, I. Sørensen, R. Sotooka, N. Sugimoto, M. Sugita, N. Sumikawa, M. Tanurdzic, G. Theissen, P. Ulvskov, S. Wakazuki, J.-K. Weng, W. W. G. T. Willats, D. Wipf, P. G. Wolf, L. Yang, A. D. Zimmer, Q. Zhu, T. Mitros, U. Hellsten, D. Loqué, R. Otillar, A. Salamov, J. Schmutz, H. Shapiro, E. Lindquist, S. Lucas, D. Rokhsar, I. V. Grigoriev, The Selaginella genome identifies genetic changes associated with the evolution of vascular plants. *Science* **332**, 960–963 (2011).

80. J. L. Bowman, T. Kohchi, K. T. Yamato, J. Jenkins, S. Shu, K. Ishizaki, S. Yamaoka, R. Nishihama, Y. Nakamura, F. Berger, C. Adam, S. S. Aki, F. Althoff, T. Araki, M. A. Arteaga-Vazquez, S. Balasubrmanian, K. Barry, D. Bauer, C. R. Boehm, L. Briginshaw, J. Caballero-Perez, B. Catarino, F. Chen, S. Chiyoda, M. Chovatia, K. M. Davies, M. Delmans, T. Demura, T. Dierschke, L. Dolan, A. E. Dorantes-Acosta, D. M. Eklund, S. N. Florent, E. Flores-Sandoval, A. Fujiyama, H. Fukuzawa, B. Galik, D. Grimanelli, J. Grimwood, U. Grossniklaus, T. Hamada, J. Haseloff, A. J. Hetherington, A. Higo, Y. Hirakawa, H. N. Hundley, Y. Ikeda, K. Inoue, S.-I. Inoue, S. Ishida, Q. Jia, M. Kakita, T. Kanazawa, Y. Kawai, T. Kawashima, M. Kennedy, K. Kinose, T. Kinoshita, Y. Kohara, E. Koide, K. Komatsu, S. Kopischke, M. Kubo, J. Kyozuka, U. Lagercrantz, S.-S. Lin, E. Lindquist, A. M. Lipzen, C.-W. Lu, E. De Luna, R. A. Martienssen, N. Minamino, M. Mizutani, M. Mizutani, N. Mochizuki, I. Monte, R. Mosher, H. Nagasaki, H. Nakagami, S. Naramoto, K. Nishitani, M. Ohtani, T. Okamoto, M. Okumura, J. Phillips, B. Pollak, A. Reinders, M. Rövekamp, R. Sano, S. Sawa, M. W. Schmid, M. Shirakawa, R. Solano, A. Spunde, N. Suetsugu, S. Sugano, A. Sugiyama, R. Sun, Y. Suzuki, M. Takenaka, D. Takezawa, H. Tomogane, M. Tsuzuki, T. Ueda, M. Umeda, J. M. Ward, Y. Watanabe, K. Yazaki, R. Yokoyama, Y. Yoshitake, I. Yotsui, S. Zachgo, J. Schmutz, Insights into Land Plant Evolution Garnered from the Marchantia polymorpha Genome. *Cell* **171**, 287–304.e15 (2017).

81. D. Lang, K. K. Ullrich, F. Murat, J. Fuchs, J. Jenkins, F. B. Haas, M. Piednoel, H. Gundlach, M. Van Bel, R. Meyberg, C. Vives, J. Morata, A. Symeonidi, M. Hiss, W. Muchero, Y. Kamisugi, O. Saleh, G. Blanc, E. L. Decker, N. van Gessel, J. Grimwood, R. D. Hayes, S. W. Graham, L. E. Gunter, S. F. McDaniel, S. N. W. Hoernstein, A. Larsson, F.-W. Li, P.-F. Perroud, J. Phillips, P. Ranjan, D. S. Rokshar, C. J. Rothfels, L. Schneider, S. Shu, D. W. Stevenson, F. Thümmler, M. Tillich, J. C. Villarreal Aguilar, T. Widiez, G. K.-S. Wong, A. Wymore, Y. Zhang, A. D. Zimmer, R. S. Quatrano, K. F. X. Mayer, D. Goodstein, J. M. Casacuberta, K. Vandepoele, R. Reski, A. C. Cuming, G. A. Tuskan, F. Maumus, J. Salse, J. Schmutz, S. A.

Rensing, The Physcomitrella patens chromosome-scale assembly reveals moss genome structure and evolution. *Plant J.* **93**, 515–533 (2018).

82. P. A. Ewels, A. Peltzer, S. Fillinger, H. Patel, J. Alneberg, A. Wilm, M. U. Garcia, P. Di Tommaso, S. Nahnsen, The nf-core framework for community-curated bioinformatics pipelines. *Nat. Biotechnol.* **38**, 276–278 (2020).

83. P. Di Tommaso, M. Chatzou, E. W. Floden, P. P. Barja, E. Palumbo, C. Notredame, Nextflow enables reproducible computational workflows. *Nat. Biotechnol.* **35**, 316–319 (2017).

84. S. Zhang, M. Xu, Z. Qiu, K. Wang, Y. Du, L. Gu, X. Cui, Spatiotemporal transcriptome provides insights into early fruit development of tomato (Solanum lycopersicum). *Sci. Rep.* **6**, 23173 (2016).

85. S. C. Stelpflug, R. S. Sekhon, B. Vaillancourt, C. N. Hirsch, C. R. Buell, N. de Leon, S. M. Kaeppler, An Expanded Maize Gene Expression Atlas based on RNA Sequencing and its Use to Explore Root Development. *Plant Genome* **9** (2016).

86. L. Xia, D. Zou, J. Sang, X. Xu, H. Yin, M. Li, S. Wu, S. Hu, L. Hao, Z. Zhang, Rice Expression Database (RED): An integrated RNA-Seq-derived gene expression database for rice. *J. Genet. Genomics* **44**, 235–241 (2017).

87. P.-F. Perroud, F. B. Haas, M. Hiss, K. K. Ullrich, A. Alboresi, M. Amirebrahimi, K. Barry, R. Bassi, S. Bonhomme, H. Chen, J. C. Coates, T. Fujita, A. Guyon-Debast, D. Lang, J. Lin, A. Lipzen, F. Nogué, M. J. Oliver, I. Ponce de León, R. S. Quatrano, C. Rameau, B. Reiss, R. Reski, M. Ricca, Y. Saidi, N. Sun, P. Szövényi, A. Sreedasyam, J. Grimwood, G. Stacey, J. Schmutz, S. A. Rensing, The Physcomitrella patens gene atlas project: large-scale RNA-seq based expression data. *Plant J.* **95**, 168–182 (2018).

88. Y.-L. Xiao, G.-S. Li, Differential expression and co-localization of transcription factors during the indirect de novo shoot organogenesis in the fern Ceratopteris richardii, *Research Square* (2023). https://doi.org/10.21203/rs.3.rs-2531906/v1.

89. G. Pertea, M. Pertea, GFF utilities: GffRead and GffCompare. *F1000Res.* **9**, 304 (2020).

90. R. Patro, G. Duggal, M. I. Love, R. A. Irizarry, C. Kingsford, Salmon provides fast and bias-aware quantification of transcript expression. *Nat. Methods* **14**, 417–419 (2017).

91. L. Huang, J. Schiefelbein, Conserved Gene Expression Programs in Developing Roots from Diverse Plants. *Plant Cell* **27**, 2119–2132 (2015).

92. N. Sharma, P. L. Bhalla, M. B. Singh, Transcriptome-wide profiling and expression analysis of transcription factor families in a liverwort, Marchantia polymorpha. *BMC Genomics* **14**, 915 (2013).

93. B. Bushnell, “BBMap: A Fast, Accurate, Splice-Aware Aligner” (LBNL-7065E, Lawrence Berkeley National Lab. (LBNL), Berkeley, CA (United States), 2014); https://www.osti.gov/biblio/1241166.
